# Expertly validated models suggest responses to climate change are related to species traits: a phylogenetically-controlled analysis of the Order Lagomorpha

**DOI:** 10.1101/001826

**Authors:** Katie Leach, Ruth Kelly, Alison Cameron, W. Ian Montgomery, Neil Reid

## Abstract

Climate change during the last five decades has impacted significantly on natural ecosystems and the rate of current climate change is of great concern among conservation biologists. Species Distribution Models (SDMs) have been used widely to project changes in species’ bioclimatic envelopes under future climate scenarios. Here, we aimed to advance this technique by assessing future changes in the bioclimatic envelopes of an entire mammalian Order, the Lagomorpha, using a novel framework for model validation based jointly on subjective expert evaluation and objective model evaluation statistics. SDMs were built using climatic, topographical and habitat variables for all 87 species under past and current climate scenarios. Expert evaluation and Kappa values were used to validate past and current distribution models and only those deemed ‘modellable’ through our framework were projected under future climate scenarios (58 species). We then used phylogenetically-controlled regressions to test whether species traits were correlated with predicted responses to climate change. Climate change will impact more than two-thirds of the Lagomorpha, with leporids (rabbits, hares and jackrabbits) likely to undertake poleward shifts with little overall change in range extent, whilst pikas are likely to show extreme shifts to higher altitudes associated with marked range declines, including the likely extinction of Kozlov’s Pika (*Ochotona koslowi*). Smaller-bodied species were more likely to exhibit range contractions and elevational increases, but showing little poleward movement, and fecund species were more likely to shift latitudinally and elevationally. Our results suggest that species traits may be important indicators of future climate change and we believe multi-species approaches, as demonstrated here, are likely to lead to more effective mitigation measures and conservation management.

## Introduction

Changes in climate are predicted to have strong influences on the ecology and distribution of species [1], [2], with pronounced impacts on terrestrial biodiversity [3]. Although the climate changes naturally (and usually slowly), the rate of recent anthropogenically-induced change [4] is causing concern amongst conservation biologists [5]. Future climate change may have large effects on species niches i.e. the biotic and abiotic conditions in which a species can persist [6]. Species are predicted to adapt their bioclimatic niche, migrate to maintain their current niche, or become range restricted and undergo population decline, local or global extinction under future scenarios [7].

The Lagomorpha are an important mammalian Order economically and scientifically as a major human food resource, model laboratory animals, valued game species, pests of agricultural significance and key elements in food chains providing scientific insights into entire trophic systems. Lagomorphs are native on all continents except Antarctica, occurring from sea level to >5,000m and from the equator to 80°N spanning a huge range of environmental conditions, and also include some very successful invasive species [8].

The taxonomy of the Lagomorpha in recent decades has been in a state of flux but all species belong to two families: the *Ochotonidae* and the *Leporidae*. The *Ochotonidae* consists of one monotypic group in the genus *Ochotona* containing 25 species of small, social, high-altitude pikas. The *Leporidae* has 32 species of large, solitary, cursorial hares and jackrabbits in a single genus *Lepus* and 30 species of medium-sized, semi-social, fossorial rabbits comprising ten genera [8]. A quarter of lagomorphs are listed in the IUCN Red List of Threatened Species (www.iucnredlist.org) with a notable number of highly range-restricted species including fourteen listed under the IUCN Criteria B, with an extent of occurrence estimated to be less than 20,000 km^2^. In addition, pikas as high-altitude specialists with very high body temperatures of 39.3 – 41.0°C [9] are extremely susceptible to changes in their environment, particularly ambient temperatures [10].

Species Distribution Models (SDMs) are widely used in ecology and relate species occurrences at known locations to environmental variables to produce models of environmental suitability, which can be spatially or temporally extrapolated to unsurveyed areas and into past or future conditions (e.g. [11]). Although SDMs have been highly influential in the field of ecology, their limitations have been widely reviewed (e.g. [12]). The impact of climate change on species distributions has been modelled in a wide range of studies and a number of responses have been described (e.g. [1], [5], [13]). Mammalian distributional changes have been well studied over the past decade and indicate that future climate change will have profound impacts (e.g. [5], [14], [15], [16]). Mammals in the Western hemisphere are unlikely to keep pace with climate change, with 87% expected to undergo range contractions [16], and mammals in Mediterranean regions, particularly endemic species, are predicted to be severely threatened by future climate change [15]; shrews are especially vulnerable to future changes [5], [15]. These distributional responses have also been noted in studies of past climatic changes, for example, Moritz *et al*. [14] found that from the early 20^th^ century to the present, small mammals in a North American national park substantially shifted their elevational range upwards corresponding to ~3°C increase in minimum temperatures.

The predicted impact of climate change on species distributions has only rarely been linked with species traits. Yet, species traits are widely accepted as potentially important indicators of responses to climate change and identifying such traits may be crucial for future conservation planning (e.g. [14], [17], [18]). Traits that directly impact climatic conditions experienced by a species, for example their activity cycle, are likely to be more important in mediating species responses to projected climate change than traits such as diet breadth. If species can broaden their occupied bioclimatic niche through trait plasticity, for example, altering their diel patterns of activity, then they may be less susceptible to future change [19]. Mammalian species active during certain times of the day will experience a limited range of climatic conditions, whereas more flexible species can select the conditions in which they are active [20], and may, therefore, be less susceptible to future change [19]. Small body size, nocturnal behaviour and burrowing may have allowed mammalian species to ‘shelter’ from climatic changes during the 66M year duration of the Cenozoic era [21]. Burrowing rabbits may be able to adapt to climate change by seeking underground refugia from extreme or fluctuating temperatures [22], whilst larger cursorial species, such as the hares and jackrabbits which, in the majority of cases, live above ground may have less variability in microclimate opportunities within which to shelter [23]. Thus, species with narrow environmental tolerances, poor dispersal ability and specialised habitats may be more susceptible to climate change [24]. Dispersal is also likely to be very important in future species distributions, with larger species being more mobile and, therefore, potentially better prepared to track climatic changes [16],

Past studies have modelled the response of large numbers of species to predicted climate change [3] or dealt with a few key species at Order-level [16], lagomorphs due to their restricted diversity provide an opportunity to rigorously examine the response of every species yielding a detailed picture of change within an entire Order for the first time. Crucially, the small number of lagomorph species, compared to other mammalian groups, means that datasets can be verified in detail, modelled individually and outputs expertly validated. Moreover, lagomorphs have a nearly global terrestrial distribution and occupy a wide range of biomes providing an opportunity to examine the response of similar species from tundra to desert and islands to mountain summits.

Here, we assess the projected change in the bioclimatic envelopes of all ‘modellable’ lagomorph species under future climate change using a framework for model validation based jointly on subjective expert evaluation cross-validated with objective model evaluation statistics. We predict lagomorph species distributions will increase in elevation and poleward movement under future climate change, but with significant differences between pikas, rabbits, hares and jackrabbits due to dissimilarities in species traits, for example body size. Lagomorph morphological and life history traits will be correlated with the predicted responses to future climate change in order to test this hypothesis. We theorise that flexibility in activity cycle and larger body sizes, which may lead to greater mobility, will result in species being less vulnerable to future climatic changes and better able to track climate niches.

## Materials and Methods

### Species Data

A total of 139,686 records including all 87 lagomorph species were either downloaded from the Global Biodiversity Information Facility (GBIF) Data Portal (data.gbif.org), collated from species experts or members of the IUCN Species Survival Commission (SSC) Lagomorph Specialist Group (LSG) and/or extracted from the literature for data deficient species as advised by experts. All past and current occurrence records, sorted by species, can be viewed on http://lagomorphclimatechange.wordpress.com/. Taxonomic accuracy was dealt with by checking all records against the latest IUCN taxonomy; if names did not match after cross-referencing with taxonomic synonyms and previous names they were rejected. Spatial data accuracy was dealt with by removing any obviously erroneous records for the target species if they fell outside the extent of the IUCN geographic range polygon. In addition, occurrences recorded with a spatial resolution of >2km were removed and duplicate records were eliminated unless they were recorded in different temporal periods (pre-1950 and post-1950). This left 41,874 records of which 3,207 were pre-1950 and 38,667 were post-1950. In this study, spatial and temporal bias in sampling was eliminated by only selecting the background data (a random sample of the environmental layers describing pseudo-absences), 10,000 points, from sites at which any lagomorph species had been recorded and ensuring there were the same proportion of pre-1950 to post-1950 background points as there were pre-1950 to post-1950 species records.

### Environmental Parameterisation

Empirical climate data for past, current and future scenarios were downloaded from the R-GIS Data Portal (http://r-gis.org/), WorldClim (http://www.worldclim.org/) and CCAFS GCM Data Portal (http://www.ccafs-climate.org/data/) respectively at 30 arc-second resolution (≈1km grid cells). Records from pre-1950 were associated with mean climate data from 1900–1949 and those post-1950 were associated with mean data from 1950-2000. Projected future climate data were obtained from the IPCC 4^th^ Assessment Report using the A2 emissions scenario. Data from the A2 future climate change scenario was used because although it was originally described as “extreme climate change” it now appears to best represent the trend in observed climate. Eight environmental variables were used in their raw format and seven composite variables were calculated (see Appendix 1 in Supporting Information).

### Species Distribution Modelling

SDMs were run using MaxEnt version 3.3.3k [11], [25]. Models were built using two different sets of input data: i) pre-and post–1950 data, or ii) post-1950 data only. The ‘samples with data’ (SWD) input format in MaxEnt was used for data entry, pairing pre–1950 records with mean climatic variables from 1900–1949 and post-1950 records with mean data from 1950–2000. The models were validated using either 10 replicate bootstrapping for species with low numbers of records (<30) or 4-fold cross-validation for species with high numbers of records (≥30). More than 30 records were needed for 4 fold cross-validation as this equates to ~8 points per replicate, which has been deemed adequate for modelling some species distributions [26], bootstrapping was used with <30 records as this maximised the points used for both training and testing. Ten bootstrap replicates were needed to minimise computational power and maximise accuracy. Linear, quadratic and product feature types were used. The 10 percentile threshold was applied to define likely presence and absence of each species.

Records for each species were associated with global land cover data downloaded from the ESA GlobCover 2009 Project (http://due.esrin.esa.int/globcover/) and any land class not occupied by the target species was marked as unsuitable. Outputs were then clipped to a minimum convex polygon of the species occurrence records, buffered by a dispersal value specific to each species which was verified by experts from the IUCN Species Survival Commission Lagomorph Specialist Group, to remove areas of over-prediction. This meant that the suitable area within reach of each species was predicted, and led to more conservative outputs than just simply predicting suitable area. Although this assumes occurrence records are complete, this was a reasonable way to correct for potential biogeographic overprediction and a similar method was advocated by Kremen *et al*. [27]. Annual dispersal distances were elicited from each species expert during the model evaluation procedure (see Appendix 2 in Supporting Information). Dispersal distances were either observed by the lagomorph experts (personal observations), documented in the literature, or for species where no data were available, the average distance was calculated from the specific group to which it belonged (i.e. Asian pikas or African hares).

Non-native ranges for the only two invasive lagomorphs, European hare (*Lepus europaeus*) and rabbit (*Oryctolagus cuniculus*), were not modelled because invasive species are not at equilibrium with the environment and their niches cannot be transferred in space and time [28]. Mountain hare (*Lepus timidus*) populations in Ireland and mainland Eurasia were modelled separately due to the distinct morphological, phenotypic, behavioural, ecological and genetic differences between the Irish sub-species (*L. t. hibernicus*) and other mountain hares (e.g. [29]), but the outputs for each geographic region were subsequently combined to produce a single model reflecting the current classification of the Irish hare as an endemic sub-species.

### Model Evaluation

A bespoke website (lagomorphclimatechange.wordpress.com) was created to allow each species expert to review the output of their allocated species. Expert evaluation, whereby an acknowledged expert on each species judges model predictions for current and past distributions, can be a useful tool prior to making future extrapolations [30]. A framework combining expert evaluation with reliable model evaluation metrics allows species distribution models to be assessed before they are used in future projections to infer likely future changes in distribution.

Forty-six lagomorph experts, including 20 members of the Lagomorph Specialist Group (LSG) of the IUCN Species Survival Commission (SSC) and 26 recognised lagomorph researchers selected based on recent publications, were paired to species (see Table S1 in Supporting Information) and asked to assess whether model projections accurately, roughly or did not capture the current and past range of each species i.e. good, medium or poor respectively according to the criteria in Anderson *et al*. [30]. Experts were asked to select the most accurate representation of the current and past range from the following models: i) pre– and post–1950 input data showing the suitable bioclimatic envelope, ii) pre– and post-1950 input data showing the suitable bioclimatic envelope restricted to suitable habitat, iii) post– 1950 input data only showing the suitable bioclimatic envelope, or iv) post–1950 input data only showing the suitable bioclimatic envelope restricted to suitable habitat.

Independent model evaluation in SDM studies often uses the Area Under the Curve (AUC) value but this has been heavily criticised [31]. AUC is not advocated for model evaluation because Receiver Operating Characteristic (ROC) curves cannot be built for presence/absence or presence/pseudo-absence data [32] and AUC values can be influenced by the extent of model predictions [31]. There are also known limitations with using alternative metrics [32] such as sensitivity (proportion of presences which are correctly predicted), specificity (proportion of absences which are correctly predicted) or True Skill Statistic (a prevalence independent model metric calculated using sensitivity and specificity). Misleading commission errors, which can arise from species not being at equilibrium with the environment, can affect such metrics. On the other hand, Kappa is an objective measure of prediction accuracy based on input species records and background points adjusted for the proportion of correct predictions expected by random chance [33]. Kappa utilises commission and omission errors [34], and although it does not take into account prevalence like the True Skill Statistic [32] and can sometimes produce misleading commission errors [35], it has widely accepted thresholds which can be useful in model evaluation [36], [37], [38]. There are documented flaws with all possible model evaluation statistics, however Kappa was chosen due to the reasons above and its common use in model evaluation (e.g. [39]), with the main focus of evaluation in this study being the expert validation approach.

The ‘accuracy’ function in the SDMTools package [40] in R (version 3.0.2) was used to calculate the Area under the Curve (AUC) value, omission rate, sensitivity, specificity, proportion correct, Kappa and True Skill Statistic for completeness, but only Kappa was taken forward for use in the model validation framework. The optimum threshold was taken as Kappa >0.4 as this value has been advocated in a range of previous studies [32], [36], [38]. Models that had a Kappa >0.4 and those that were ranked as either good or medium by expert evaluation were defined as “modellable” because they demonstrate good model fit and predictive ability; these species were carried forward for projection and further analysis. Those models with a Kappa <0.4 or ranked as poor by expert evaluation were defined as “unmodellable”, with poor model fit and predictive ability, and were rejected from further analysis (see Figure S1). Hereafter, species are referred to as “modellable” or “unmodellable” as explicitly defined above.

### Future Predictions

The model settings, for example, the input data used (pre-or post-1950) or restriction/ no restriction to occupied land classes, which provided the optimal outcome i.e. the highest Kappa value and best expert evaluation were used to project modellable species bioclimatic envelopes under future climate during the 2020s, 2050s and 2080s. Future predictions were clipped to the buffered minimum convex polygon of the target species further buffered by the dispersal distance (kilometres/year) of each species multiplied by the number of years elapsed from the present (1950-2000) taken as 1975. Predicted range size, mean latitude and minimum, mean and maximum elevation for each species and each time period were calculated. Model outputs for each species can be found in Appendix 2; unmodellable species projections are included only for reference.

### Species Traits

Species trait data were downloaded from the PanTHERIA database [41] and updated by searching the literature. Correlated traits were removed to reduce multicollinearity (i.e. tolerance <2 and VIF >5) and the final set of traits used to describe each modellable species were: activity cycle (nocturnal only, diurnal only, flexible), adult body mass (g), diet breadth (number of dietary categories eaten from: vertebrates, invertebrates, fruit, flowers/nectar/pollen, leaves/branches/bark, seeds, grass and roots/tubers), gestation length (days), habitat breadth (above ground-dwelling, aquatic, fossorial and ground-dwelling), home range (km^2^), litters per year, litter size, population density (*n*/km^2^) and age at sexual maturity (days).

### Statistical Analysis

To illustrate projected changes in the distribution of lagomorph species, the difference in predicted species richness per cell was calculated between the 1930s (1900–1949) and 2080s (2070–2099). The difference in model output metrics (range size, mean latitude and minimum, mean and maximum elevation) was calculated between the 1930s and 2080s. Change in range size was expressed in percentage change but change in latitude was represented as degree movement towards the poles and change in elevation in metres. Generalised Least Square (GLS) models, with an autoregressive moving average (ARMA) correlation structure, in the ‘nlme’ package in R version 3.0.2 [42] were used to test the differences in temporal trends for range change, poleward movement and elevation between lagomorph groups: 1) pikas, 2) rabbits and 3) hares and jackrabbits. ARMA was used to explicitly account for the non-independent nature of the time-series periods. Phylogenetic Generalized Least-Squares (PGLS) regressions were performed to test whether changes in model predictions varied with morphological or life history traits. PGLS analysis was run using the ‘caper’ package in R [43]. A lagomorph phylogeny was extracted from the mammalian supertree provided by Fritz *et al*. [44]. Likely clade membership for five species not included in this phylogeny was determined from Ge *et al*. [45], and then missing tips were grafted on using an expanded tree approach [46]. Outliers were removed prior to analysis, because species with large residuals may overly influence the results of the regression, and they were identified as those with a studentized residual >3 units following Cooper *et al*. [47]. All subsequent models exhibited normally distributed residuals tested using Shapiro-Wilk. The scaling parameter lambda varies from 0, where traits are independent of phylogeny, to 1 where species’ traits covary in direct proportion to their shared evolutionary history [48]. We estimated lambda for each model and tested whether it was significantly different from 0 or 1 during the PGLS analysis. All subset regressions were run using the ‘dredge’ function in the ‘MuMIn’ [49] package in R, using AICc as the rank estimate, and then model averaging was used to describe the effect of each variable.

## Results

There was a high degree of agreement between expert model classification and Kappa values, however experts were often more critical because, for example, they classified 25 species as ‘poor’ but these species had a mean Kappa of ~0.6 (Fig. 1). Fifty-eight species (67%) were deemed modellable with expert validation classed as medium or good and Kappa >0.4 and 29 species (33%) rejected as unmodellable with expert validation classed as poor and/or Kappa <0.4. Unmodellable species were 4 times more likely to be listed by the IUCN as Data Deficient than modellable species, with 8 unmodellable species (28%) listed as threatened. The majority of species with small sample sizes were classed as unmodellable; the median number of occurrence points for modellable species was 36 and for unmodellable 13. Hereafter, all results are for modellable species only and, therefore, are a highly conservative estimate of the impact of climate change on the Order.

Global changes in predicted lagomorph species richness suggest that almost a third of the Earth’s terrestrial surface (31.5 million km^2^) is predicted to experience loss of lagomorph species by the 2080s (Fig. 2). The desert regions of North-eastern China and hills of Sichuan become increasingly unsuitable, potentially losing up to 10 species by 2080, including the woolly hare (*Lepus oiostolus*) and Glover’s pika (*Ochotona gloveri*) which are predicted to undergo dramatic movements to higher elevations. In contrast, predicted species gains were notable across: (a) northern Eurasia, due to poleward movement of the mountain hare (*Lepus timidus);* and, (b) North America, where some regions e.g. the Upper Missouri catchment of Montana and North Dakota were predicted to gain up to 5 species. This includes the desert and eastern cottontails (*Sylvilagus audubonii* and *S. floridanus* respectively) which are predicted to exhibit strong poleward movements. The majority of African lagomorph species were classed as unmodellable and as such Fig. 2 is largely data deficient for the continent.

Thirty-six lagomorph species are predicted to experience range loss (63%), 48 poleward movements (83%) and 51 elevational increases (88%). Thirty-five species (60%) are predicted to undergo range declines *and* either poleward movements *or* elevational increases. On average, all three groups of lagomorphs exhibited significant poleward shifts (*F*_df=1,234_=13.5, *p*<0.001) and elevational increases (*F*_df=1,234_ = 44.2, *p*<0.001) by 2080 (Fig. 3, also see Table S2). The average poleward shift for the Order was 1.1° and elevational shift 165m. Pikas exhibited the most substantial mean increase in elevation becoming increasingly isolated on mountain summits (e.g. the Rockies in North America and the Tibetan Plateau and high Himalayas) resulting in a significant 31% range contraction (*F*_df = 1,234_ = 3.7, *p* = 0.03). In addition, lagomorph species occupying islands (*n*=6) will, on average, lose 8km^2^ of their ranges compared to 0.2km^2^ gain for continental species (*n* = 52), whilst mountain dwelling species (*n* = 24) will lose 37km^2^ of their ranges compared to 25km^2^ gain for lowland species (*n* = 34).

PGLS models indicate that members of each group were capable of showing a variety of responses i.e. species of pika, rabbits, hares or jackrabbits exhibited both increases and decreases in each response variable (Fig. 4). All traits used in the PGLS models are described in the methods and full results can be found in Table S3. Here we present only the significant results. There was a significant positive relationship between predicted range change and adult body mass (*β*=0.258, *F*_df=4,52_=2.308, *p*=0.021) with hares and jackrabbits generally increasing their range by 2080 and pikas exhibiting range contraction (Fig. 4; also see Table S3). Adult body mass was positively associated with predicted poleward movement (*β*=0.196, *F*_df=5,49_=1.989, *p*=0.047) and inversely related to predicted elevational change (*β*= −0.183, *F*_df=2,50_=2.019, *p*=0.043). The average adult body mass for lagomorph species living in mountainous regions was 836g, compared to 1.8kg in lowland areas. The number of litters a species produces per year was positively related to predicted latitudinal and maximum elevational shifts (*β*=0.215, *F*_df = 5,49_ = 2.731, *p*=0.006 and *β*=0.160, *F*_df=2,53_=1.746, *p*=0.081 respectively) with more fecund species being capable of more extreme upward or poleward shifts. There was also a positive relationship between species dietary niche and the degree of predicted poleward shift (*β*=0.181, *F*_df=5,49_=2.190, *p*=0.029). No significant relationships were found between activity cycle and predicted changes in species distribution.

## Discussion

The Lagomorpha as a whole are predicted to exhibit much greater poleward (mean ± 95%CI = 1.1° ± 0.5°) and elevational shifts (165m ± 73m) by 2100 than calculated in a recent metaanalysis collating information on a wide variety of taxonomic groups [1]. On average birds, butterflies and alpine herbs were predicted to shift ~0.8° poleward and increase in elevation by ~90m by 2100. Thuiller *et al*. [5] found that mammals were less vulnerable to change than other groups, but more than 50% of shrews could lose more than 30% of their suitable climatic space. In comparison, our study shows that only 34% of modellable lagomorphs will lose more than 30% of their suitable climatic space, but lagomorphs appear to show more notable changes in elevation and poleward movement. A study by Schloss *et al*. [16] found that, in some scenarios, 50% of mammal species in North and South America will be unable to keep pace with future climate change. Our results, therefore, indicate that lagomorphs follow similar trends to other mammalian climate change studies, but with more substantial poleward and elevational shifts. Furthermore, our results are conservative estimates due to the exclusion of unmodellable species, most notably African species. Parts of Africa are expected to become drier and warmer under future climate, with substantial increases in arid land [4] which will likely lead to negative consequences for lagomorphs not assessed here.

In contrast, a recent study of empirical climate studies by McCain & King [19] has shown that only about 50% of, mostly North American, mammal species respond to climate change by shifting in latitude and elevation. Results of four lagomorph studies are included which show that the pygmy rabbit (*Brachylagus idahoensis*) undergoes extirpation and contraction due to climatic changes [50], the range of the snowshoe hare (*Lepus americanus*) and collared pika (*Ochotona collaris*) does not change [51], [52] and the American pika (*Ochotona princeps*) mostly undergoes extirpation and upslope contraction, but some sites within the range show no change [14], [53]. Although we find that more than 50% of lagomorphs respond to climate change, our results are largely congruent with these empirical studies, indicating range contraction in the American pika and pygmy rabbit, and little change in the ranges of the snowshoe hare and collared pika.

Pikas are predicted to show elevational rather than latitudinal shifts because these high– altitude specialists are known to be extremely susceptible to increases in temperature or unpredictable seasonality, which could potentially lead to heat stress [9]. On the other hand leporids, typically being lowland species, exhibited less substantial increases in elevation but greater poleward shifts, for example, the mountain hare (*Lepus timidus*) which is predicted to shift poleward. This is probably due to the high sensitivity of its boreo-alpine niche to changes in temperature [54], [55]. Indeed, even the Irish hare (*L. t. hibernicus*) which inhabits temperate grasslands, unlike other mountain hares, is predicted to experience a contraction in the south-east of Ireland. As global temperatures increase, northern latitudes will become more climatically suitable for southern leporids and, therefore, species bioclimatic envelopes will track poleward to match. Rabbits, hares and jackrabbits were thus predicted to exhibit little overall change in the total extent of their ranges. The majority of the modellable rabbit species were from the genus *Sylvilagus* inhabiting south and eastern USA and Mexico; a region projected to become generally drier [56] inducing latitudinal shifts in species to track suitable habitats or vegetation. Thus, by shifting their ranges poleward, leporids are predicted to be able to maintain or increase their range extent suggesting they are considerably less sensitive to projected climate change than pikas.

PGLS models estimated lambda to be close to zero for changes in range, poleward movement and elevation indicating that observed trends were independent of phylogeny [48]. Smaller species, principally pikas and some rabbits, were typically more likely to make elevational shifts in range whilst larger species, principally hares and jackrabbits, had a greater tendency for poleward shifts. This relationship was also discovered in empirical climate studies (e.g. [19]) and is probably due to the large number of small–bodied species that dwell in mountainous regions (*n*=24) which have more opportunity to shift altitudinally. A 1m elevational shift is equivalent to a 6km latitudinal shift [1], so it is easier for smaller species to shift in elevation rather than latitudinally. This could be explained due to a relationship between adult body mass and dispersal distance, which was used to clip model projections, but a Spearman’s Rank correlation suggested no correlation between these two variables in our dataset (see Table S4).

Trait analyses also showed a significant positive relationship between fecundity and extreme elevational or poleward movement which is comparable to a study by Moritz *et al*. [14], who found that fecund small mammals in Yosemite National Park, USA, were more likely to expand their ranges upward than less fecund counterparts. We found no association between activity cycle and vulnerability to climate change in the Lagomorpha as hypothesized and reported in McCain & King [19], but this may be due to the varied nature of lagomorph activity. The activity of the European hare is known to be less consistent and partially diurnal in the summer [57] and the American pika also shows seasonal changes in activity patterns [58]. However, we do acknowledge the potential drawbacks of linking traits to modelled distributional changes rather than empirical-based studies, i.e. non-independence of trait results; nevertheless the analysis presented here enables understanding of the traits which could potentially lead to vulnerability to future climate change.

The uncertainties in SDM using projected future climate scenarios are well described [12]. Models are vulnerable to sampling bias [59], spatial scale issues [60], lack of data for rare species [26], uncertainty regarding future climate conditions [61] and insufficient independent evaluation [60]. We have tackled these explicitly by accounting for sampling bias by restricting the set of background points used, using data with the highest spatial resolution available (30 arc-second) and selecting species records to match, bootstrapping models for rare species with few records, averaging climatic data from five global circulation models and using a framework of joint expert and metric-driven model validation to segregate current distribution model outputs into those that were modellable and unmodellable before subsequent projection and analysis. Nevertheless, our predictions could still be potentially confounded by species-area relationships [62] and biological interactions [63].

Regardless of individual model outcomes (see Appendix 2), the overall trends observed across the Order Lagomorpha as a whole are compelling. This study did not take account of shifts in habitats, vegetation or human impact in response to climate change, but we have shown that adaptation to future climate conditions may be possible as some species were predicted to exhibit poleward movements, with only modest shifts in range or elevation e.g. eastern (*Sylvilagus floridanus*) and Appalachian cottontail (*Sylvilagus obscurus*).

The predicted changes in climate conditions are likely to have greater impacts on isolated lagomorph populations, i.e. those on islands and in high-altitude systems. If changes continue at the rate currently predicted until the 2080s, then there may be no climatically suitable range available for some montane or isolated species e.g. the Tres Marias cottontail (*Sylvilagus graysoni*) or black jackrabbit (*Lepus insularis*). Conservation strategies, such as assisted migration, could be the only option for these highly range-restricted species. Furthermore, conservation management will need to focus on small-bodied mammals as these are predicted to show more dramatic responses to changing climate. Small mammals are key in food webs sustaining predator populations, impacting plant communities by grazing and soil biology and hydrology by burrowing. Thus, fundamental shifts in lagomorphs globally may cause trophic cascade effects, especially in northern latitudes such as the cyclic systems of the Arctic.

The advancing knowledge of past extinction rates for the Lagomorpha [45], along with significantly better bioclimatic envelope modelling, could aid the prediction and prevention of future extinctions. Our models suggest that Kozlov’s pika (*Ochotona koslowi*) may become extinct by the 2080s as the elevational increases required to maintain its current bioclimatic envelope disappear as it reaches the maximum elevation available. We have shown here how expert validation can be effectively integrated into the model evaluation process in order to improve model predictions and advocate use of this framework in future SDM studies. Assessment of vulnerability to climate change is needed urgently to identify how and to what extent species, taxa, communities and ecosystems are susceptible to future changes, taking into account likely changes to vegetation, human pressures and interspecific interactions, and to direct conservation management in an efficient and effective manner. Multi-species approaches are likely to lead to more effective mitigation measures and contribute to our understanding of the general principles underpinning the biogeographical and ecological consequences of climate change impacts.

## Acknowledgments

We thank all lagomorph experts for validating models (see Table S1) and particular thanks to Michael Barbour, Kai Collins and Christy Bragg, Deyan Ge, David Hik, Dana Lee, Andrey Lissovsky, Andrew Smith, Franz Suchentrunk, Zelalem Tolesa and Weihe Yang for contributing data, taxonomic expertise and also validating models.

## Appendix S1. Supplementary Methods and References

## Appendix S2.

89 × 1 page species accounts (including: species information, predicted changes in distributions, plots of change in range, latitude and elevation, model evaluation metrics and variable response curves).

Table S1. Lagomorph experts, institutions and species evaluated.

Figure S1. Framework for assessing whether species were “modellable” or “unmodellable” based on Kappa values and expert evaluation classification.

Table S2. Results of Generalised Least Square models characterising predicted lagomorph bioclimatic envelope changes.

Table S3. Results of phylogenetically-controlled generalised least square regressions.

Table S4. Results of Spearman’s rank correlation.

## Figure Legends

**Figure 1.**
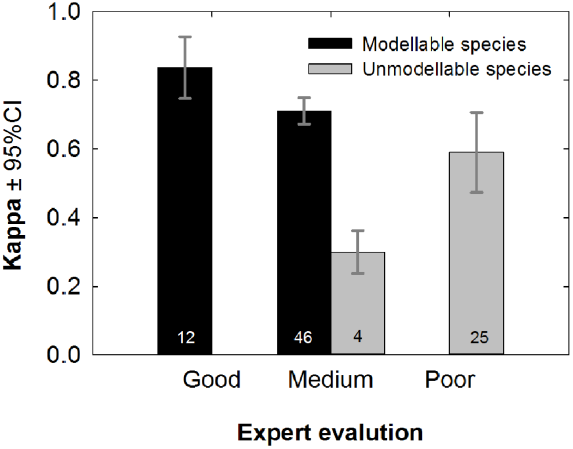
Agreement between expert evaluation and model accuracy. Mean Kappa ±95% confidence intervals within the categories assigned by expert evaluation. Black bars indicate species that were deemed “modellable” and retained for further analyses, whereas grey bars indicate “unmodellable species” that were rejected. Sample sizes (i.e. numbers of species) are shown in the bars.

**Figure 2.**
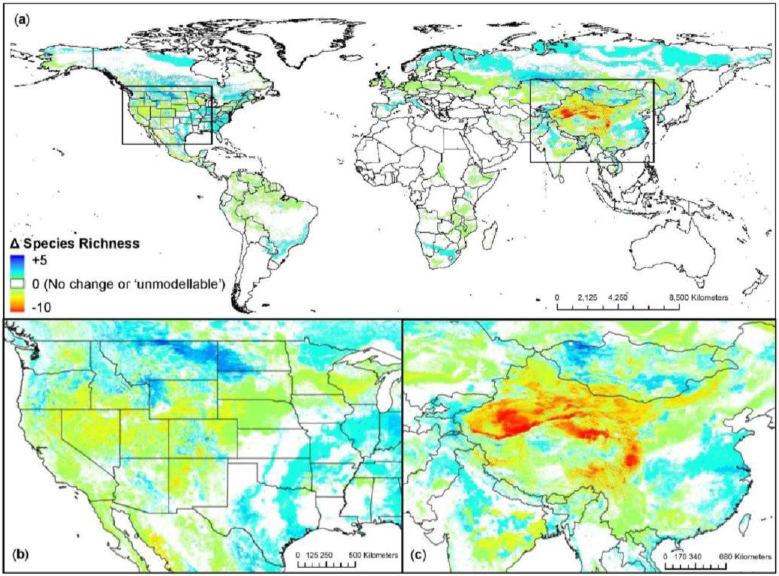
Change in predicted lagomorph species richness from the 1930s to 2080s. (**a**) Global patterns in predicted species loss and gain showing details in (**b**) North America and (**c**) Asia. White areas indicate no change in species richness or areas that are occupied by “unmodellable” species with uncertain outcomes (i.e. data deficient)

**Figure 3.**
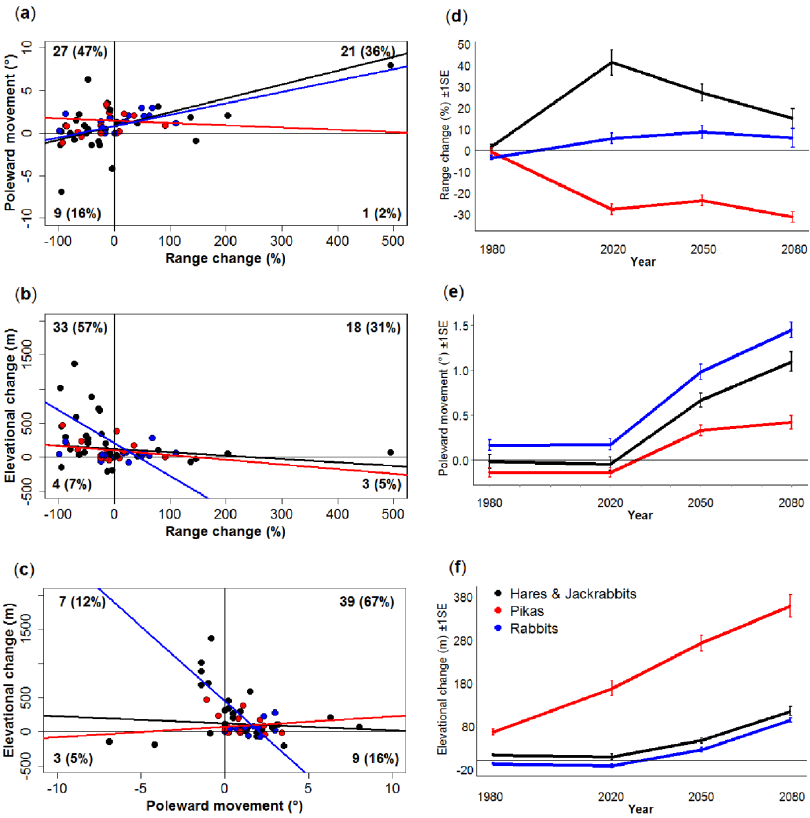
Characterisation of predicted lagomorph bioclimatic envelope change. Scatterplots show the linear relationship between range change (%) and (**a**) poleward movement (°), (**b**) elevational change (m) and (**c**) poleward movement and elevational change. The numbers of species in each quadrant that exhibited positive or negative change on each axis are shown in plain text with percentages in bold parentheses. Temporal trends for (**d**) range change, (**e**) poleward movement and (**f**) elevational change ± 1 standard error for each species group; pikas (red), rabbits (blue) and hares and jackrabbits (black).

**Figure 4.**
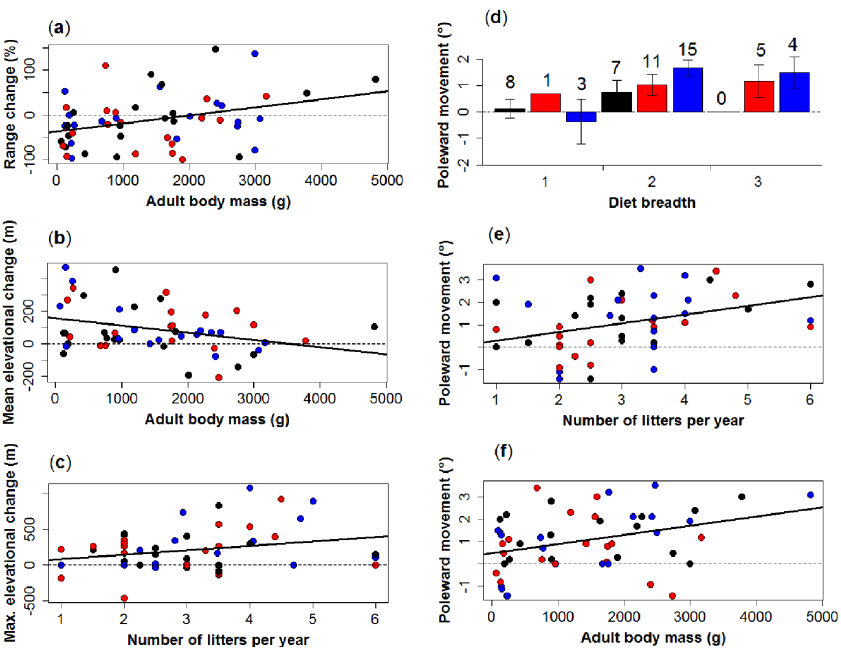
Relationships between species traits and responses to future climate change. The ability of species’ traits to predict changes in (**a**) range (**b**) mean elevation (**c**) maximum elevation and (**d**) (**e**) (**f**) poleward movement under future climate (between ~1930s and ~2080s) for each group; pikas (red), rabbits (blue) and hares and jackrabbits (black). Diet breadth is a categorical variable and is therefore represented as a bar plot (±1 standard error) with sample sizes (i.e. numbers of species) shown above the bars. Only significant predictors of change are shown here. The dashed line at zero indicates no change in the response variable.

## Appendix S1 Supplementary Methods

## Environmental parameterisation

Evapotranspiration was calculated using the Hargreaves equation [1]: 
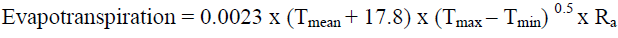

where,

R_a_: = ((24 × 60)/π) × G_sc_ × d_r_ [(ω_s_ × sinφ × sinδ)+(cosφ × cosδ × sinω_s_)]) × 0.408
G_sc_: = solar constant = 0.0820 MJ m-2 min-1
d_r_: = inverse relative distance Earth-Sun = 1 + 0.033 cos ((2π/365) × J)
ω_s_: = sunset hour angle [rad] = arcos [−tan (φ) tan (δ)]
φ: = latitude [radians] (grid file in decimal degrees converted to radians (multiply by π/180))
δ: = solar decimation [rad] = 0.409 sin (((2π/365) × J) – 1.39)
*J*: = number of days in the year. J at the middle of the month is approximately given by J = INTEGER (30.4 Month – 15) = 15 on average.

Annual evapotranspiration was taken as the sum of all monthly values. Annual water balance was calculated by subtracting annual evapotranspiration from mean annual precipitation. The number of months with a positive water balance was calculated by subtracting each monthly evapotranspiration from its corresponding monthly precipitation, and then converting these into a binary format, where a value greater than zero was given a value of one and a value less than zero was kept at zero [2]. The twelve binary files were then summed to calculate the number of months with a positive water balance.

Mean annual Normalised Difference Vegetation Index (NDVI) was calculated from monthly values which were downloaded from the EDIT Geoplatform (http://edit.csic.es/Soil-Vegetation-LandCover.html). NDVI is commonly used to measure the density of plant growth and is obtained from NOAA AVHRR satellite images. A negative value indicates snow or ice, a value around 0 indicates barren areas, values between 0.2 and 0.3 indicate grassland, and values near 1 indicate rainforests [3]. Human influence index was downloaded from the SEDAC website (http://sedac.ciesin.columbia.edu/data/set/wildareas-v2-human-influence-index-geographic). This was a composite of human population density, railways, roads, navigable rivers, coastlines, night-time lights, built-up areas and agricultural and urban land cover. Values within the index range from 0 to 64, where zero equalled no human influence and 64 represented maximum human influence [4]. Solar radiation was calculated using the Spatial Analyst function in ArcGIS 10.1 (ESRI, California, USA). Solar radiation is defined as the total amount of incoming solar insolation (direct and diffuse), or global radiation, and was measured in watt hours per square meter or WH/m^2^ [5]. An index of surface roughness was also calculated by finding the difference between maximum and minimum gradient values, based on a global Digital Elevation Model at 30 arc-second resolution downloaded from WorldClim [6]. Altitude was not included as a variable independently because organisms perceive climatic and habitat variables as proxies for altitude [7].

## Future Climate Data

Pierce *et al*. [8] report that using data averaged across five global circulation models (GCMs) is substantially better than any one individual model and significantly reduces model error. We averaged all variables for the following five GCMs: CCCma-CGCM3.1/T47, CNRM-CM3, CSIRO-Mk3.0, HadCM3 and NASA-GISS-ER. Future projections adopted the time periods: 2020s (2010-2039), 2050s (2040-2069) and 2080s (2070-2099). Data from the A2 future climate change scenario was used because although it was originally described as “extreme climate change” it now appears to best represent the trend in observed climate.

## Appendix S2

## Species Accounts (#1–89)

## Key for Response Curves

| Abbreviation | Description |
| --- | --- |
| **HII** | Human Influence index |
| **Hilliness** | Surface Roughness index |
| **MaxPrec** | Maximum precipitation |
| **MaxTemp** | Maximum temperature |
| **MinPrec** | Minimum precipitation |
| **MinTemp** | Minimum temperature |
| **PrecSea** | Precipitation seasonality |
| **RealMAR** | Mean annual precipitation |
| **RealMAT** | Mean annual temperature |
| **TempSea** | Temperature seasonality |
| **etpsum** | Annual evapotranspiration |
| **ndvi** | Normalised difference vegetation index |
| **radiation** | Solar radiation |
| **wbann** | Annual water balance |
| **wbpos** | Number of months with a positive water balance |

The past and current occurrence records underlying these species accounts can be viewed on http://lagomorphclimatechange.wordpress.com/.

**Species account #1 - Pygmy rabbit** (*Brachylagus idahoensis) *n* =*39

**Expert**: Penny Becker, Washington Dept. of Fish & Wildlife, USA

**Expert evaluation**: Medium

**Data**: Modern and historic

**Envelope**: Climatic and habitat

**Dispersal distance**: 15km/year (Expert)

**Status**: MODELLABLE; **Included in final analysis**: √

| Model evaluation metric |  |
| --- | --- |
| AUC | 0.95 |
| Omission rate | 0.10 |
| Sensitivity | 0.90 |
| Specificity | 1.00 |
| Proportion correct | 1.00 |
| **Kappa** | **0.75** |
| True Skill Statistic | 0.90 |

**Summary**: The Pygmy rabbit’s bioclimatic envelope is predicted to decline by 87% with a 1° mean latitudinal poleward shift and mean increase in elevation of ~300m driven predominately by an increase in mean minimum elevation (>600m) with little change in mean maximum elevation (~50m). 95% of the permutation importance of the model was contributed to by mean annual temperature (64.5%), maximum temperature (25.2%) and annual water balance (5.9%).

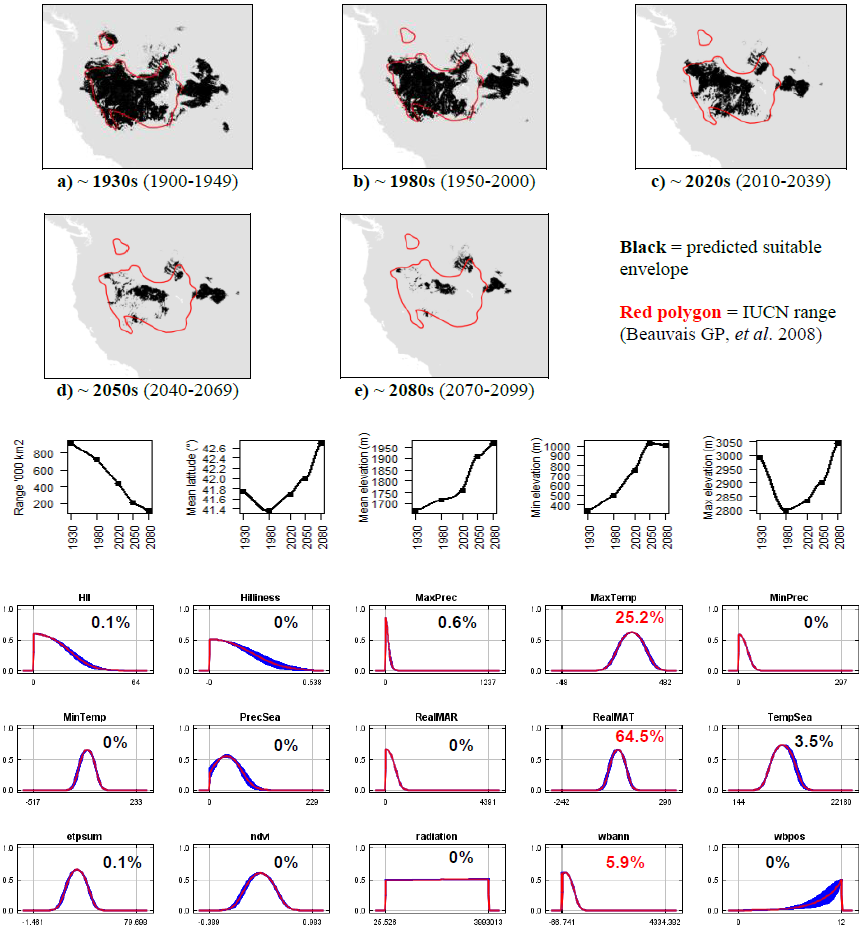

**#2 – Riverine rabbit** (*Bunolagus monticularis*) *n* = 109

**Status**: Kai Collins, University of Pretoria, South Africa

**Expert evaluation**: Good

**Data**: Modern and historic

**Envelope**: Climatic and habitat

**Dispersal distance**: 7.5km/year (Expert)

**Status**: MODELLABLE; **Included in final analysis**: √

| Model evaluation metric |  |
| --- | --- |
| AUC | 0.98 |
| Omission rate | 0.03 |
| Sensitivity | 0.97 |
| Specificity | 1.00 |
| Proportion correct | 1.00 |
| **Kappa** | **0.85** |
| True Skill Statistic | 0.97 |

**Summary**: The Riverine rabbit’s bioclimatic envelope is predicted to decline by 85% with a ~1° mean latitudinal poleward shift and mean increase in elevation of ~200m driven by similar increases in both minimum and maximum elevation. 95% of the permutation importance of the model was contributed to by minimum temperature (33.1%) and precipitation (22.5%), temperature seasonality (22.3%), mean annual temperature (6.7%) and precipitation (5.0%), annual evapotranspiration (4.3%) and precipitation seasonality (2.6%).

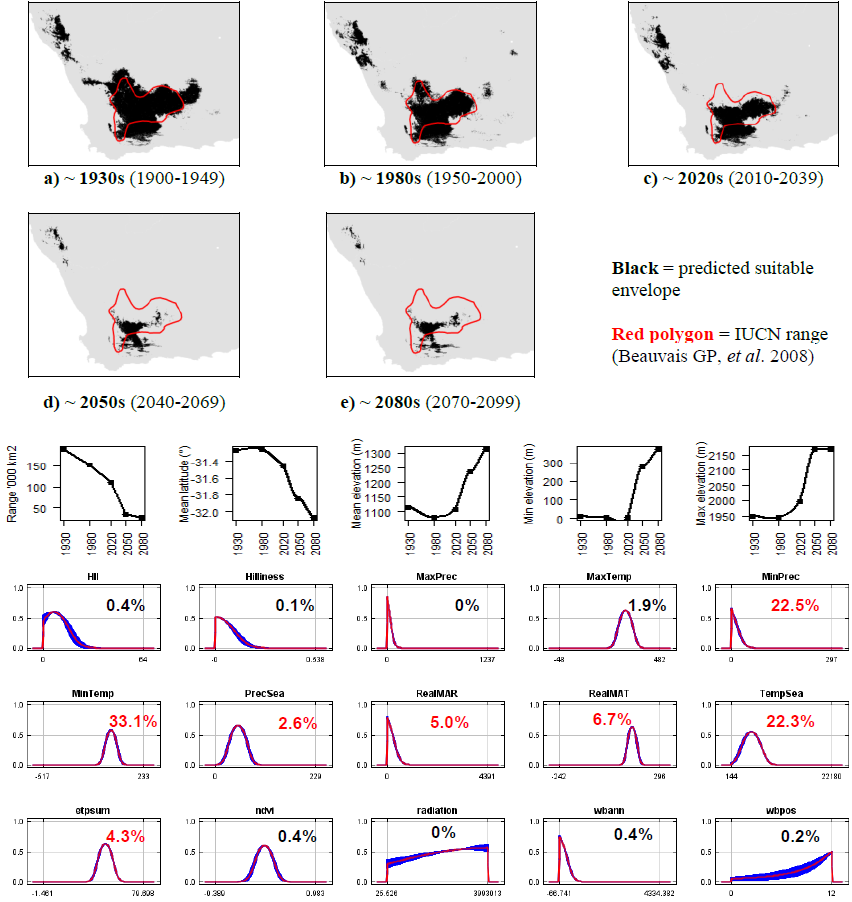

**#3 – Hispid hare** (*Caprolagus hispidus*) *n* = 18

**Expert**: Gopinathan Maheswaran, Zoological Survey of India

**Expert evaluation**: Medium

**Data**: Modern and historic

**Envelope**: Climatic and habitat

**Dispersal distance**: 5km/year (Expert)

**Status**: MODELLABLE; **Included in final analysis**: √

| Model evaluation metric |  |
| --- | --- |
| AUC | 0.97 |
| Omission rate | 0.06 |
| Sensitivity | 0.94 |
| Specificity | 0.99 |
| Proportion correct | 0.99 |
| **Kappa** | **0.81** |
| True Skill Statistic | 0.94 |

**Summary:** The Hispid hare’s bioclimatic envelope is predicted to increase by 21% with a ~1.5^o^ mean latitudinal poleward shift and mean increase in elevation of ~70m driven by increases in maximum elevation. 95% of the permutation importance of the model was contributed to by mean annual temperature (52.7%), precipitation seasonality (29.0%), annual evapotranspiration (6.6%), number of months with a positive water balance (2.9%), maximum precipitation (2.2%) and minimum temperature (1.7%).

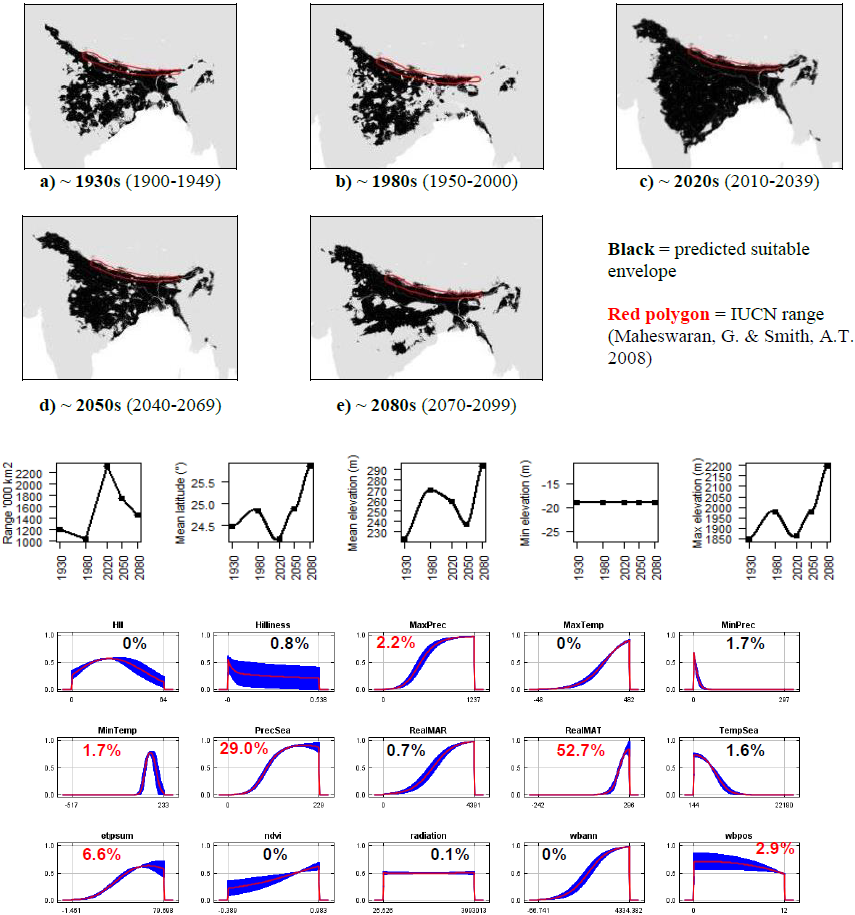

**#4 – Antelope jackrabbit** (*Lepus alleni*) *n* = 32

**Expert**: Paul Krausman, University of Montana

**Expert evaluation**: Medium

**Data**: Modern and historic

**Envelope**: Climatic and habitat

**Dispersal distance**: 25km/year (Expert)

**Status**: UNMODELLABLE; **Included in final analysis**: X

| Model evaluation metric |  |
| --- | --- |
| AUC | 0.91 |
| Omission rate | 0.16 |
| Sensitivity | 0.84 |
| Specificity | 0.99 |
| Proportion correct | 0.98 |
| **Kappa** | **0.26** |
| True Skill Statistic | 0.83 |

**Summary**: The Antelope jackrabbit’s bioclimatic envelope is predicted to increase by 172% with a ~3° mean latitudinal poleward shift and mean increase in elevation of ~20m driven by increases in maximum elevation. 95% of the permutation importance of the model was contributed to by precipitation seasonality (24.8%), minimum precipitation (21.1%), annual water balance (16.3%), minimum temperature (13.0%), temperature seasonality (7.3%), normalised difference vegetation index (5.8%), annual evapotranspiration (5.2%) and human influence index (3.7%).

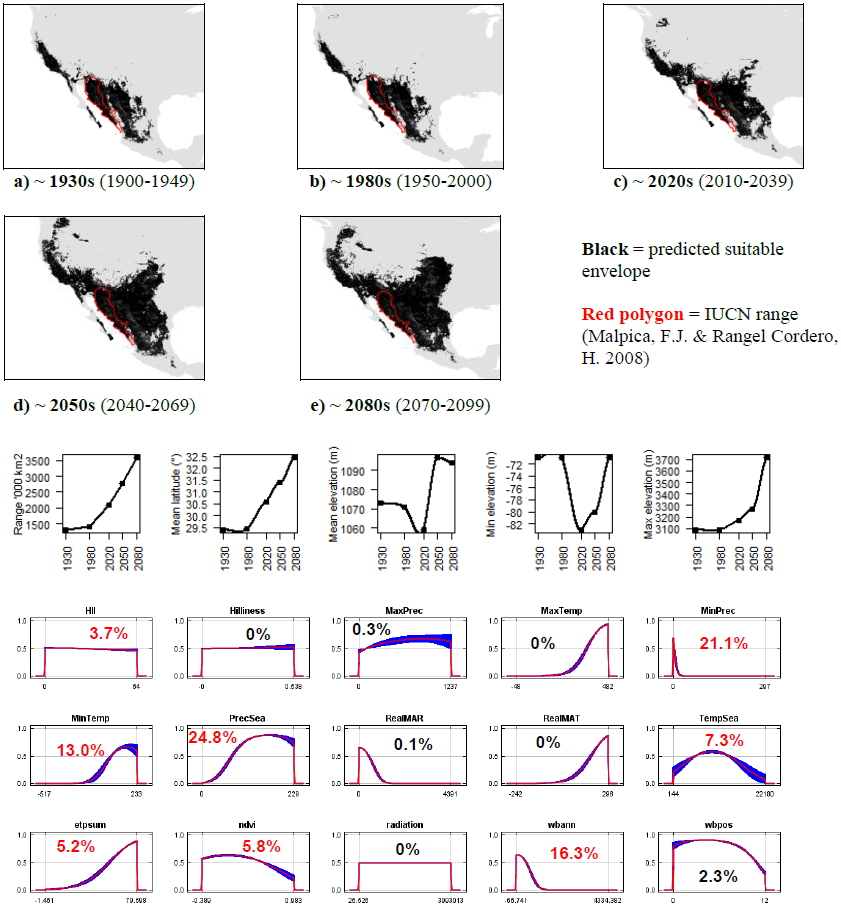

**#5 – Snowshoe hare** (*Lepus americanus*) *n* = 506

**Expert**: Charles Krebs, University of British Colombia & Rudy Boonstra, University of Toronto Scarborough

**Expert evaluation**: Good

**Data**: Only modern

**Envelope**: Climatic and habitat

**Dispersal distance**: 24km/year (Expert)

**Status**: MODELLABLE; **Included in final analysis**: √

| Model evaluation metric |  |
| --- | --- |
| AUC | 0.95 |
| Omission rate | 0.07 |
| Sensitivity | 0.93 |
| Specificity | 0.97 |
| Proportion correct | 0.97 |
| **Kappa** | **0.72** |
| True Skill Statistic | 0.90 |

**Summary:** The Snowshoe hare’s bioclimatic envelope is predicted to decline by 7% with a ~2° mean latitudinal poleward shift and mean decrease in elevation of ~10m, but with increases in both minimum and maximum elevation. 95% of the permutation importance of the model was contributed to by mean annual temperature (83.1%), maximum temperature (7.2%), normalised difference vegetation index (2.1%), mean annual precipitation (1.5%) and annual evapotranspiration (1.2%).

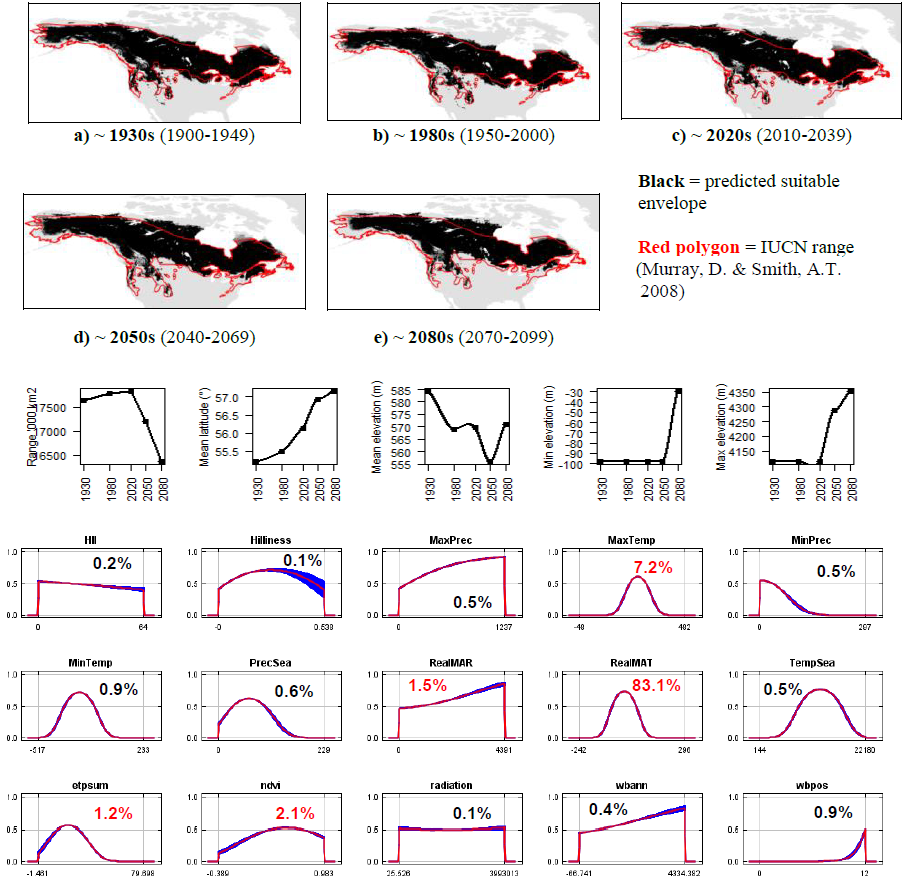

**#6 - Arctic hare** (*Lepus arcticus*) *n* = 18

**Expert**: David Gray, Grayhound Information Services

**Expert evaluation**: Poor

**Data**: Modern and historic

**Envelope**: Climatic and habitat

**Dispersal distance**: 2km/year (Chapman & Flux, 1990)

**Status**: UNMODELLABLE; **Included in final analysis**: X

| Model evaluation metric |  |
| --- | --- |
| AUC | 0.97 |
| Omission rate | 0.06 |
| Sensitivity | 0.94 |
| Specificity | 0.99 |
| Proportion correct | 0.99 |
| **Kappa** | **0.36** |
| True Skill Statistic | 0.94 |

**Summary**: The Arctic hare’s bioclimatic envelope is predicted to decline by 30% with a ~2° mean latitudinal poleward shift and mean decrease in elevation of ~10m driven by decreases in maximum elevation. 95% of the permutation importance of the model was contributed to by maximum temperature (34.0%), annual evapotranspiration (15.4%), surface roughness index (14.4%), human influence index (10.1%), normalised difference vegetation index (9.2%), mean annual temperature (4.4%), precipitation seasonality (1.5%), mean annual precipitation (1.5%), maximum precipitation (0.3%), minimum precipitation (0.3%), solar radiation (0.3%), minimum temperature (0.1%) and number of months with a positive water balance (0.1%).

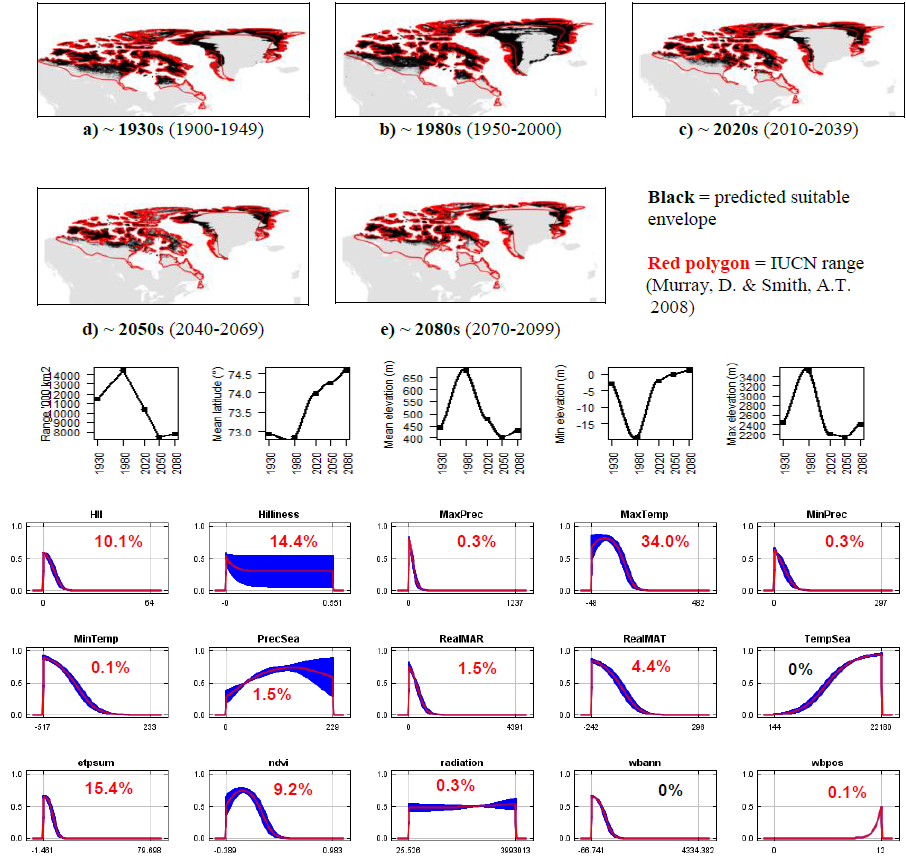

**#7 – Japanese hare** (*Lepus brachyurus*) *n* = 9

**Expert**: Koji Shimano, Shinshu University, Japan

**Expert evaluation**: Medium

**Data**: Modern and historic

**Envelope**: Climatic and habitat

**Dispersal distance**: 1km/year (Expert)

**Status**: MODELLABLE; **Included in final analysis**: √

| Model evaluation metric |  |
| --- | --- |
| AUC | 0.99 |
| Omission rate | 0.00 |
| Sensitivity | 1.00 |
| Specificity | 0.99 |
| Proportion correct | 0.99 |
| **Kappa** | **0.43** |
| True Skill Statistic | 0.99 |

**Summary**: The Japanese hare’s bioclimatic envelope is predicted to increase by 9% with no latitudinal poleward shift and a mean increase in elevation of ~10m driven by a decrease in minimum elevation. 95% of the permutation importance of the model was contributed to by temperature seasonality (31.8%), annual water balance (20.6%), human influence index (13.5%), mean annual precipitation (8.9%), maximum precipitation (6.2%), precipitation seasonality (5.4%) and minimum precipitation (4.4%).

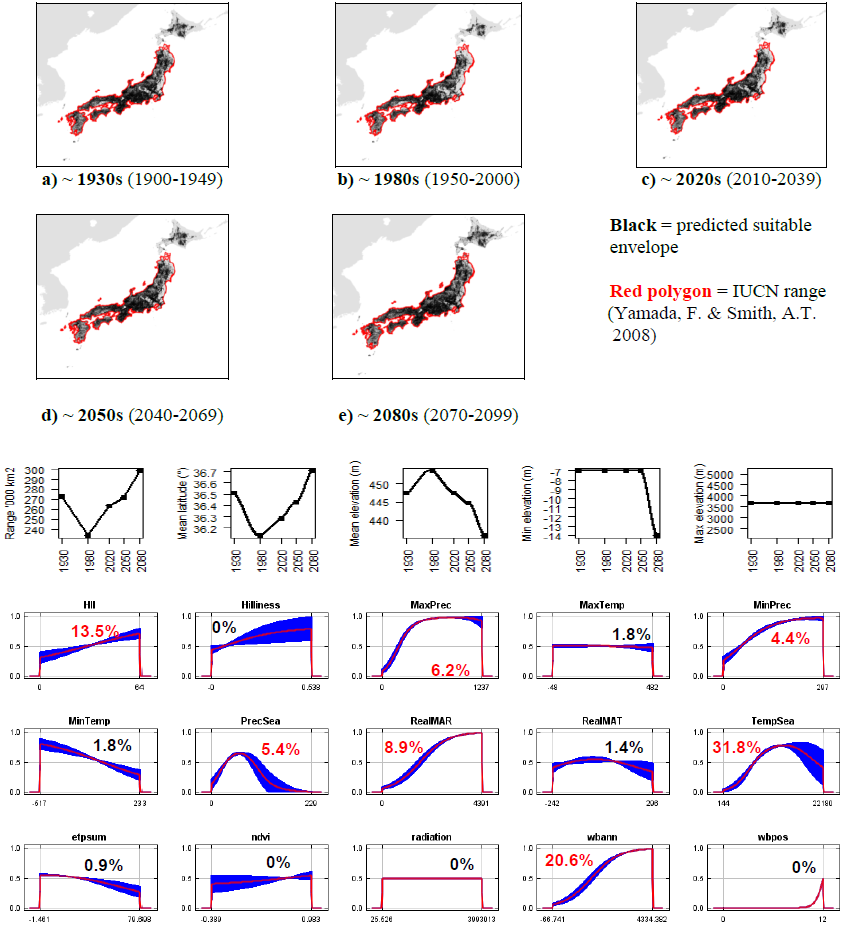

**#8 – Black-tailed jackrabbit** (*Lepus californicus*) *n* = 970

**Expert**: Alejandro Velasquez, UNAM-Canada

**Expert evaluation**: Good

**Data**: Modern and historic

**Envelope**: Climatic and habitat

**Dispersal distance**: 1.5km/year (Expert)

**Status**: MODELLABLE; **Included in final analysis**: √

| Model evaluation metric |  |
| --- | --- |
| AUC | 0.93 |
| Omission rate | 0.07 |
| Sensitivity | 0.93 |
| Specificity | 0.94 |
| Proportion correct | 0.94 |
| **Kappa** | **0.69** |
| True Skill Statistic | 0.87 |

**Summary:** The Black-tailed jackrabbit’s bioclimatic envelope is predicted to decline by 25% with a ~2° mean latitudinal poleward shift and mean decrease in elevation of ~75m, but with increases in both minimum and maximum elevation. 95% of the permutation importance of the model was contributed to by precipitation seasonality (31.8%), annual evapotranspiration (18.5%), maximum temperature (17.0%), mean annual temperature (8.9%), minimum temperature (8.0%), minimum precipitation (4.0%), human influence index (3.9%) and temperature seasonality (3.5%).

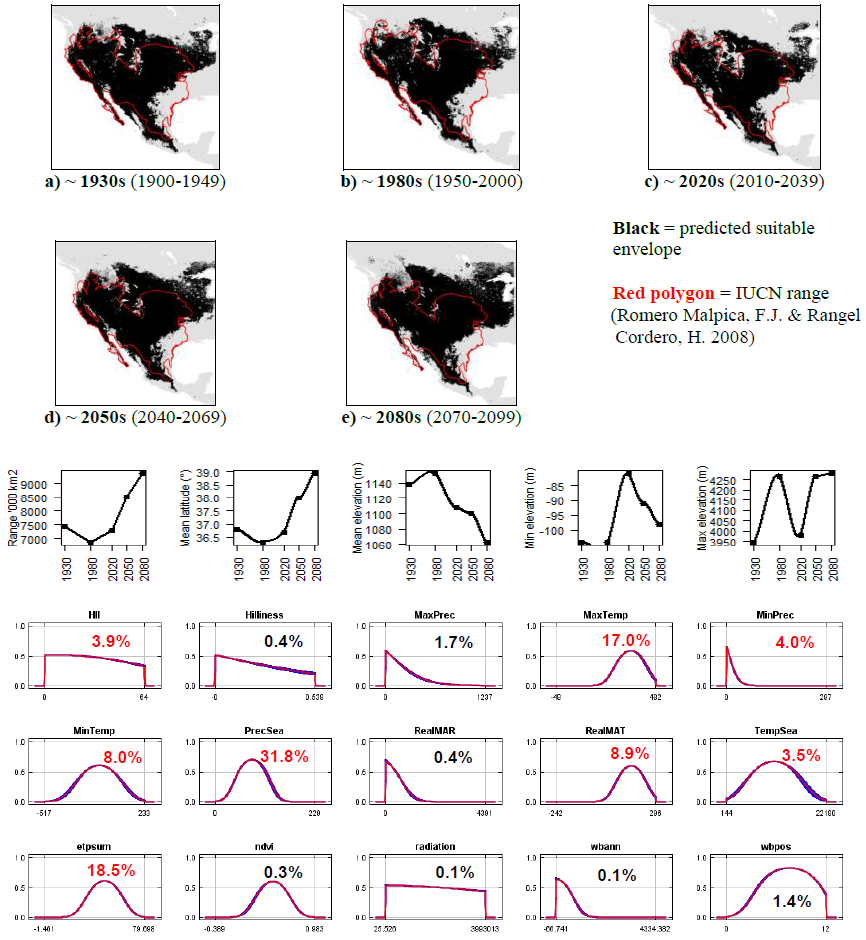

**#9 – White-sided jackrabbit** (*Lepus callotis*) *n* = 37

**Expert**: Jennifer Frey, New Mexico State University

**Expert evaluation**: Medium

**Data**: Modern and historic Envelope: Climatic and habitat

**Dispersal distance**: 18.9km/year (Average for NA leporids)

**Status**: UNMODELLABLE; **Included in final analysis**: X

| Model evaluation metric |  |
| --- | --- |
| AUC | 0.97 |
| Omission rate | 0.05 |
| Sensitivity | 0.95 |
| Specificity | 0.99 |
| Proportion correct | 0.99 |
| **Kappa** | **0.36** |
| True Skill Statistic | 0.93 |

**Summary:** The White-sided jackrabbit’s bioclimatic envelope is predicted to increase by 3% with a ~1° mean latitudinal poleward shift and a mean increase in elevation of ~150m driven by an increase in maximum elevation. 95% of the permutation importance of the model was contributed to by precipitation seasonality (35.4%), annual evapotranspiration (22.3%), minimum temperature (17.0%), mean annual temperature (10.5%), minimum precipitation (6.5%), maximum precipitation (6.5%), surface roughness index (3.0%) and number of months with a positive water balance (2.7%).

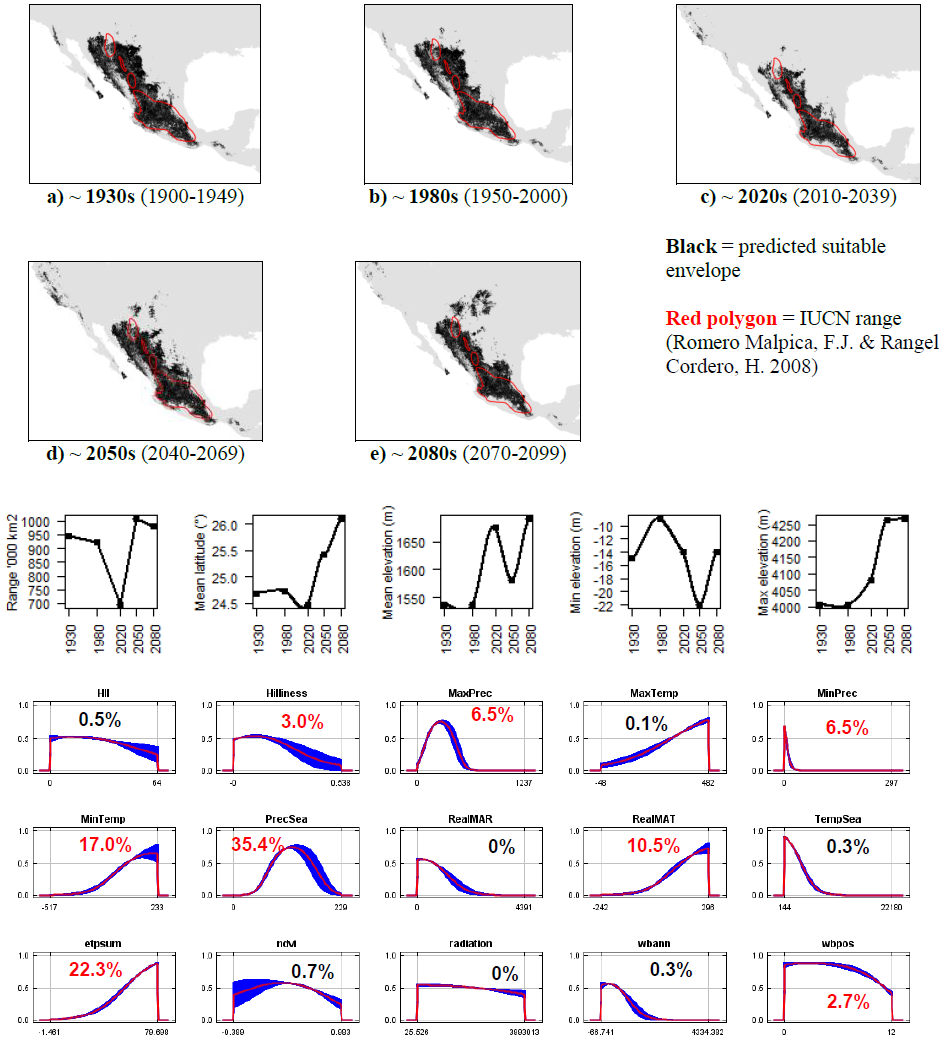

**#10 – Cape hare** (*Lepus capensis*) *n* = 231

**Expert**: John Flux, IUCN Lagomorph Specialist Group

**Expert evaluation**: Poor

**Data**: Modern and historic

**Envelope**: Climatic and habitat

**Dispersal distance**: 35km/year (Expert)

**Status**: UNMODELLABLE; **Included in final analysis**: X

| Model evaluation metric |  |
| --- | --- |
| AUC | 0.93 |
| Omission rate | 0.10 |
| Sensitivity | 0.90 |
| Specificity | 0.97 |
| Proportion correct | 0.97 |
| **Kappa** | **0.56** |
| True Skill Statistic | 0.87 |

**Summary**: The Cape hare’s bioclimatic envelope is predicted to decrease by 45% with ~2° mean latitudinal shift towards the Equator and a mean increase in elevation of ~330m driven by an increase in maximum elevation. 95% of the permutation importance of the model was contributed to by annual evapotranspiration (33.1%), minimum precipitation (29.6%), maximum temperature (9.7%), human influence index (7.2%), normalised difference vegetation index (4.6%), minimum temperature (3.2%), number of months with a positive water balance (2.9%), maximum precipitation (2.2%), mean annual precipitation (2.1%) and precipitation seasonality (2.0%).

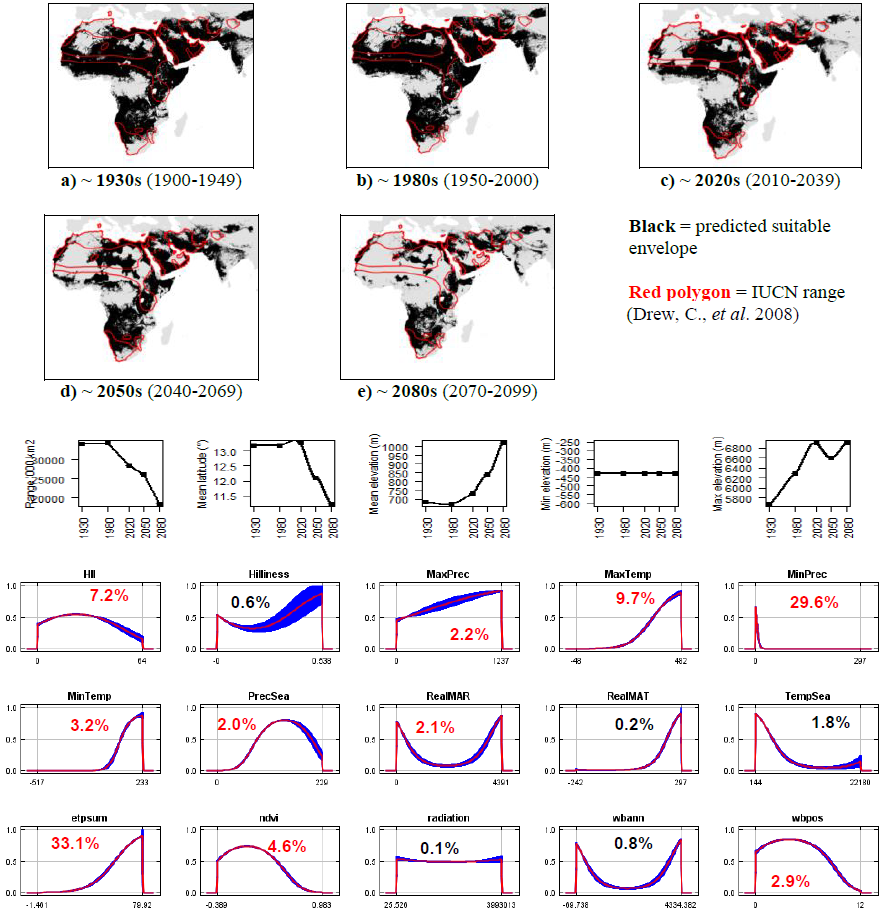

**#11 - Broom hare** (*Lepus castroviejoi*) *n* = 164

**Expert:** Pelayo Acevedo, University of Porto

**Expert evaluation:** Medium

**Data:** Only modern

**Envelope:** Climatic and habitat

**Dispersal distance:** 1km/year (Average for European leporids)

**Status:** MODELLABLE; **Included in final analysis:** √

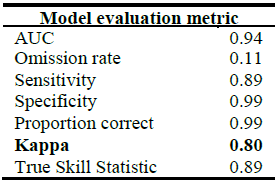

| Model evaluation metric |  |
| --- | --- |
| AUC | 0.94 |
| Omission rate | 0.11 |
| Sensitivity | 0.89 |
| Specificity | 0.99 |
| Proportion correct | 0.99 |
| **Kappa** | **0.80** |
| True Skill Statistic | 0.89 |

**Summary:** The Broom hare’s bioclimatic envelope is predicted to decrease by 90% with a ~0.2° mean latitudinal poleward shift and a mean increase in elevation of ~450m driven by an increase in minimum elevation. 95% of the permutation importance of the model was contributed to by mean annual temperature (62.0%), maximum temperature (20.6%), temperature seasonality (10.9%) and surface roughness index (3.8%).

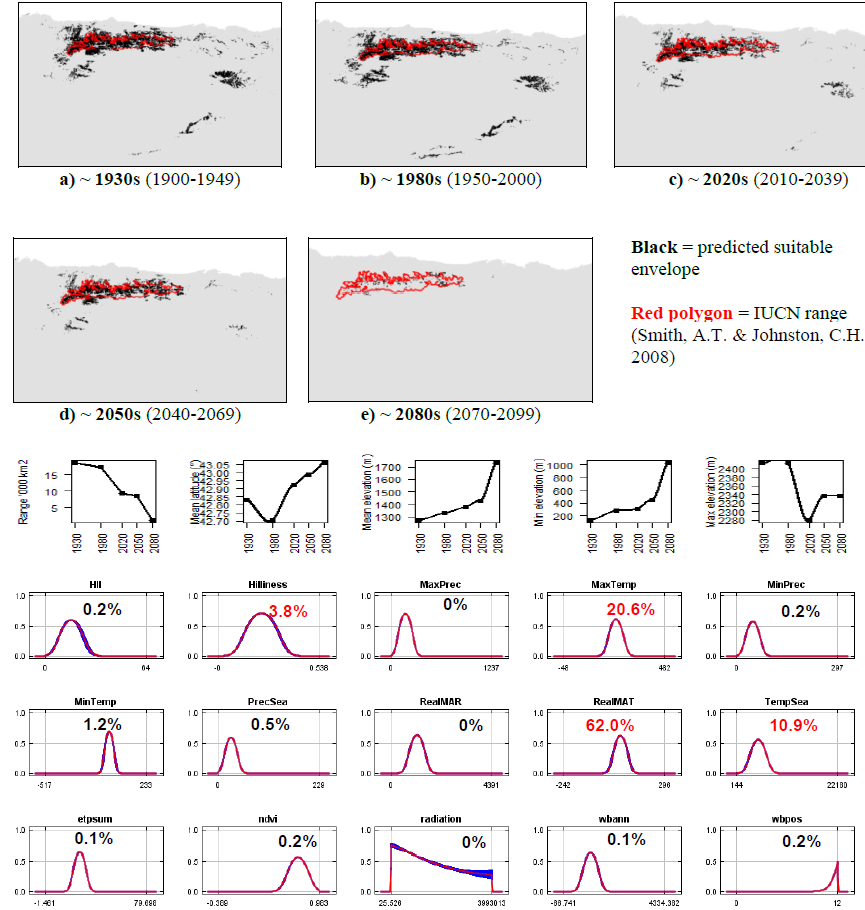

**#12 - Yunnan hare** (*Lepus comus*) *n* = 59

**Expert:** Weihe Yang, Institute of Zoology, Chinese Academy of Sciences

**Expert evaluation:** Medium

**Data:** Modern and historic Envelope: Climatic and habitat

**Dispersal distance:** 2.5km/year (Average for Asian leporids)

**Status:** MODELLABLE; **Included in final analysis:** √

| Model evaluation metric |  |
| --- | --- |
| AUC | 0.92 |
| Omission rate | 0.15 |
| Sensitivity | 0.85 |
| Specificity | 0.99 |
| Proportion correct | 0.99 |
| **Kappa** | **0.67** |
| True Skill Statistic | 0.84 |

**Summary:** The Yunnan hare’s bioclimatic envelope is predicted to decrease by 65% with a ~0.1° mean latitudinal poleward shift and a mean increase in elevation of ~100m driven by both increases in maximum and minimum elevation. 95% of the permutation importance of the model was contributed to by precipitation seasonality (59.9%), maximum temperature (26.8%), temperature seasonality (2.9%), number of months with a positive water balance (2.6%), annual evapotranspiration (2.0%) and mean annual temperature (1.7%).

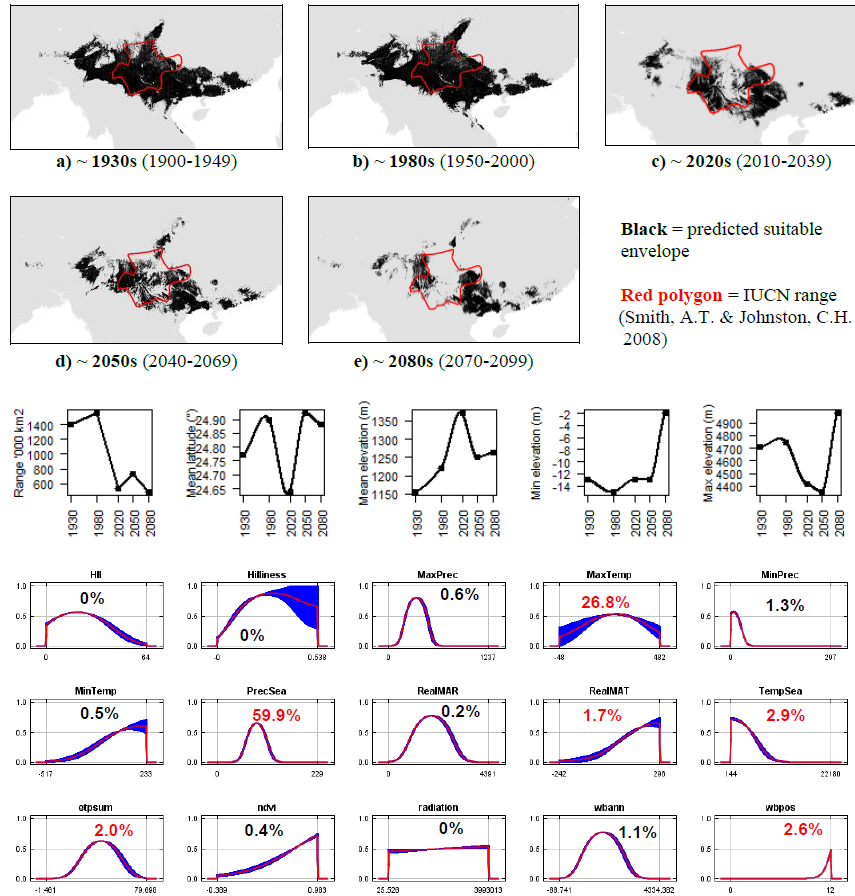

**#13 - Korean hare** (*Lepus coreanus*) *n* = 6

**Expert:** Weihe Yang, Institute of Zoology, Chinese Academy of Sciences

**Expert evaluation:** Medium

**Data:** Modern and historic

**Envelope:** Climatic and habitat

**Dispersal distance:** 2.5km/year (Average for Asian leporids)

**Status:** MODELLABLE; **Included in final analysis:** √

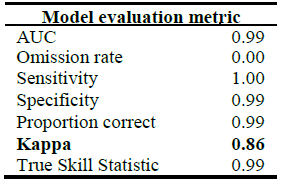

**Summary:** The Korean hare’s bioclimatic envelope is predicted to increase by 500% with a ~8° mean latitudinal poleward shift and a mean increase in elevation of ~70m driven by an increase in minimum elevation. 95% of the permutation importance of the model was contributed to by temperature seasonality (27.2%), mean annual precipitation (25.6%), minimum temperature (17.7%), annual water balance (13.1%), normalised difference vegetation index (4.7%), precipitation seasonality (4.5%) and human influence index (2.4%).

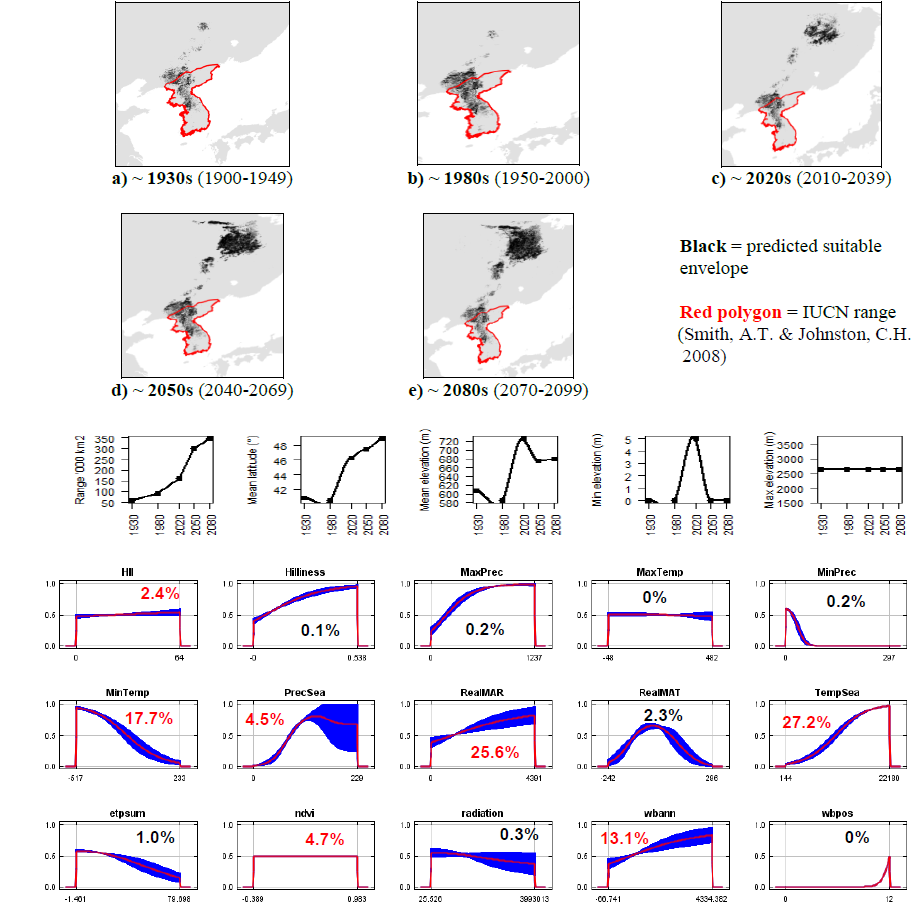

**#14 - Apennine hare** (*Lepus corsicanus*) *n* = 59

**Expert:** Francesco Angelici, Italian Foundation of Vertebrate Zoology

**Expert evaluation:** Medium

**Data:** Only modern

**Envelope:** Climatic and habitat

**Dispersal distance:** 3km/year (Expert)

**Status:** MODELLABLE; **Included in final analysis:** √

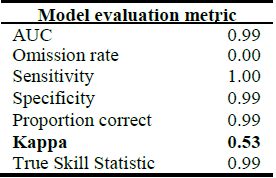

**Summary:** The Apennine hare’s bioclimatic envelope is predicted to increase by 125% with a ~2° mean latitudinal poleward shift and a mean decrease in elevation of ~60m. 95% of the permutation importance of the model was contributed to by minimum temperature (37.9%), annual evapotranspiration (34.3%), temperature seasonality (11.4%), minimum precipitation (11.1%) and maximum temperature (2.8%).

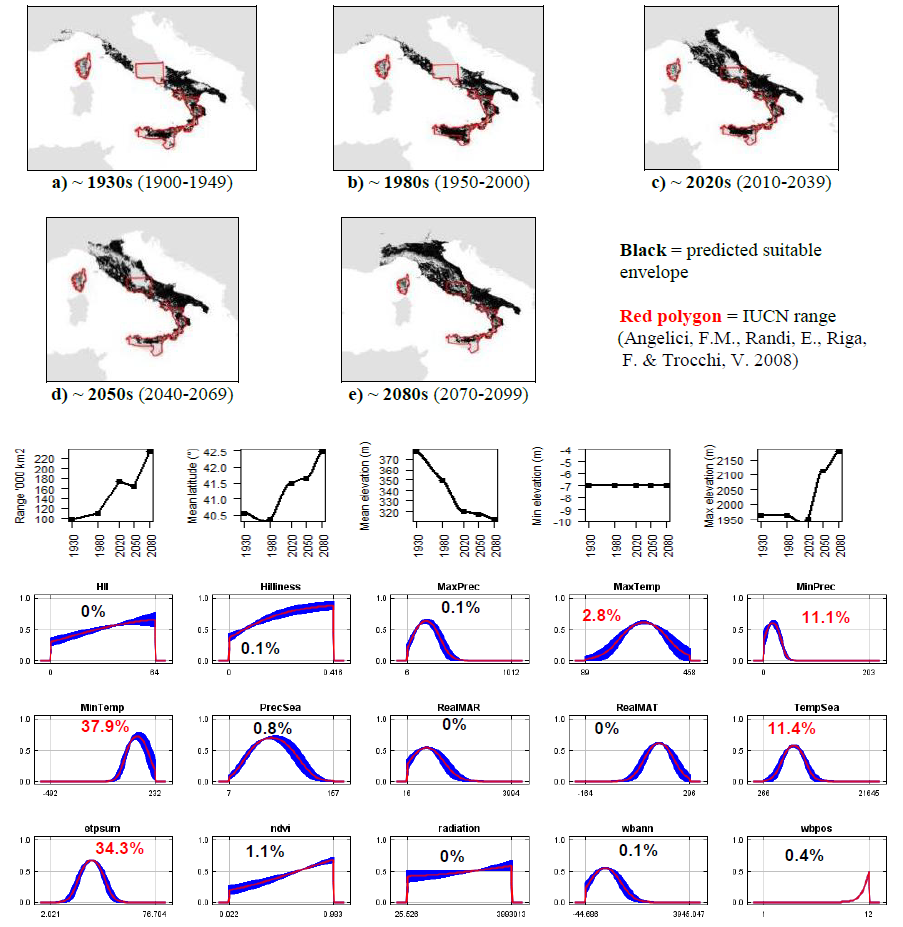

**#15 - European hare** (*Lepus europaeus*)-native range only *n* = 6,186

**Expert:** Neil Reid, Queen’s University Belfast

**Expert evaluation:** Medium

**Data:** Only modern

**Envelope:** Climatic and habitat Dispersal distance: 2km/year (Chapman & Flux, 1990)

**Status:** MODELLABLE; **Included in final analysis:** √

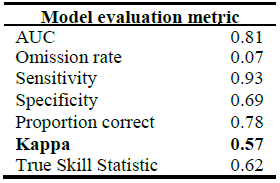

| Model evaluation metric |  |
| --- | --- |
| AUC | 0.81 |
| Omission rate | 0.07 |
| Sensitivity | 0.93 |
| Specificity | 0.69 |
| Proportion correct | 0.78 |
| **Kappa** | **0.57** |
| True Skill Statistic | 0.62 |

**Summary:** The European hare’s bioclimatic envelope is predicted to increase by 50% with a ~3° mean latitudinal poleward shift and a mean increase in elevation of ~20m driven by an increase in both maximum and minimum elevation. 95% of the permutation importance of the model was contributed to by annual evapotranspiration (58.0%), minimum temperature (9.0%), temperature seasonality (7.3%), surface roughness index (5.5%), human influence index (5.1%), precipitation seasonality (3.3%), annual water balance (3.3%), minimum precipitation (3.0%), normalised difference vegetation index (2.4%) and mean annual temperature (1.5%).

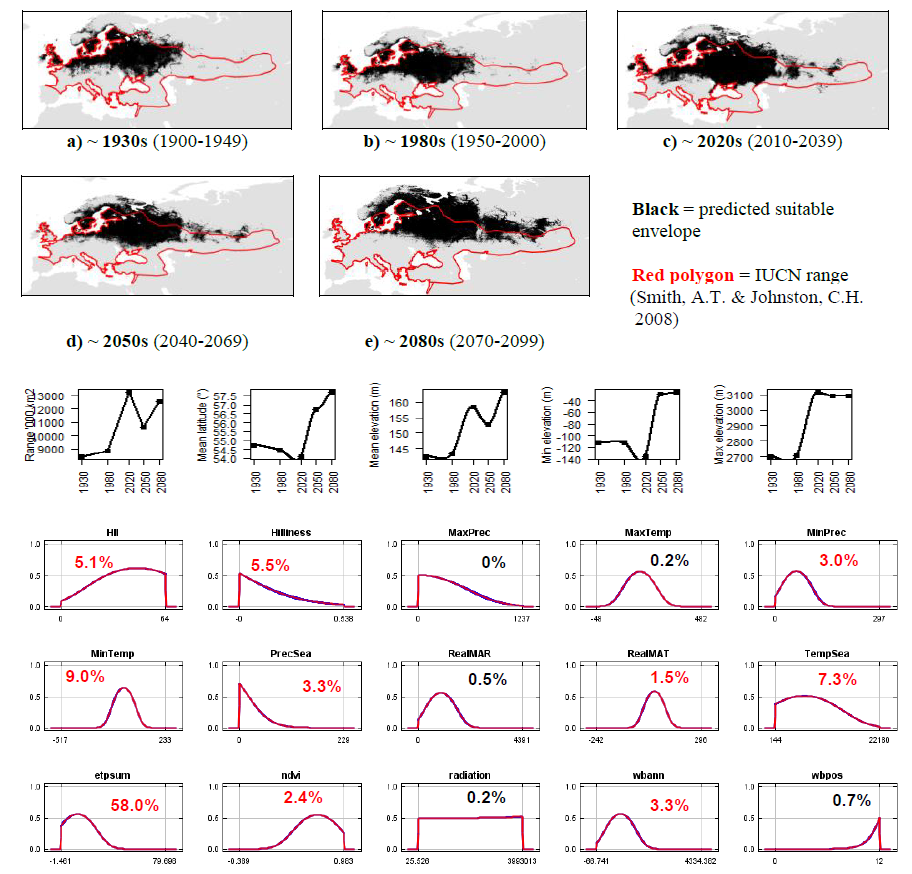

**#16 - Ethiopian hare** (*Lepus fagani*) *n* = 9

**Expert:** Zelalem Tolesa, Addis Ababa University

**Expert evaluation:** Poor

**Data:** Modern and historic

**Envelope:** Climatic and habitat

**Dispersal distance:** 8km/year (Average for African leporids)

**Status:** UNMODELLABLE; **Included in final analysis:** X

| Model evaluation metric |  |
| --- | --- |
| AUC | 0.99 |
| Omission rate | 0.00 |
| Sensitivity | 1.00 |
| Specificity | 0.99 |
| Proportion correct | 0.99 |
| **Kappa** | **0.56** |
| True Skill Statistic | 0.99 |

**Summary:** The Ethiopian hare’s bioclimatic envelope is predicted to decrease by 15% with no latitudinal poleward shift and a mean increase in elevation of ~200m driven by an increase in maximum and minimum elevation. 95% of the permutation importance of the model was contributed to by temperature seasonality (92.7%) and annual evapotranspiration (5.3%).

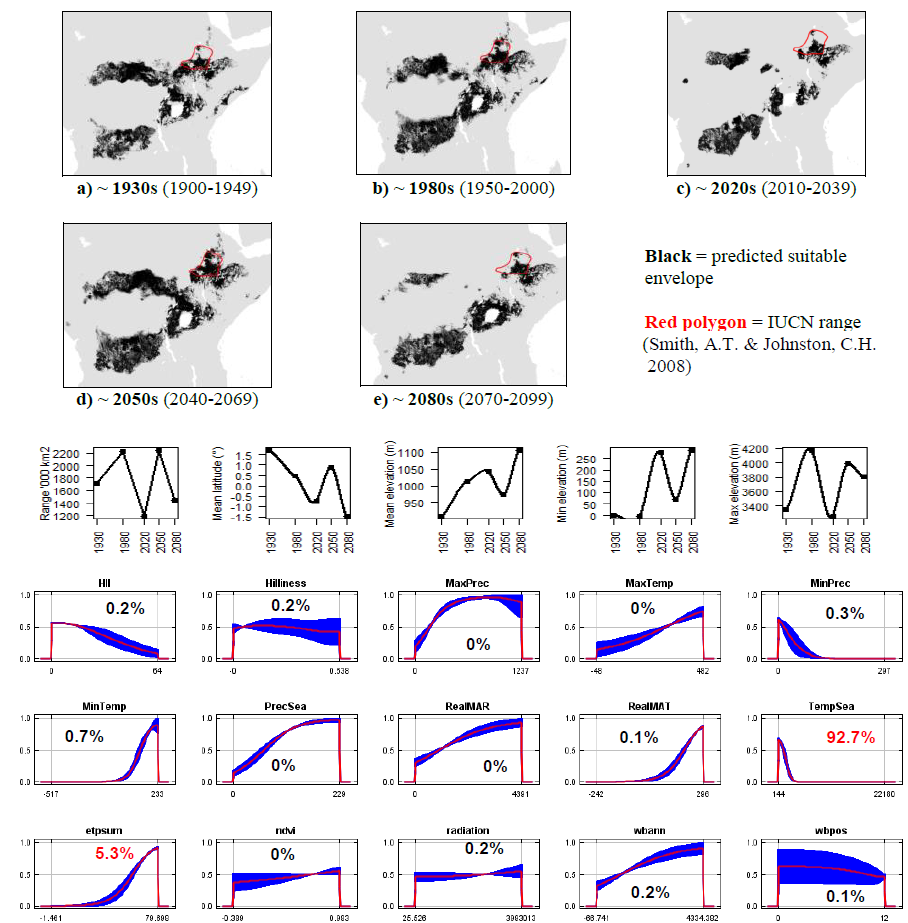

**#17 - Tehuantepec jackrabbit** (*Lepus flavigularis*) *n* = 8

**Expert:** Arturo Carillo-Reyes, Universidad de Ciencias y Artes de Chiapas

**Expert evaluation:** Poor

**Data:** Modern and historic

**Envelope:** Climatic and habitat

**Dispersal distance:** 0.01km/year (Expert)

**Status:** UNMODELLABLE; **Included in final analysis:** X

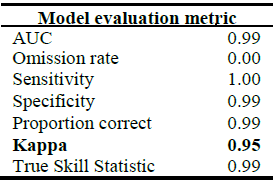

**Summary:** The Tehuantepec jackrabbit’s bioclimatic envelope is predicted to decrease by 45% with a ~1° mean latitudinal poleward shift and a mean increase in elevation of ~450m driven by an increase in maximum and minimum elevation. 95% of the permutation importance of the model was contributed to by temperature seasonality (81.7%), mean annual temperature (2.7%) and normalised difference vegetation index (1.7%).

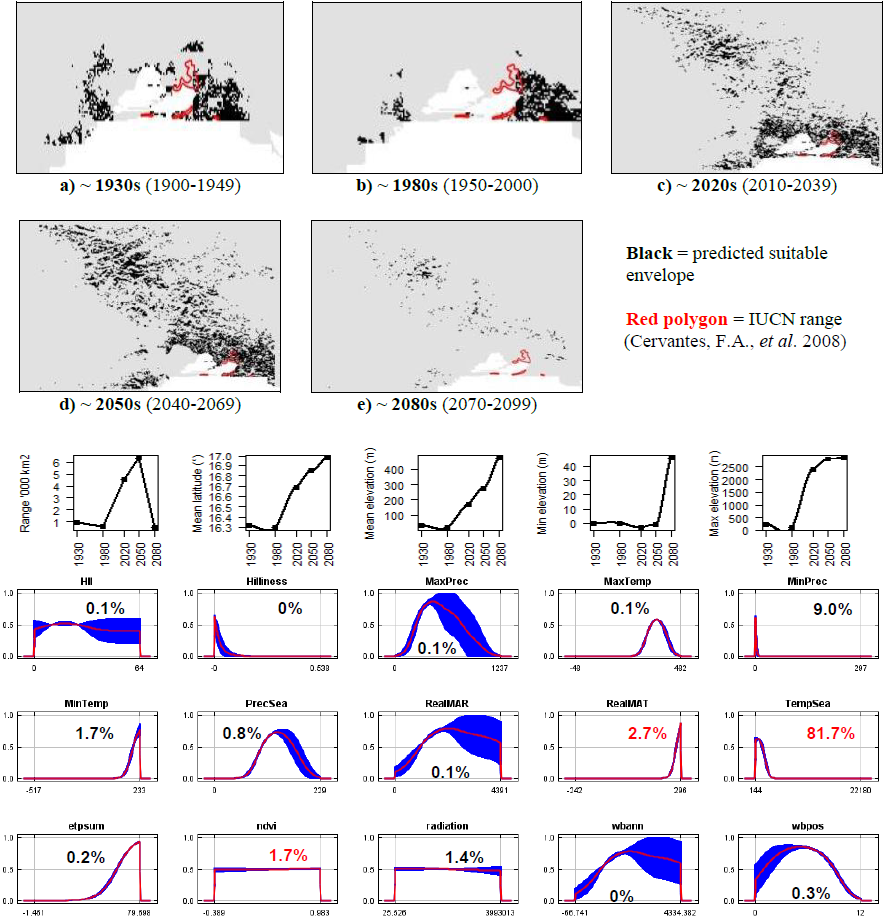

**#18 - Iberian hare** (*Lepus granatensis*) *n* = 1675

**Expert:** Pelayo Acevedo, University of Porto

**Expert evaluation:** Medium

**Data:** Modern and historic

**Envelope:** Climatic and habitat Dispersal distance: 7km/year (Expert)

**Status:** MODELLABLE; **Included in final analysis:** √

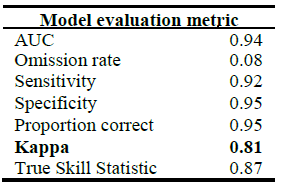

| Model evaluation metric |  |
| --- | --- |
| AUC | 0.94 |
| Omission rate | 0.08 |
| Sensitivity | 0.92 |
| Specificity | 0.95 |
| Proportion correct | 0.95 |
| **Kappa** | **0.81** |
| True Skill Statistic | 0.87 |

**Summary:** The Iberian hare’s bioclimatic envelope is predicted to increase by 40% with a ~1° mean latitudinal poleward shift and a mean increase in elevation of ~10m driven by an increase in maximum elevation. 95% of the permutation importance of the model was contributed to by maximum precipitation (39.0%), annual evapotranspiration (38.0%), minimum temperature (15.0%) and maximum temperature (3.0%).

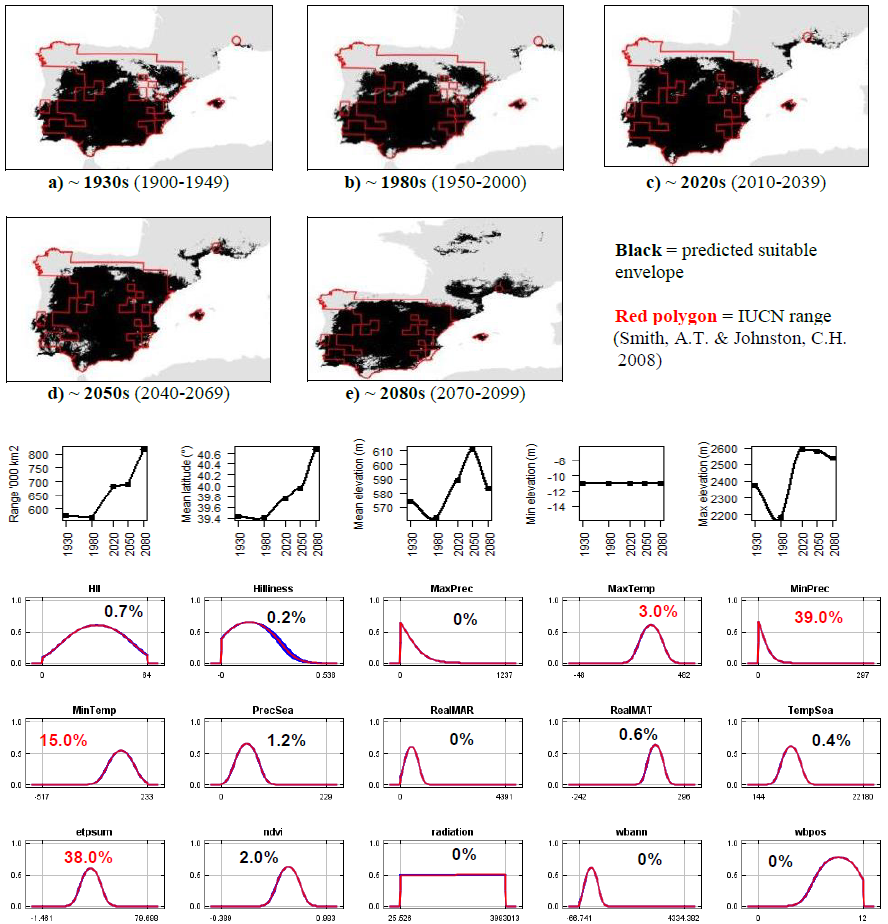

**#19 - Abyssinian hare** (*Lepus habessinicus*) *n* = 7

**Expert:** Zelalem Tolesa, Addis Ababa University

**Expert evaluation:** Medium

**Data:** Modern and historic

**Envelope:** Climatic and habitat

**Dispersal distance:** 25km/year (Average for African leporids)

**Status:** MODELLABLE; **Included in final analysis:** √

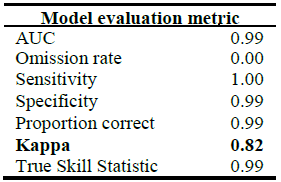

**Summary:** The Abyssinian hare’s bioclimatic envelope is predicted to decrease by 4% with a ~4° mean latitudinal shift towards the Equator and a mean decrease in elevation of ~200m driven by an decrease in maximum elevation. 95% of the permutation importance of the model was contributed to by temperature seasonality (87.6%), annual evapotranspiration (4.9%) and minimum precipitation (4.1%).

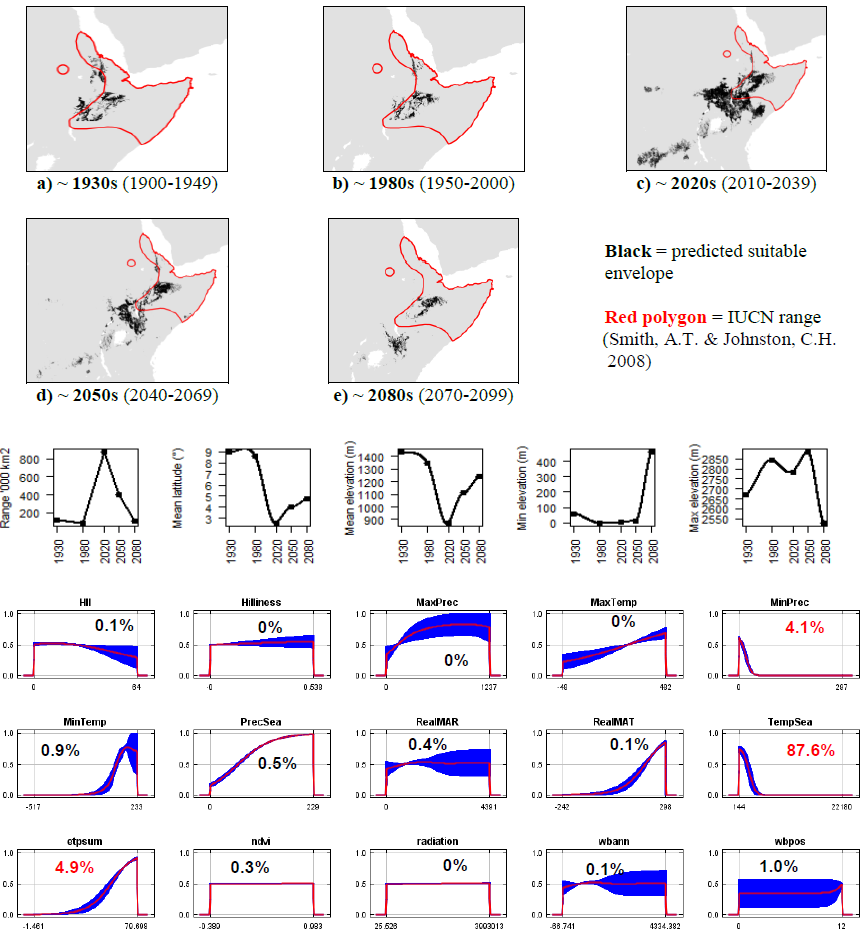

**#20 - Hainan hare** (*Lepus hainanus*) *n* = 9

**Expert:** Youhua Chen, Wuhan University, China

**Expert evaluation:** Good

**Data:** Modern and historic

**Envelope:** Climatic and habitat

**Dispersal distance:** 0.01km/year (Average for island leporids)

**Status:** MODELLABLE; **Included in final analysis:** √

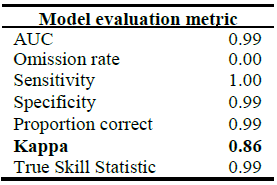

**Summary:** The Hainan hare’s bioclimatic envelope is predicted to increase by 4% with no latitudinal poleward shift and a mean increase in elevation of ~20m driven by an increase in maximum elevation. 95% of the permutation importance of the model was contributed to by minimum temperature (71.0%), mean annual temperature (22.8%) and temperature seasonality (5.4%).

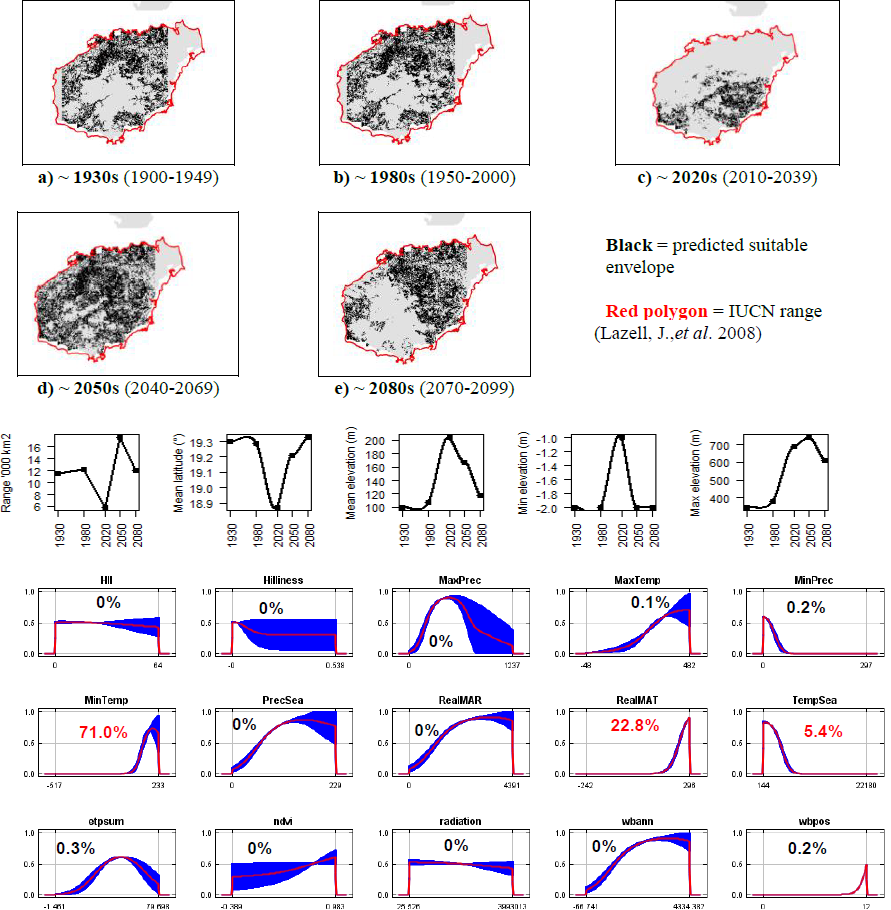

**#21 - Black jackrabbit** (*Lepus insularis*) *n* = 3

**Expert:** Tamara Rioja Pardela, Universidad de Ciencias y Artes de Chiapas, Mexico

**Expert evaluation:** Good

**Data:** Modern and historic

**Envelope:** Climatic and habitat

**Dispersal distance:** 0.01km/year (Average for island leporids)

**Status:** MODELLABLE; **Included in final analysis:** √

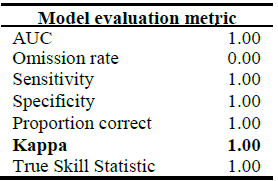

**Summary:** The Black jackrabbit’s bioclimatic envelope is predicted to decrease by 100% with a ~0.3° mean latitudinal polewards shift and a mean increase in elevation of ~50m driven by an increase in both minimum and maximum elevation. 95% of the permutation importance of the model was contributed to by precipitation seasonality (28.6%), minimum precipitation (16.5%), annual water balance (13.3%), minimum temperature (12.9%), mean annual temperature (7.1%), temperature seasonality (5.5%), surface roughness index (5.0%) and normalised difference vegetation index (3.2%).

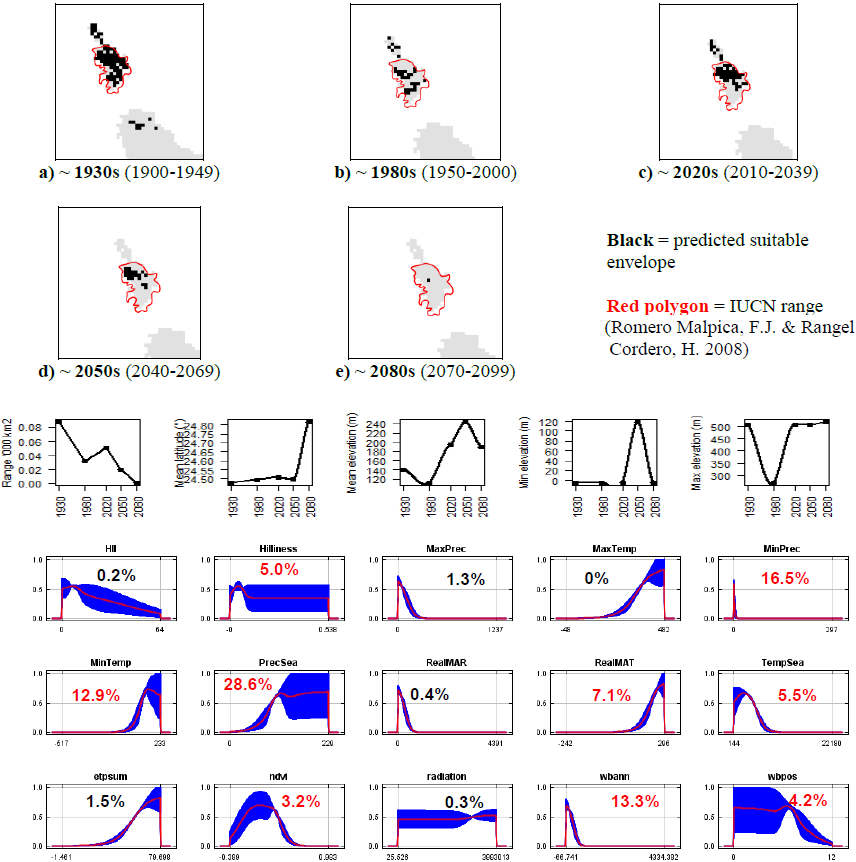

**#22 - Manchurian hare** (*Lepus mandshuricus*) *n* = 36

**Expert:** Deyan Ge, Institute of Zoology, Chinese Academy of Sciences

**Expert evaluation:** Medium

**Data:** Modern and historic

**Envelope:** Climatic and habitat

**Dispersal distance:** 3km/year (Sokolov, V.E. *et al.,* 2009)

**Status:** MODELLABLE; **Included in final analysis:** √

| Model evaluation metric |  |
| --- | --- |
| AUC | 0.96 |
| Omission rate | 0.08 |
| Sensitivity | 0.92 |
| Specificity | 0.99 |
| Proportion correct | 0.99 |
| **Kappa** | **0.78** |
| True Skill Statistic | 0.92 |

**Summary:** The Manchurian hare’s bioclimatic envelope is predicted to decrease by 50% with a ~1° mean latitudinal polewards shift and a mean increase in elevation of ~70m driven by an increase in maximum and minimum elevation. 95% of the permutation importance of the model was contributed to by precipitation seasonality (77.3%), mean annual temperature (10.4%), minimum temperature (4.5%), normalised difference vegetation index (2.6%) and annual evapotranspiration (2.4%).

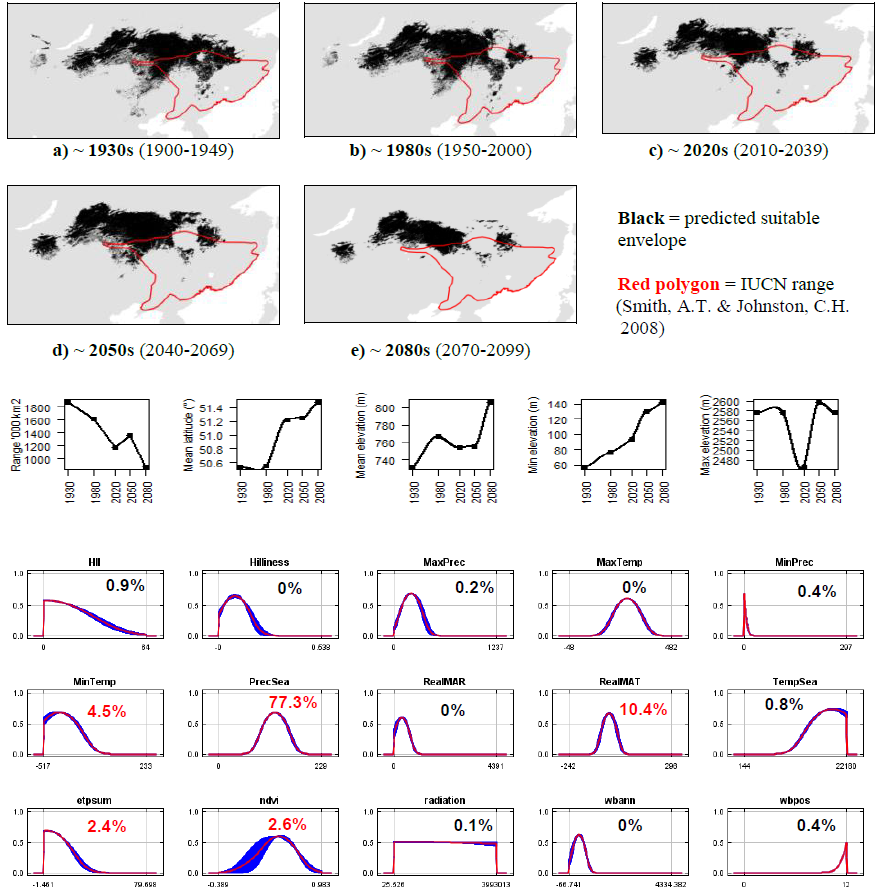

**#23 - African savannah hare** (*Lepus microtis*) *n* = 82

**Expert:** John Flux, IUCN Lagomorph Specialist Group

**Expert evaluation:** Medium

**Data:** Modern and historic

**Envelope:** Climatic only Dispersal distance: 15km/year (Expert)

**Status:** MODELLABLE; **Included in final analysis:** √

| Model evaluation metric |  |
| --- | --- |
| AUC | 0.93 |
| Omission rate | 0.13 |
| Sensitivity | 0.87 |
| Specificity | 0.99 |
| Proportion correct | 0.99 |
| **Kappa** | **0.62** |
| True Skill Statistic | 0.86 |

**Summary:** The African savannah hare’s bioclimatic envelope is predicted to decrease by 15% with a ~3° mean latitudinal polewards shift and a mean increase in elevation of ~100m driven by an increase in maximum and minimum elevation. 95% of the permutation importance of the model was contributed to by mean annual temperature (40.9%), maximum temperature (18.5%), annual evapotranspiration (9.2%), minimum precipitation (6.2%), temperature seasonality (5.4%), mean annual precipitation (5.4%), minimum temperature (4.9%), precipitation seasonality (4.1%) and annual water balance (1.7%).

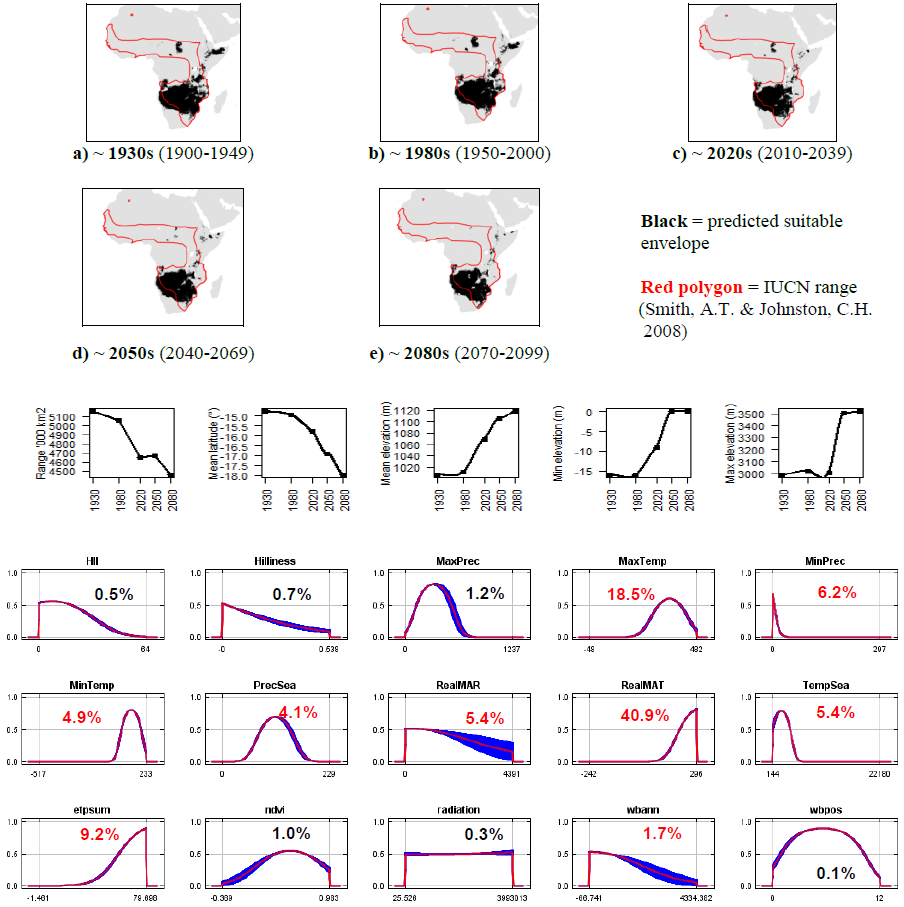

**#24 - Indian hare** (*Lepus nigricollis*) *n* = 17

**Expert:** Gopinathan Maheswaran, Zoological Survey of India

**Expert evaluation:** Good

**Data:** Modern and historic

**Envelope:** Climatic and habitat

**Dispersal distance:** 6km/year (Expert)

**Status:** MODELLABLE; **Included in final analysis:** √

| Model evaluation metric |  |
| --- | --- |
| AUC | 0.99 |
| Omission rate | 0.00 |
| Sensitivity | 1.00 |
| Specificity | 0.99 |
| Proportion correct | 0.99 |
| **Kappa** | **0.59** |
| True Skill Statistic | 0.99 |

**Summary:** The Indian hare’s bioclimatic envelope is predicted to decrease by 10% with a ~2° mean latitudinal polewards shift and a mean increase in elevation of ~80m driven by an increase in maximum and minimum elevation. 95% of the permutation importance of the model was contributed to by precipitation seasonality (48.0%), mean annual temperature (14.9%), human influence index (11.9%), minimum temperature (10.3%), temperature seasonality (7.0%), number of months with a positive water balance (2.8%) and minimum precipitation (1.5%).

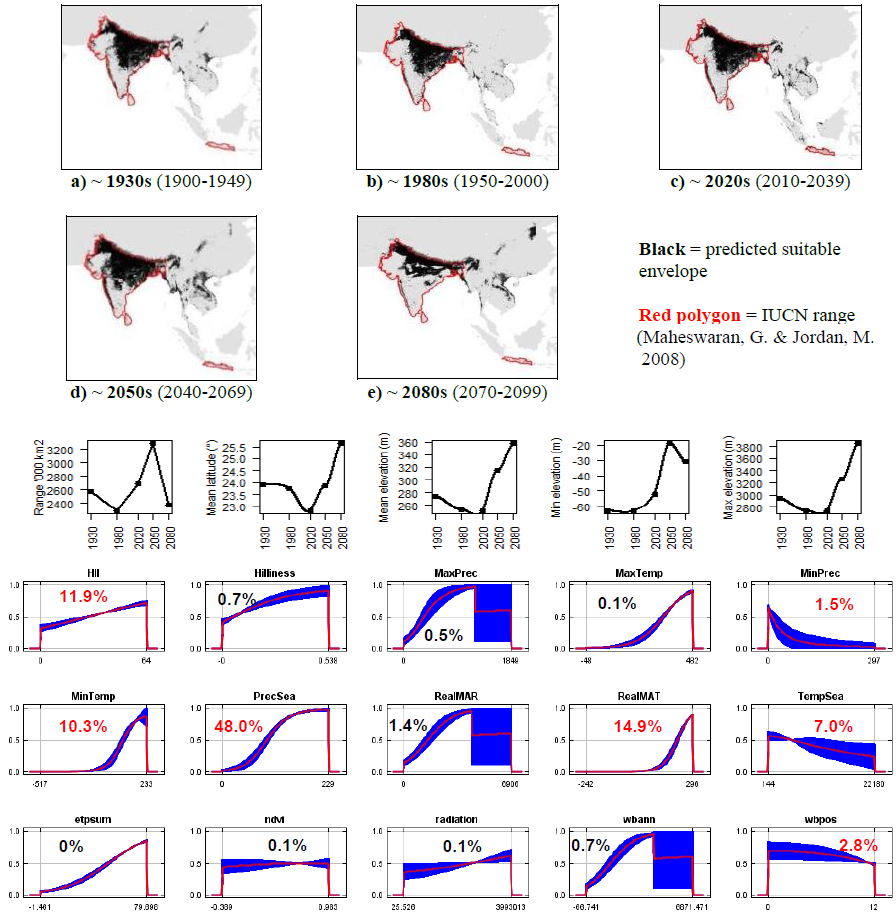

**#25 - Woolly hare** (*Lepus oiostolus*) *n* = 84

**Expert:** Weihe Yang, Institute of Zoology, Chinese Academy of Sciences

**Expert evaluation:** Medium

**Data:** Only modern Envelope: Climatic and habitat

**Dispersal distance:** 2.5km/year (Average for Asian leporids)

**Status:** MODELLABLE; **Included in final analysis:** √

| Model evaluation metric |  |
| --- | --- |
| AUC | 0.94 |
| Omission rate | 0.11 |
| Sensitivity | 0.89 |
| Specificity | 0.99 |
| Proportion correct | 0.99 |
| **Kappa** | **0.63** |
| True Skill Statistic | 0.89 |

**Summary:** The Woolly hare’s bioclimatic envelope is predicted to decrease by 25% with a ~1° mean latitudinal shift towards the Equator and a mean increase in elevation of ~680m driven by an increase in maximum and minimum elevation. 95% of the permutation importance of the model was contributed to by minimum precipitation (82.1%), maximum temperature (5.0%), minimum temperature (4.6%) and annual evapotranspiration (3.8%).

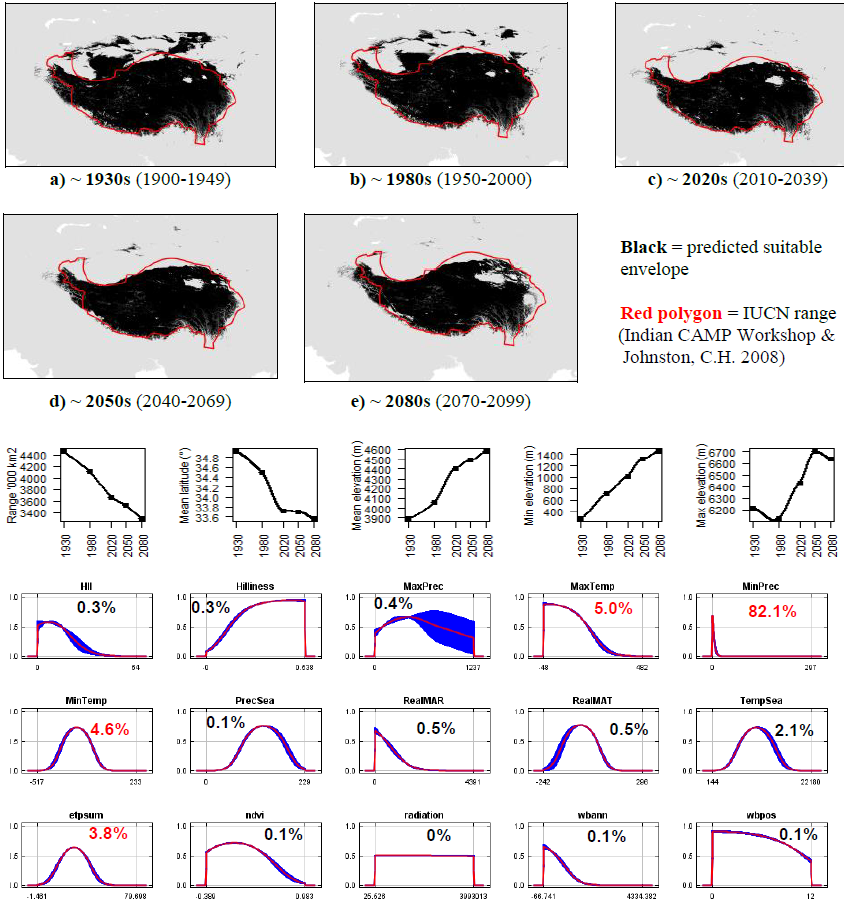

**#26 - Alaskan hare** (*Lepus othus*) *n* = 8

**Expert:** Eric Waltari, City University of New York

**Expert evaluation:** Medium

**Data:** Modern and historic

**Envelope:** Climatic only

**Dispersal distance:** 2km/year (Average for Arctic leporids)

**Status:** MODELLABLE; **Included in final analysis:** √

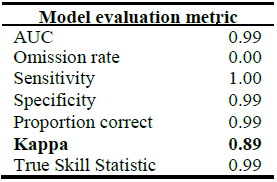

**Summary:** The Alaskan hare’s bioclimatic envelope is predicted to increase by 80% with a ~3° mean latitudinal polewards shift and a mean increase in elevation of ~100m driven by an increase in minimum elevation. 95% of the permutation importance of the model was contributed to by annual evapotranspiration (52.8%), mean annual temperature (26.4%), precipitation seasonality (11.3%) and mean annual precipitation (5.0%).

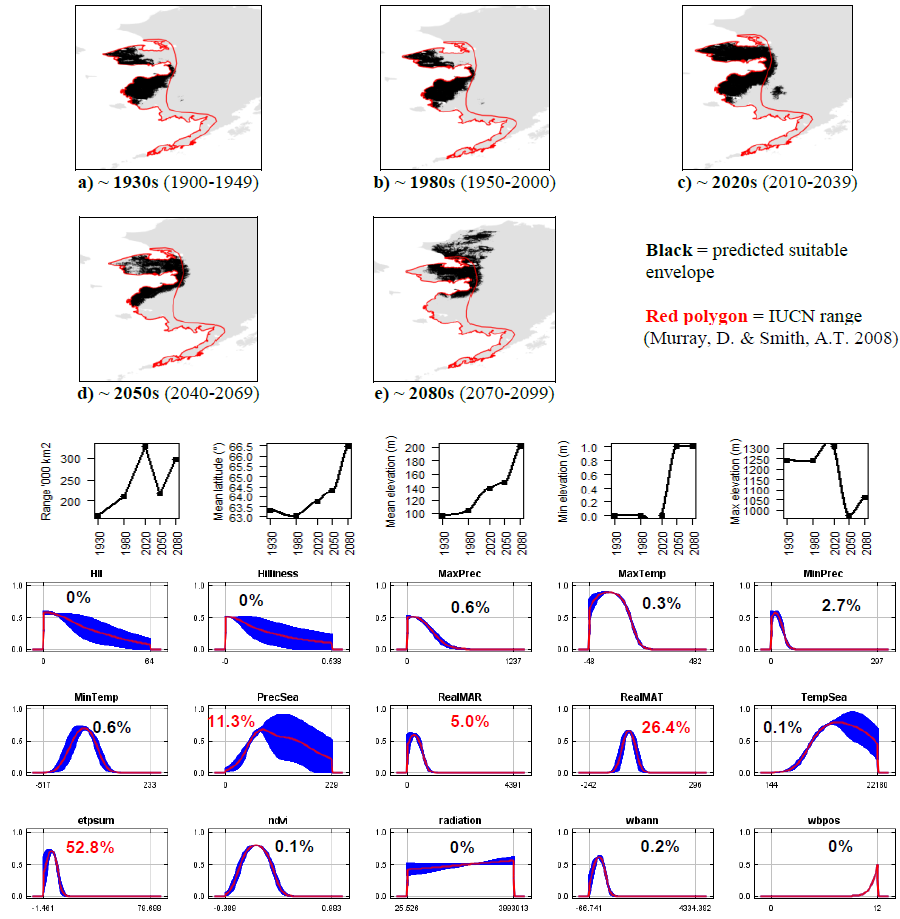

**#27 - Burmese hare** (*Lepus peguensis*) *n* = 7

**Expert:** Thomas Gray, WWF Greater Mekong

**Expert evaluation:** Medium

**Data:** Modern and historic

**Envelope:** Climatic and habitat

**Dispersal distance:** 2.5km/year (Average for Asian leporids)

**Status:** MODELLABLE; **Included in final analysis:** √

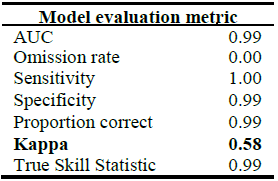

**Summary:** The Burmese hare’s bioclimatic envelope is predicted to increase by 40% with a ~2° mean latitudinal polewards shift and a mean increase in elevation of ~180m driven by an increase in minimum and maximum elevation. 95% of the permutation importance of the model was contributed to by temperature seasonality (52.6%), minimum precipitation (32.9%), solar radiation (6.5%) and mean annual temperature (4.3%).

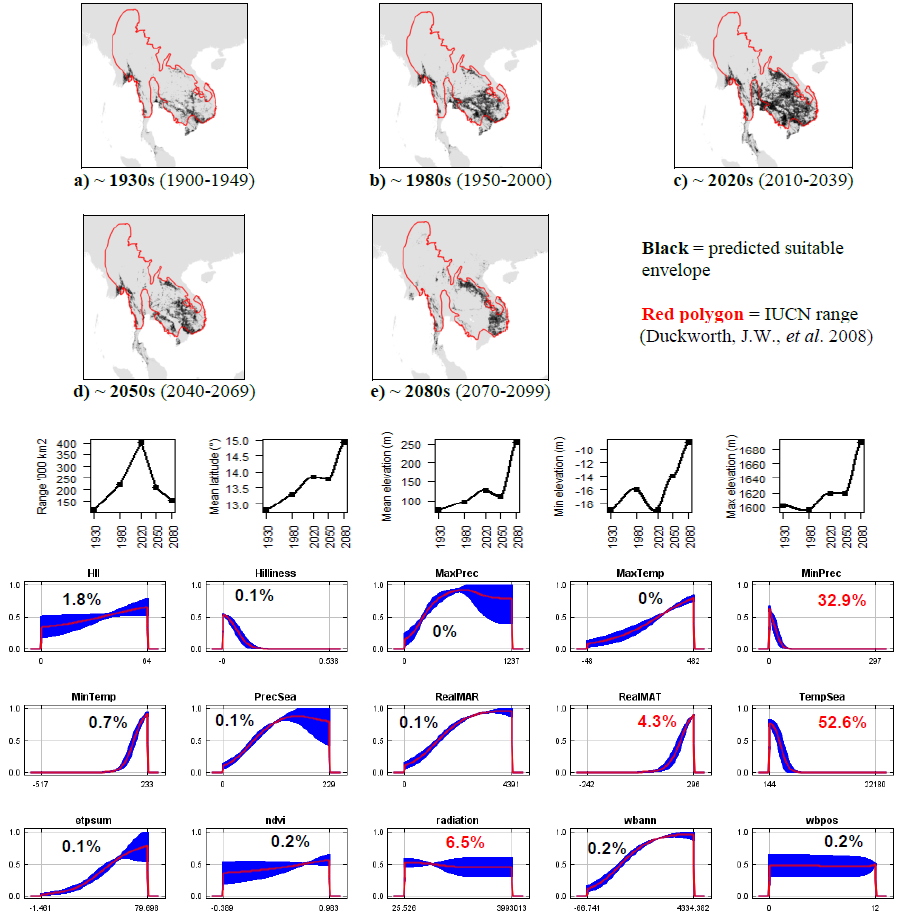

**#28 - Scrub hare** (*Lepus saxatilis*) *n* = 39

**Expert:** Kai Collins, University of Pretoria

**Expert evaluation:** Poor

**Data:** Only modern

**Envelope:** Climatic and habitat

**Dispersal distance:** 25km/year (Average for African leporids)

**Status:** UNMODELLABLE; **Included in final analysis:** X

| Model evaluation metric |  |
| --- | --- |
| AUC | 0.95 |
| Omission rate | 0.08 |
| Sensitivity | 0.92 |
| Specificity | 0.97 |
| Proportion correct | 0.97 |
| **Kappa** | **0.18** |
| True Skill Statistic | 0.89 |

**Summary:** The Scrub hare’s bioclimatic envelope is predicted to decrease by 15% with a ~2° mean latitudinal shift towards the Equator and a mean increase in elevation of ~65m driven by an increase in maximum and minimum elevation. 95% of the permutation importance of the model was contributed to by annual evapotranspiration (52.8%), mean annual temperature (26.4%), precipitation seasonality (11.3%) and mean annual precipitation (5.0%).

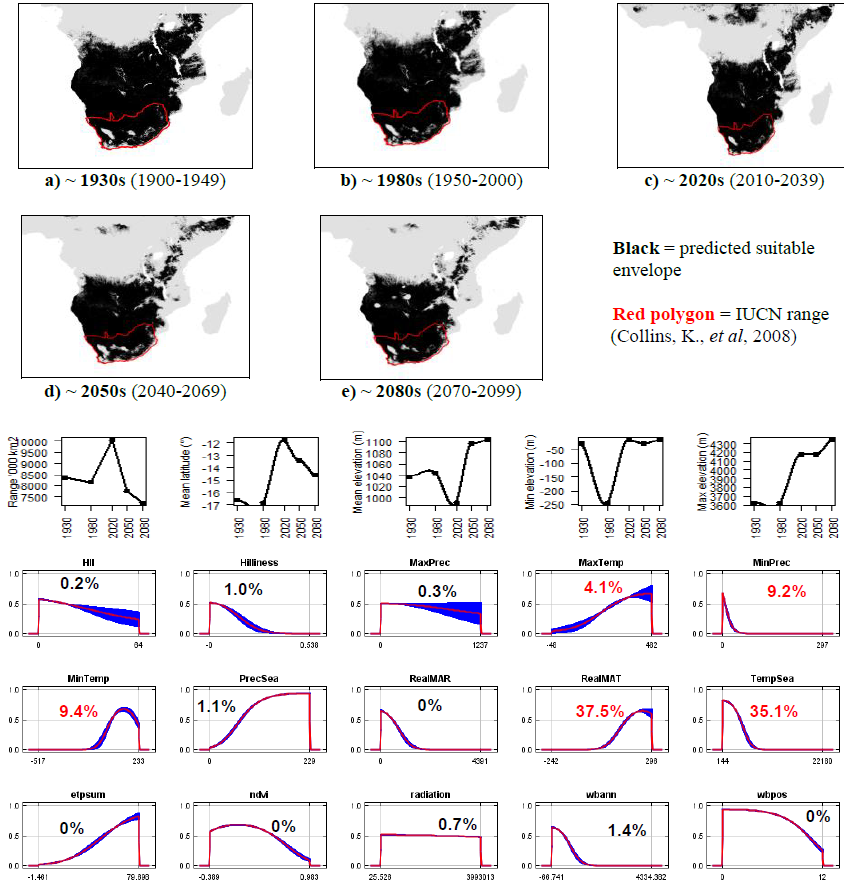

**#29 - Chinese hare** (*Lepus sinensis*) *n* = 141

**Expert:** Weihe Yang, Institute of Zoology, Chinese Academy of Sciences

**Expert evaluation:** Medium

**Data:** Modern and historic

**Envelope:** Climatic and habitat

**Dispersal distance:** 2.5km/year (Average for Asian leporids)

**Status:** MODELLABLE; **Included in final analysis:** √

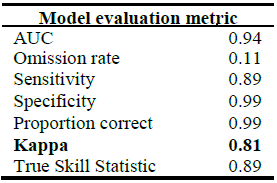

| Model evaluation metric |  |
| --- | --- |
| AUC | 0.94 |
| Omission rate | 0.11 |
| Sensitivity | 0.89 |
| Specificity | 0.99 |
| Proportion correct | 0.99 |
| **Kappa** | **0.81** |
| True Skill Statistic | 0.89 |

**Summary:** The Chinese hare’s bioclimatic envelope is predicted to increase by 60% with a ~2° mean latitudinal polewards shift and a mean increase in elevation of ~25m driven by an increase in maximum elevation. 95% of the permutation importance of the model was contributed to by mean annual temperature (56.5%), temperature seasonality (22.9%), precipitation seasonality (8.0%), mean annual precipitation (5.4%) and annual evapotranspiration (3.5%).

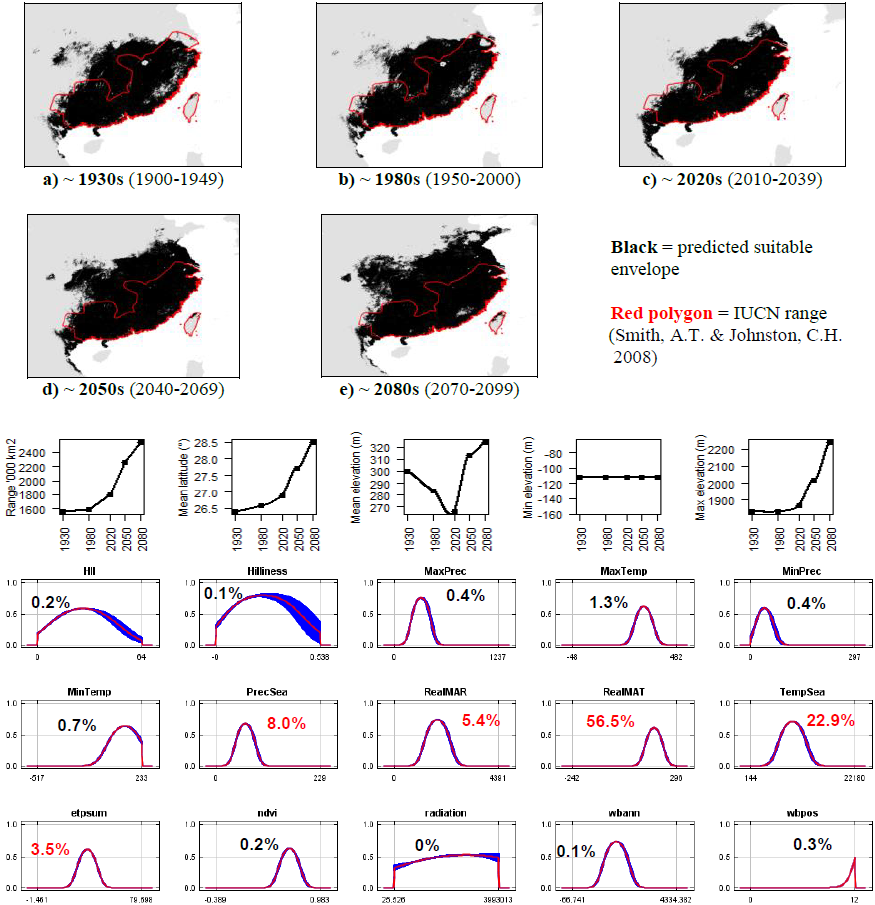

**#30 - Ethiopian highland hare** (*Lepus starcki*)

*n* = 13

**Expert:** Zelalem Tolesa, Addis Ababa University

**Expert evaluation:** Medium

**Data:** Modern and historic

**Envelope:** Climatic and habitat

**Dispersal distance:** 8km/year (Average for African leporids)

**Status:** MODELLABLE; **Included in final analysis:** √

| Model evaluation metric |  |
| --- | --- |
| AUC | 0.96 |
| Omission rate | 0.08 |
| Sensitivity | 0.92 |
| Specificity | 0.99 |
| Proportion correct | 0.99 |
| **Kappa** | **0.86** |
| True Skill Statistic | 0.92 |

**Summary:** The Ethiopian highland hare’s bioclimatic envelope is predicted to decrease by 90% with a ~7° mean latitudinal shift towards the Equator and a mean decrease in elevation of ~140m driven by a decrease in maximum elevation. 95% of the permutation importance of the model was contributed to by temperature seasonality (80.6%) and minimum temperature (18.5%).

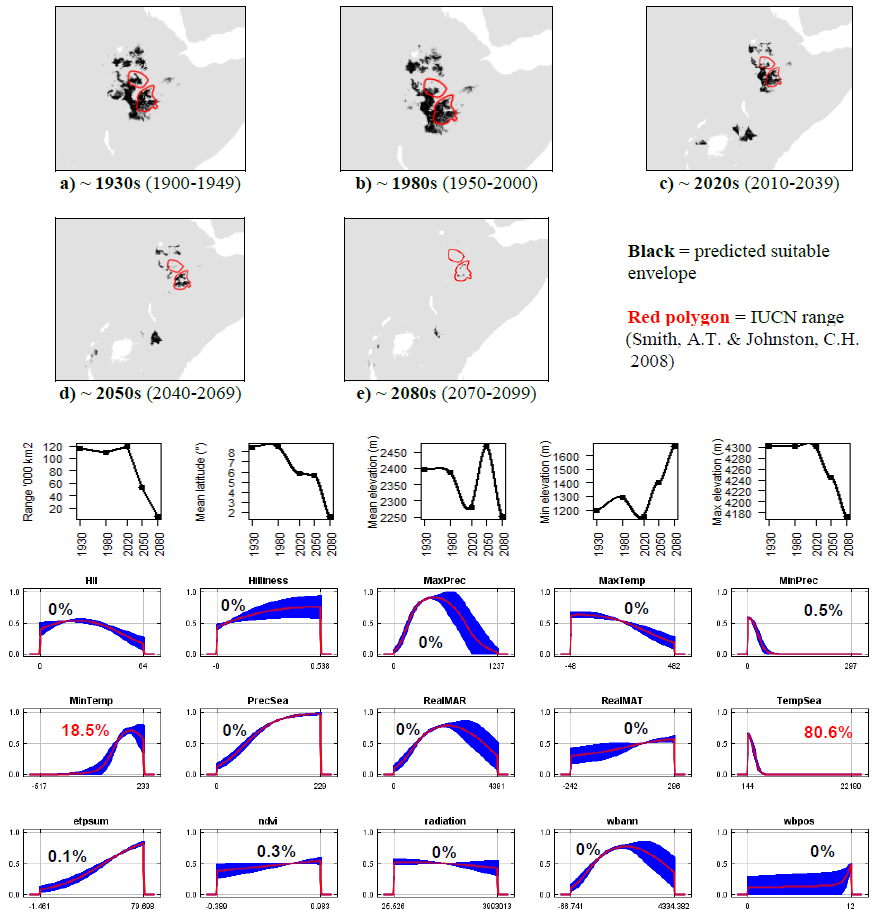

**#31 - Desert hare** (*Lepus tibetanus*) *n* = 55

**Expert:** Chelmala Srinivasulu, Osmania University, India

**Expert evaluation:** Medium

**Data:** Only modern

**Envelope:** Climatic and habitat

**Dispersal distance:** 2.5km/year (Average for Asian leporids)

**Status:** MODELLABLE; **Included in final analysis:** √

| Model evaluation metric |  |
| --- | --- |
| AUC | 0.92 |
| Omission rate | 0.15 |
| Sensitivity | 0.85 |
| Specificity | 0.99 |
| Proportion correct | 0.99 |
| **Kappa** | **0.57** |
| True Skill Statistic | 0.85 |

**Summary:** The Desert hare’s bioclimatic envelope is predicted to decrease by 50% with no latitudinal shift towards the Equator, but a mean increase in elevation of ~320m driven by an increase in maximum elevation. 95% of the permutation importance of the model was contributed to by minimum precipitation (54.5%), minimum temperature (19.3%), maximum precipitation (17.3%), annual evapotranspiration (2.8%) and human influence index (1.7%).

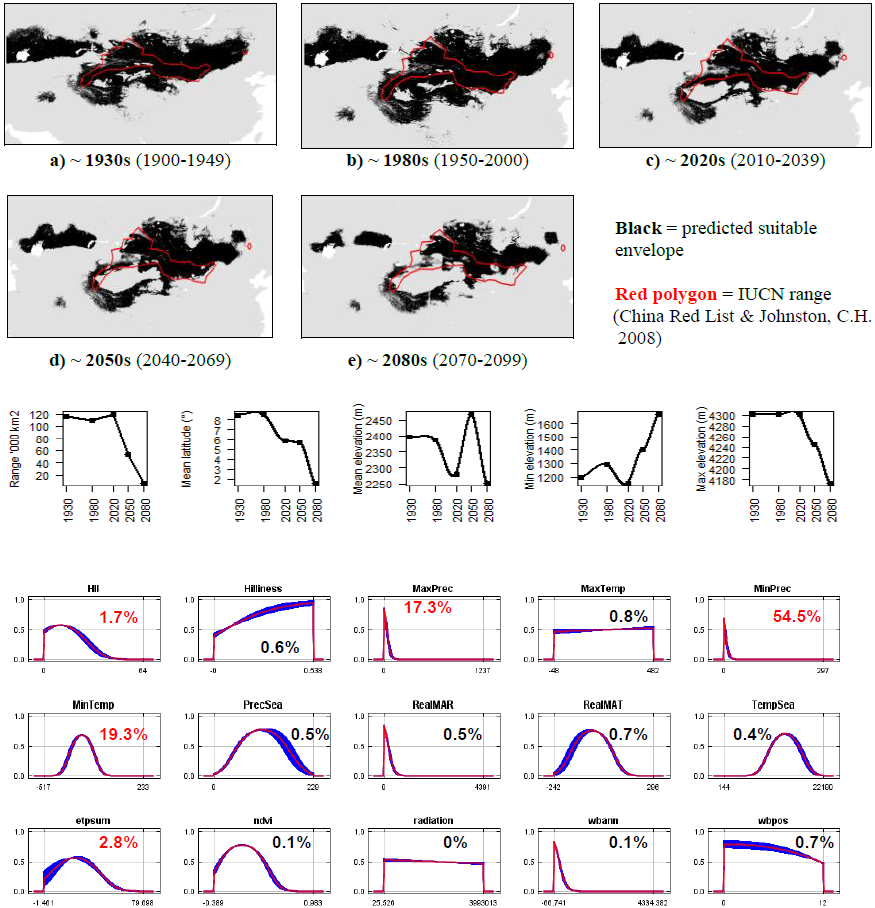

**#32 - Mountain hare** (*Lepus timidus*) - Eurasian populations *n* = 2,460

**Expert:** Neil Reid, Queen’s University Belfast

**Expert evaluation:** Medium

**Data:** Only modern

**Envelope:** Climatic and habitat

**Dispersal distance:** 2km/year (Expert)

**Status:** MODELLABLE; **Included in final analysis:** √

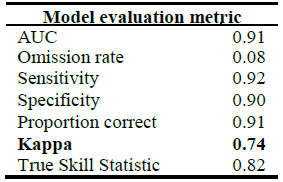

**Summary:** The Mountain hare’s bioclimatic envelope is predicted to decrease by 10% with a ~4° mean latitudinal polewards shift and a mean decrease in elevation of ~10m driven by a decrease in maximum elevation. 95% of the permutation importance of the model was contributed to by annual evapotranspiration (87.9%), temperature seasonality (3.9%), minimum precipitation (2.1%) and minimum temperature (1.6%).

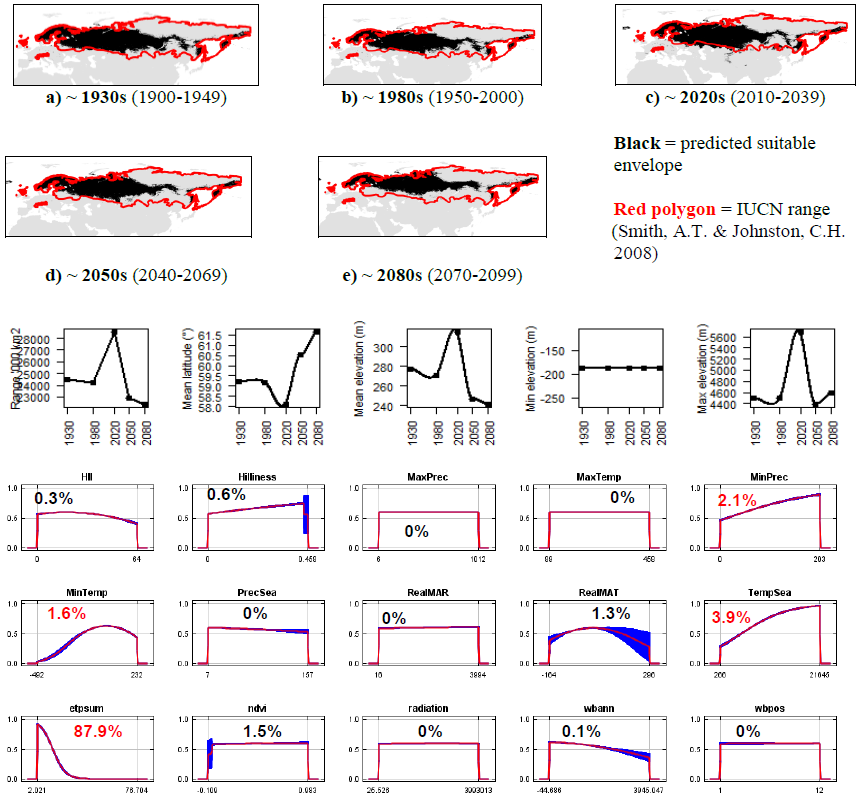

**#33 - Irish hare** (*Lepus timidus hibernicus*) *n* = 706

**Expert:** Neil Reid, Queen’s University Belfast

**Expert evaluation:** Medium

**Data:** Only modern

**Envelope:** Climatic and habitat

**Dispersal distance:** 2km/year (Expert)

**Status:** MODELLABLE; **Included in final analysis:** √

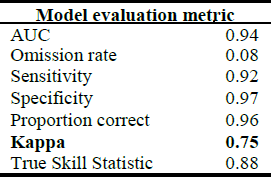

| Model evaluation metric |  |
| --- | --- |
| AUC | 0.94 |
| Omission rate | 0.08 |
| Sensitivity | 0.92 |
| Specificity | 0.97 |
| Proportion correct | 0.96 |
| **Kappa** | **0.75** |
| True Skill Statistic | 0.88 |

**Summary:** The Irish hare’s bioclimatic envelope is predicted to decrease by 50% with a ~0.5° mean latitudinal polewards shift and a mean decrease in elevation of ~10m driven by a decrease in maximum elevation. 95% of the permutation importance of the model was contributed to by temperature seasonality (44.6%), annual evapotranspiration (41.5%), normalised difference vegetation index (6.4%), precipitation seasonality (3.6%) and maximum precipitation (2.5%).

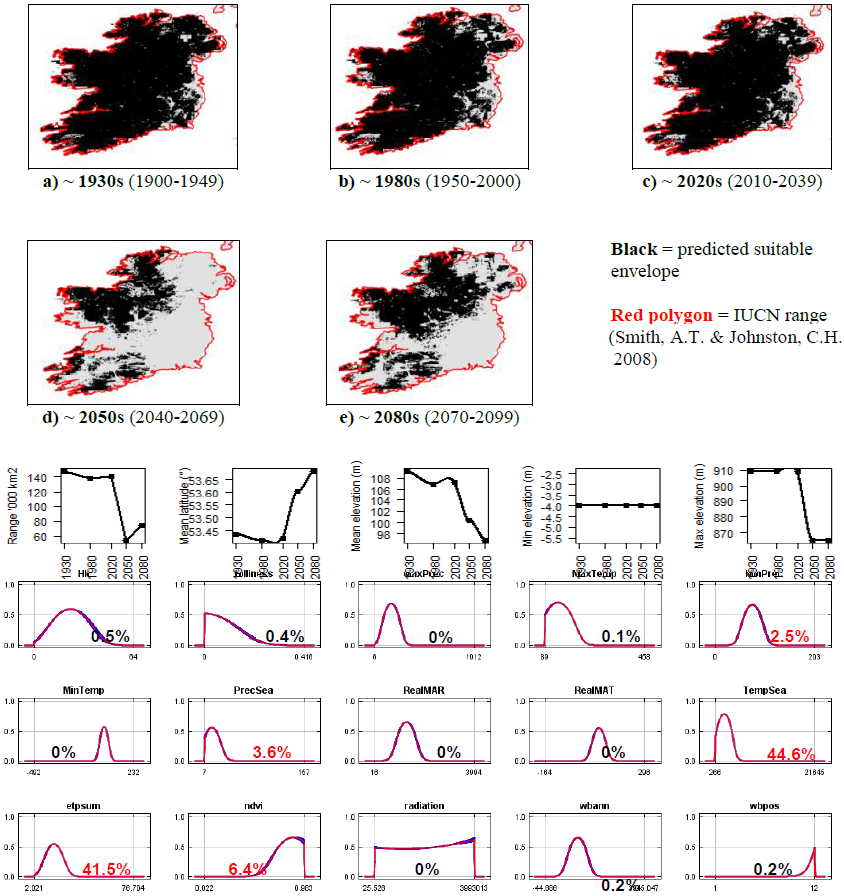

**#34 - Mountain hare** (*Lepus timidus*) - Eurasian & Irish populations combined *n* = 3,166

**Expert:** Neil Reid, Queen’s University Belfast

**Expert evaluation:** Medium

**Data:** Only modern

**Envelope:** Climatic and habitat

**Dispersal distance:** 2km/year (Expert)

**Status:** MODELLABLE; **Included in final analysis:** √

| Model evaluation metric |  |
| --- | --- |
| AUC | 0.92 |
| Omission rate | 0.07 |
| Sensitivity | 0.93 |
| Specificity | 0.91 |
| Proportion correct | 0.91 |
| **Kappa** | **0.78** |
| True Skill Statistic | 0.84 |

**Summary:** The Mountain hare’s bioclimatic envelope is predicted to decrease by 10% with a ~2° mean latitudinal polewards shift and a mean decrease in elevation of ~40m driven by a decrease in maximum elevation. 95% of the permutation importance of the model was contributed to by annual evapotranspiration (87.6%), temperature seasonality (4.1%), minimum precipitation (2.1%) and minimum temperature (1.6%).

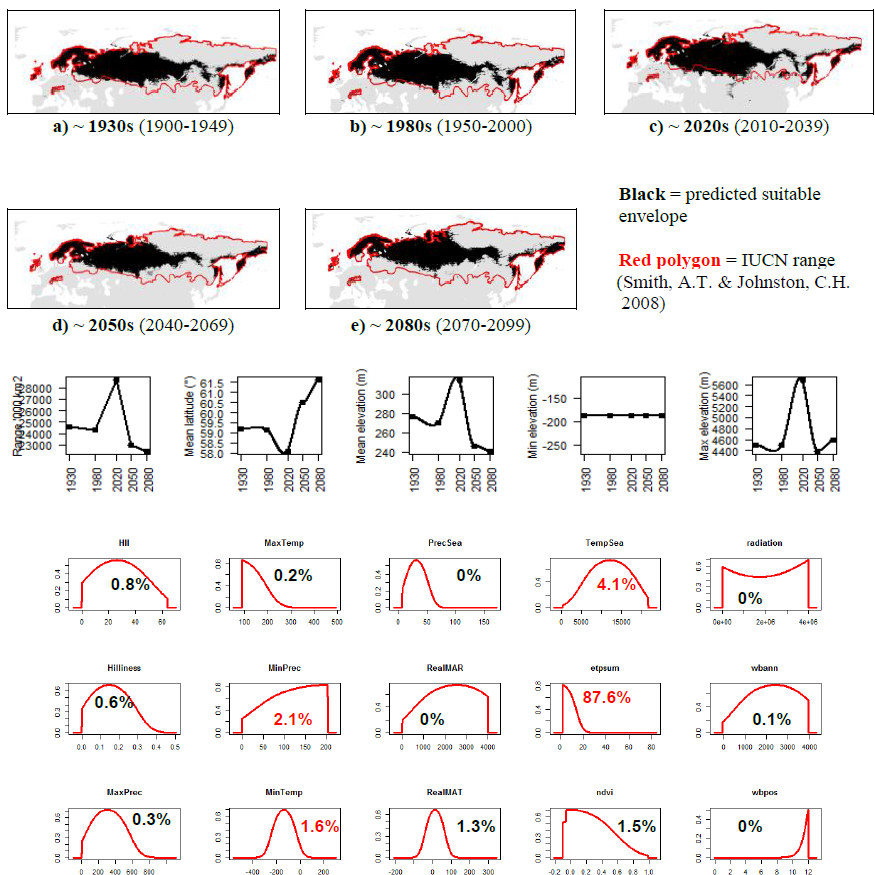

**#35 - Tolai hare** (*Lepus tolai*) *n* = 316

**Expert:** Chelmala Srinivasulu, Osmania University, India

**Expert evaluation:** Medium

**Data:** Only modern

**Envelope:** Climatic and habitat

**Dispersal distance:** 2.5km/year (Average for Asian leporids)

**Status:** MODELLABLE; **Included in final analysis:** √

| Model evaluation metric |  |
| --- | --- |
| AUC | 0.94 |
| Omission rate | 0.11 |
| Sensitivity | 0.89 |
| Specificity | 0.99 |
| Proportion correct | 0.98 |
| **Kappa** | **0.76** |
| True Skill Statistic | 0.88 |

**Summary:** The Tolai hare’s bioclimatic envelope is predicted to increase by 70% with a ~3° mean latitudinal polewards shift and a mean increase in elevation of ~280m driven by an increase in maximum elevation. 95% of the permutation importance of the model was contributed to by temperature seasonality (43.5%), maximum temperature (17.3%), mean annual temperature (15.2%), precipitation seasonality (9.6%), annual evapotranspiration (4.6%), mean annual precipitation (3.7%) and minimum precipitation (1.8%).

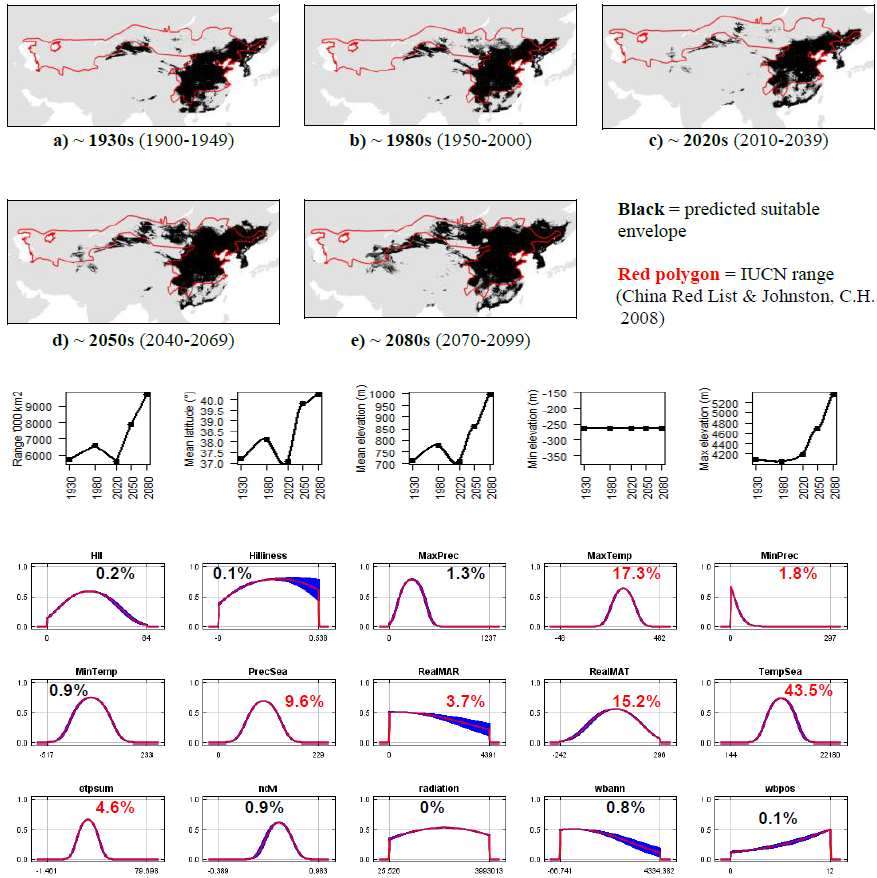

**#36 - White-tailed jackrabbit** (*Lepus townsendii*) *n* = 275

**Expert:** Eric Waltari, City University of New York

**Expert evaluation:** Medium

**Data:** Only modern

**Envelope:** Climatic and habitat

**Dispersal distance:** 18.9km/year (Average for NA leporids)

**Status:** MODELLABLE; **Included in final analysis:** √

| Model evaluation metric |  |
| --- | --- |
| AUC | 0.94 |
| Omission rate | 0.10 |
| Sensitivity | 0.90 |
| Specificity | 0.99 |
| Proportion correct | 0.99 |
| **Kappa** | **0.76** |
| True Skill Statistic | 0.89 |

**Summary:** The White-tailed jackrabbit’s bioclimatic envelope is predicted to decrease by 10% with a ~4° mean latitudinal polewards shift and a mean increase in elevation of ~200m driven by an increase in minimum elevation. 95% of the permutation importance of the model was contributed to by mean annual temperature (62.5%), maximum temperature (28.5%), temperature seasonality (3.4%) and annual evapotranspiration (1.6%).

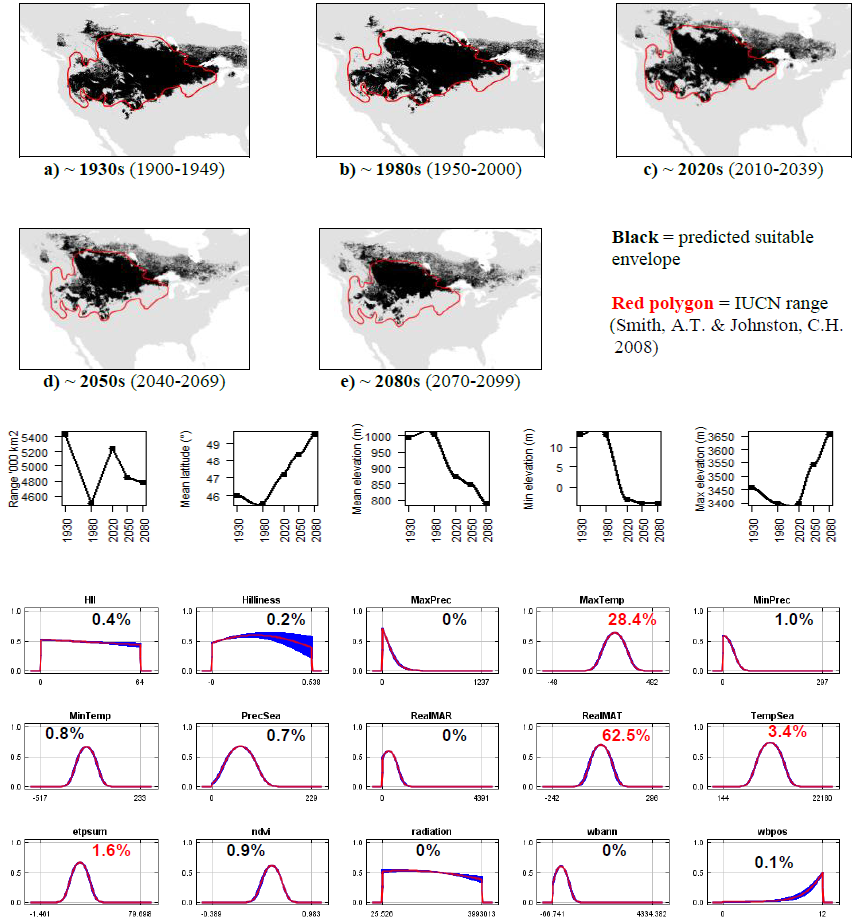

**#37 - Yarkand hare** (*Lepus yarkandensis*) *n* = 49

**Expert:** Weihe Yang, Institute of Zoology, Chinese Academy of Sciences

**Expert evaluation:** Medium

**Data:** Modern and historic

**Envelope:** Climatic and habitat

**Dispersal distance:** 2km/year (Smith & Xie, 2008)

**Status:** MODELLABLE; **Included in final analysis:** √

| Model evaluation metric |  |
| --- | --- |
| AUC | 0.95 |
| Omission rate | 0.10 |
| Sensitivity | 0.90 |
| Specificity | 0.99 |
| Proportion correct | 0.99 |
| **Kappa** | **0.74** |
| True Skill Statistic | 0.90 |

**Summary:** The Yarkand hare’s bioclimatic envelope is predicted to decrease by 100% with a ~1° mean latitudinal shift towards the Equator and a mean increase in elevation of ~1000m driven by an increase in minimum elevation. 95% of the permutation importance of the model was contributed to by minimum precipitation (58.8%), maximum precipitation (16.3%), mean annual precipitation (11.9%) and minimum temperature (11.7%).

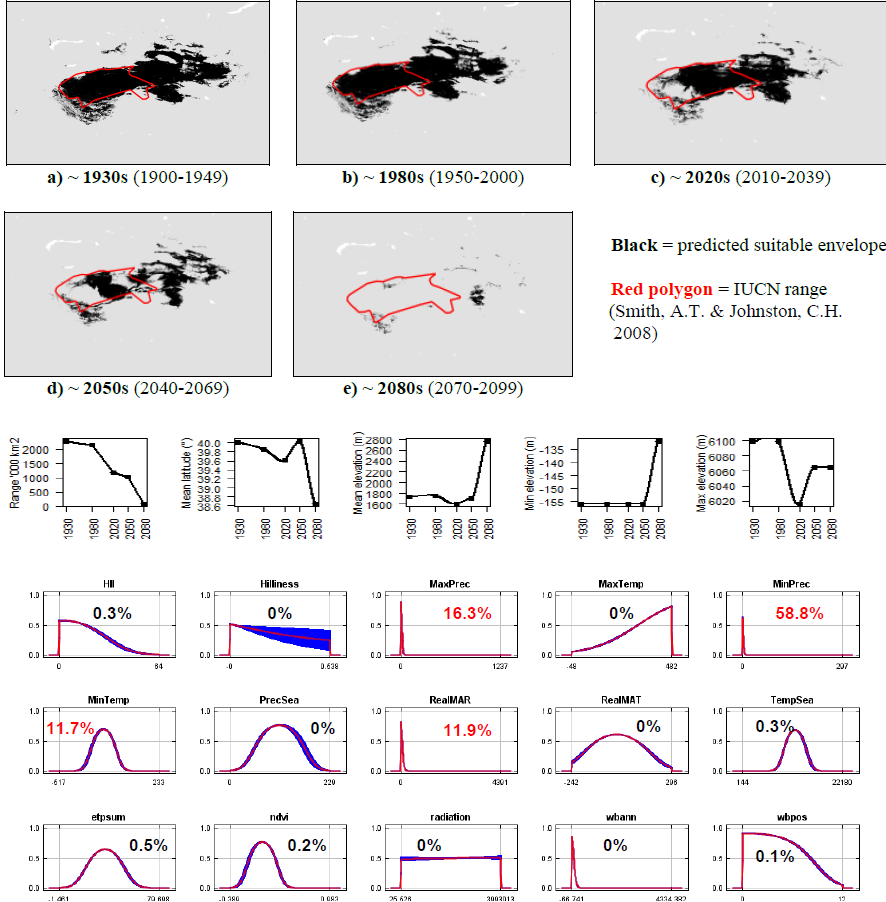

**#38 - Sumatran striped rabbit** (*Nesolagus netscheri*) *n* = 11

**Expert:** Hariyo Wibisono, Wildlife Conservation Society,

Indonesia

**Expert evaluation:** Poor

**Data:** Modern and historic

**Envelope:** Climatic and habitat

**Dispersal distance:** 0.01km/year (Expert)

**Status:** UNMODELLABLE; **Included in final analysis:** X

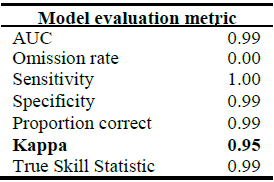

**Summary:** The Sumatran striped rabbit’s bioclimatic envelope is predicted to decrease by 91% with a ~1° mean latitudinal shift towards the Equator and a mean increase in elevation of ~330m driven by an increase in minimum elevation. 95% of the permutation importance of the model was contributed to by temperature seasonality (99.3%).

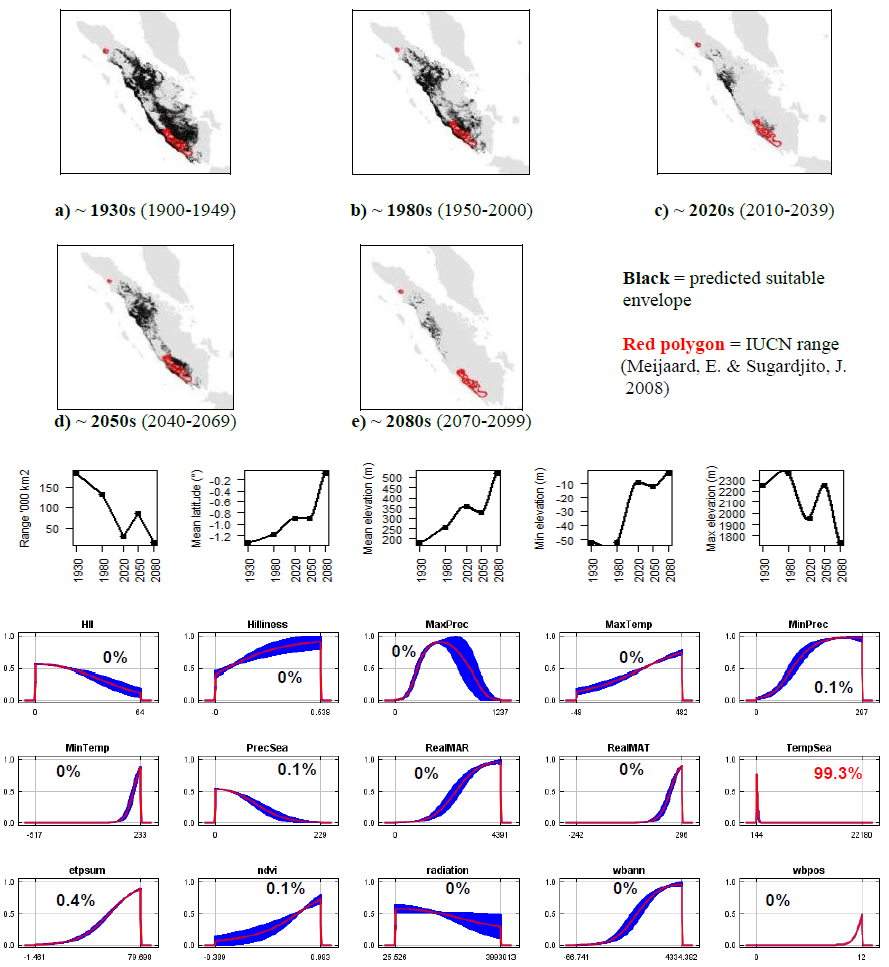

**#39 - Annamite striped rabbit** (*Nesolagus timminsi*) *n* = 4

**Expert:** Thomas Gray, WWF Greater Mekong & Andrew Tilker, University of Texas Austin

**Expert evaluation:** Poor

**Data:** Only modern

**Envelope:** Climatic and habitat

**Dispersal distance:** 10km/year (Expert)

**Status:** UNMODELLABLE; **Included in final analysis:** X

| Model evaluation metric |  |
| --- | --- |
| AUC | 0.99 |
| Omission rate | 0.00 |
| Sensitivity | 1.00 |
| Specificity | 0.99 |
| Proportion correct | 0.99 |
| **Kappa** | **0.50** |
| True Skill Statistic | 0.99 |

**Summary:** The Annamite striped rabbit’s bioclimatic envelope is predicted to decrease by 1500% with a ~0.3° mean latitudinal shift towards the Equator and a mean decrease in elevation of ~20m driven by an decrease in minimum elevation. 95% of the permutation importance of the model was contributed to by solar radiation (43.3%), minimum temperature (30.0%), mean annual temperature (14.1%), human influence index (6.4%) and annual evapotranspiration (2.3%).

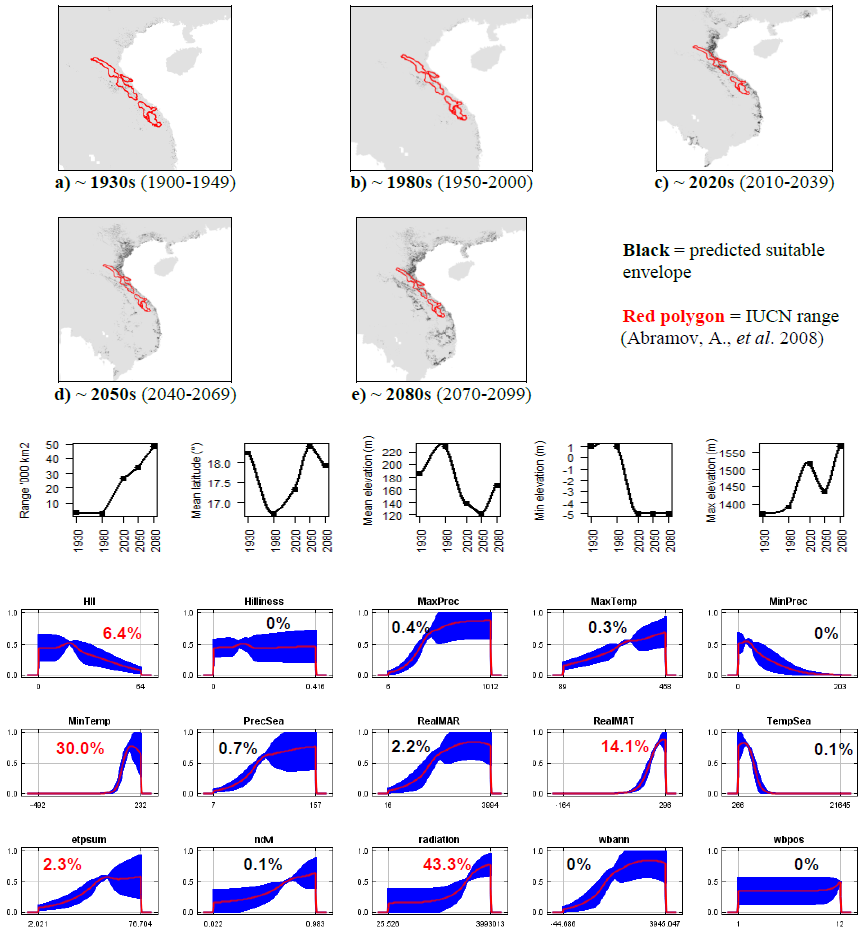

**#40 - Alpine pika** (*Ochotona alpina*) *n* = 16

**Expert:** Sumiya Ganzorig, Hokkaido University

**Expert evaluation:** Poor

**Data:** Modern and historic

**Envelope:** Climatic and habitat

**Dispersal distance:** 10km/year (Similar ecology to *O.pallasi*)

**Status:** UNMODELLABLE; **Included in final analysis:** X

| Model evaluation metric |  |
| --- | --- |
| AUC | 0.99 |
| Omission rate | 0.00 |
| Sensitivity | 1.00 |
| Specificity | 0.99 |
| Proportion correct | 0.99 |
| **Kappa** | **0.29** |
| True Skill Statistic | 0.99 |

**Summary:** The Alpine pika’s bioclimatic envelope is predicted to decrease by 10% with a ~4° mean latitudinal polewards shift and a mean decrease in elevation of ~80m. 95% of the permutation importance of the model was contributed to by minimum precipitation (50.1%), mean annual temperature (31.1%), number of months with a positive water balance (10.9%), minimum temperature (2.3%) and annual evapotranspiration (1.3%).

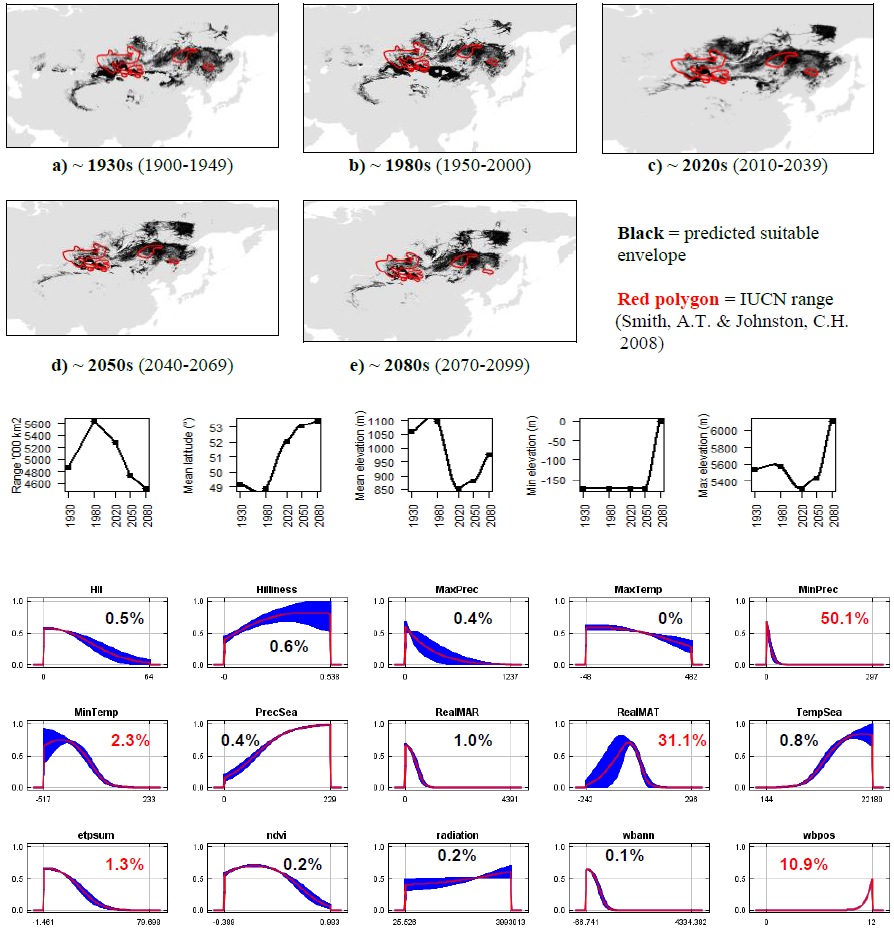

**#41 - Silver pika** (*Ochotona argentata*) *n* = 4

**Expert:** Andrew Smith, Arizona State University

**Expert evaluation:** Poor

**Data:** Only modern

**Envelope:** Climatic and habitat

**Dispersal distance:** 3km/year (Average for Asian pikas)

**Status:** UNMODELLABLE; **Included in final analysis:** X

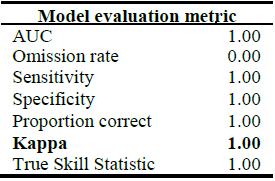

**Summary:** The Silver pika’s bioclimatic envelope is predicted to decrease by 80% with a ~4° mean latitudinal polewards shift and a mean increase in elevation of ~360m driven by an increase in minimum elevation. 95% of the permutation importance of the model was contributed to by minimum precipitation (62.7%), human influence index (17.1%), mean annual precipitation (8.6%), minimum temperature (5.4%) and annual evapotranspiration (1.8%).

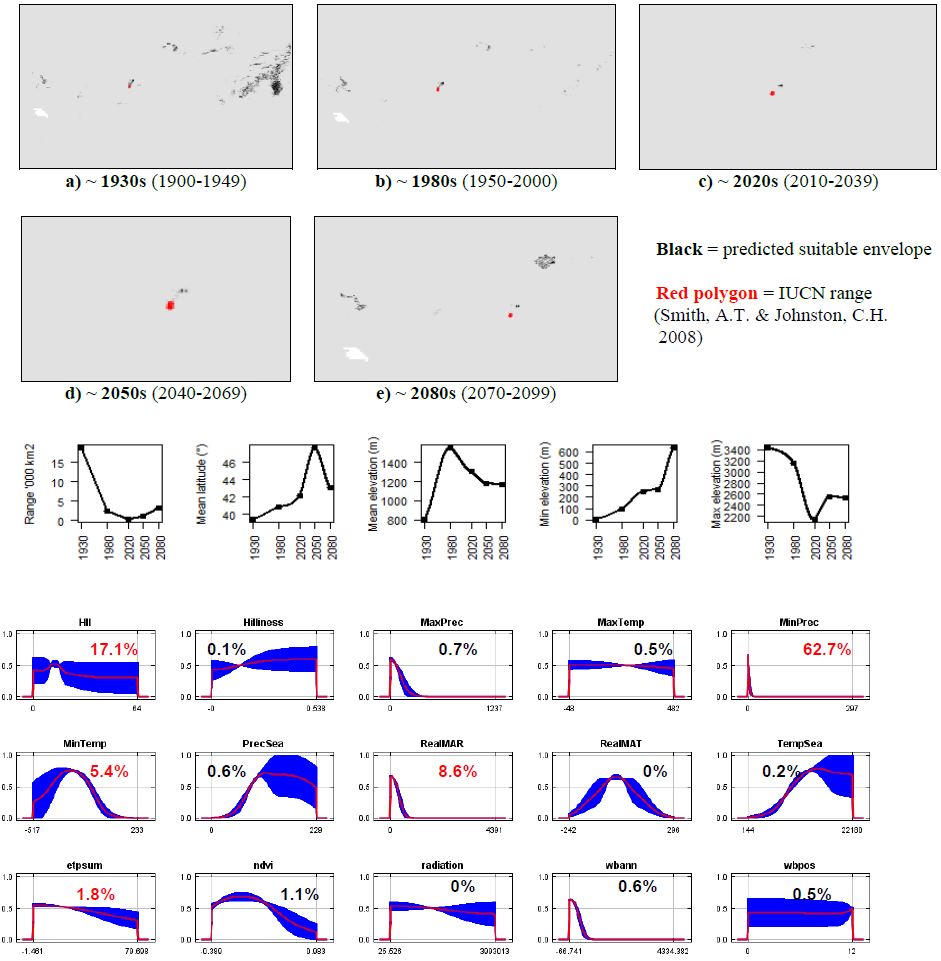

**#42 - Gansu pika** (*Ochotona cansus*) *n* = 38

**Expert:** Andrew Smith, Arizona State University

**Expert evaluation:** Medium

**Data:** Modern and historic

**Envelope:** Climatic and habitat

**Dispersal distance:** 1.5km/year (Similar ecology to *O.roylei*)

**Status:** MODELLABLE; **Included in final analysis:** √

| Model evaluation metric |  |
| --- | --- |
| AUC | 0.95 |
| Omission rate | 0.11 |
| Sensitivity | 0.89 |
| Specificity | 0.99 |
| Proportion correct | 0.99 |
| **Kappa** | **0.61** |
| True Skill Statistic | 0.89 |

**Summary:** The Gansu pika’s bioclimatic envelope is predicted to decrease by 60% with a ~0.4° mean latitudinal shift towards the Equator and a mean increase in elevation of ~230m driven by an increase in minimum elevation. 95% of the permutation importance of the model was contributed to by minimum precipitation (89.8%) and minimum temperature (6.9%).

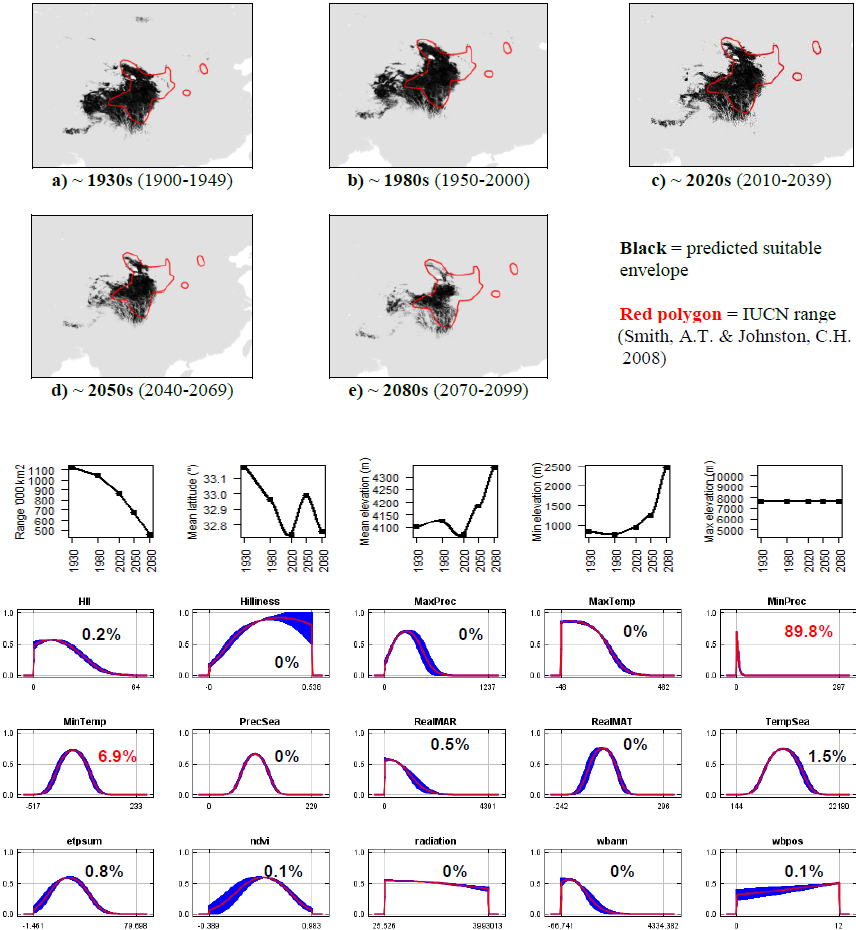

**#43 - Collared pika** (*Ochotona collaris*) *n* = 193

**Expert:** Hayley Lanier, University of Michigan & David Hik,

University of Alberta

**Expert evaluation:** Poor

**Data:** Modern and historic

**Envelope:** Climatic and habitat

**Dispersal distance:** 1km/year (Expert)

**Status:** UNMODELLABLE; **Included in final analysis:** X

| Model evaluation metric |  |
| --- | --- |
| AUC | 0.95 |
| Omission rate | 0.10 |
| Sensitivity | 0.90 |
| Specificity | 0.99 |
| Proportion correct | 0.99 |
| **Kappa** | **0.86** |
| True Skill Statistic | 0.90 |

**Summary:** The Collared pika’s bioclimatic envelope is predicted to increase by 20% with a ~2° mean latitudinal polewards shift and a mean decrease in elevation of ~140m. 95% of the permutation importance of the model was contributed to by annual evapotranspiration (86.7%), mean annual temperature (3.3%), normalised difference vegetation index (3.2%) and maximum temperature (2.0%).

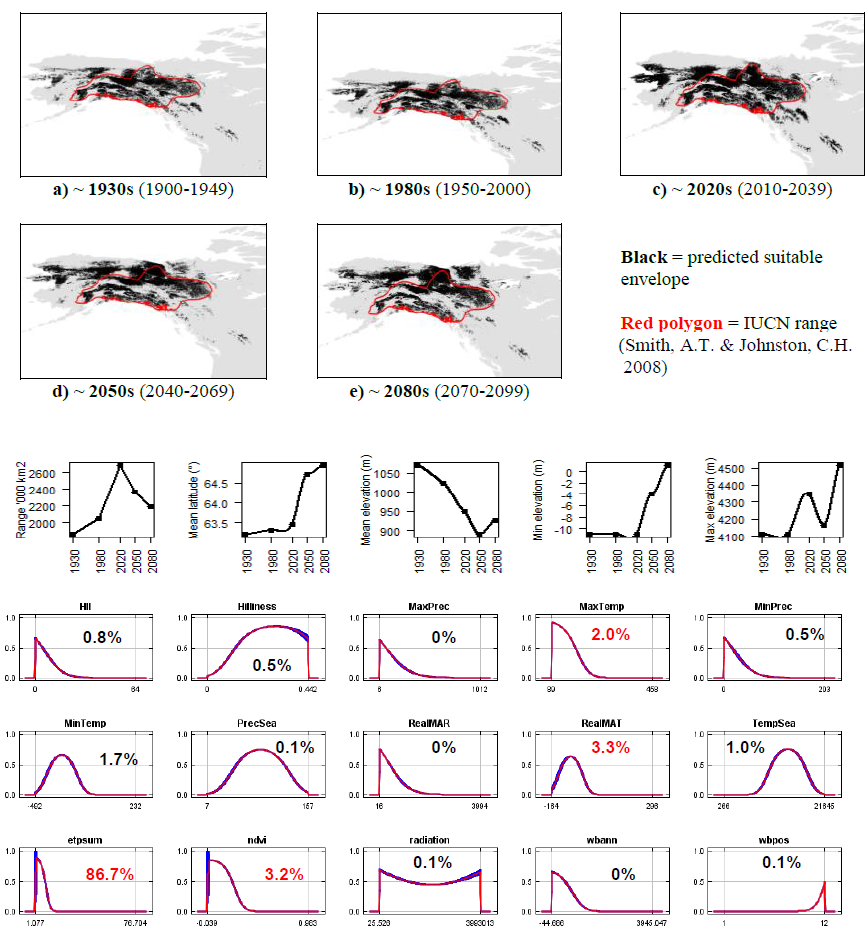

**#44 - Plateau pika** (*Ochotona curzoniae*) *n* = 131

**Expert:** Andrew Smith, Arizona State University

**Expert evaluation:** Good

**Data:** Only modern

**Envelope:** Climatic and habitat

**Dispersal distance:** 0.1km/year (Expert)

**Status:** MODELLABLE; **Included in final analysis:** √

| Model evaluation metric |  |
| --- | --- |
| AUC | 0.94 |
| Omission rate | 0.11 |
| Sensitivity | 0.89 |
| Specificity | 0.99 |
| Proportion correct | 0.99 |
| **Kappa** | **0.76** |
| True Skill Statistic | 0.88 |

**Summary:** The Plateau pika’s bioclimatic envelope is predicted to decrease by 30% with a ~1° mean latitudinal shift towards the Equator and a mean increase in elevation of ~700m driven by an increase in minimum and maximum elevation. 95% of the permutation importance of the model was contributed to by minimum precipitation (48.0%), maximum temperature (42.0%) and annual evapotranspiration (6.8%).

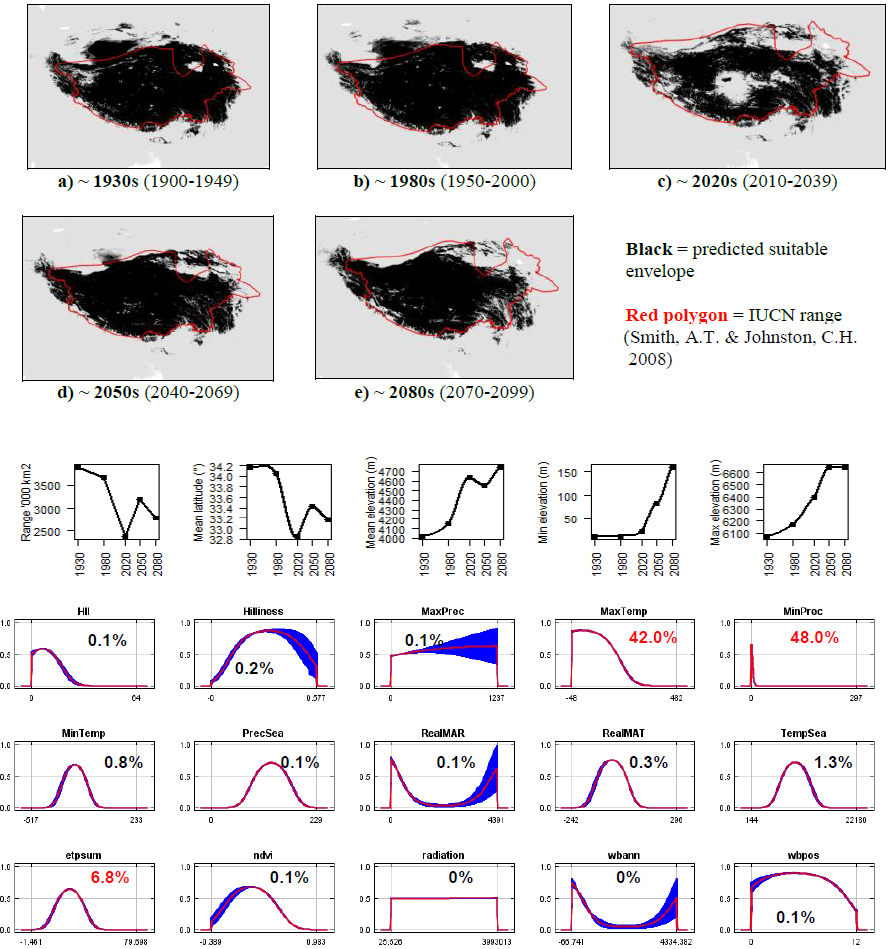

**#45 – Daurian pika** (*Ochotona dauurica*) *n* = 131

**Expert:** Andrew Smith, Arizona State University

**Expert evaluation:** Medium

**Data:** Only modern

**Envelope:** Climatic and habitat

**Dispersal distance:** 0.1km/year (Similar ecology to *O.curzoniae*)

**Status:** MODELLABLE; **Included in final analysis:** √

| Model evaluation metric |  |
| --- | --- |
| AUC | 0.95 |
| Omission rate | 0.10 |
| Sensitivity | 0.90 |
| Specificity | 0.99 |
| Proportion correct | 0.99 |
| **Kappa** | **0.66** |
| True Skill Statistic | 0.89 |

**Summary:** The Daurian pika’s bioclimatic envelope is predicted to decrease by 25% with a ~1° mean latitudinal polewards shift and a mean decrease in elevation of ~60m. 95% of the permutation importance of the model was contributed to by minimum precipitation (92.3%) and number of months with a positive water balance (2.7%).

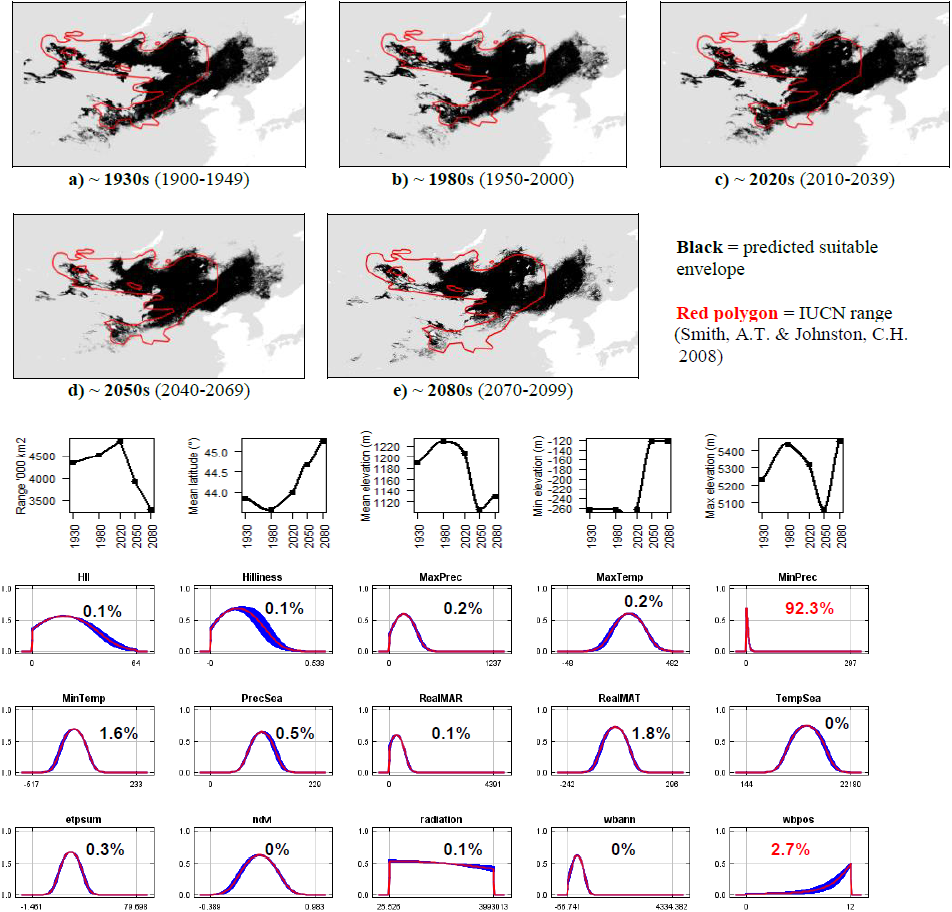

**#46 - Chinese red pika** (*Ochotona erythrotis*) *n* = 39

**Expert:** Andrew Smith, Arizona State University

**Expert evaluation:** Poor

**Data:** Modern and historic

**Envelope:** Climatic and habitat

**Dispersal distance:** 3km/year (Average for Asian pikas)

**Status:** UNMODELLABLE; **Included in final analysis:** X

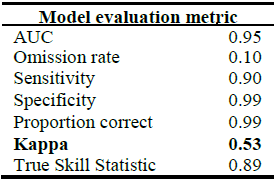

**Summary:** The Chinese red pika’s bioclimatic envelope is predicted to decrease by 20% with a ~3° mean latitudinal polewards shift and a mean decrease in elevation of ~2400m driven by a decrease in minimum elevation. 95% of the permutation importance of the model was contributed to by minimum precipitation (87.2%), minimum temperature (7.2%) and mean annual temperature (2.9%).

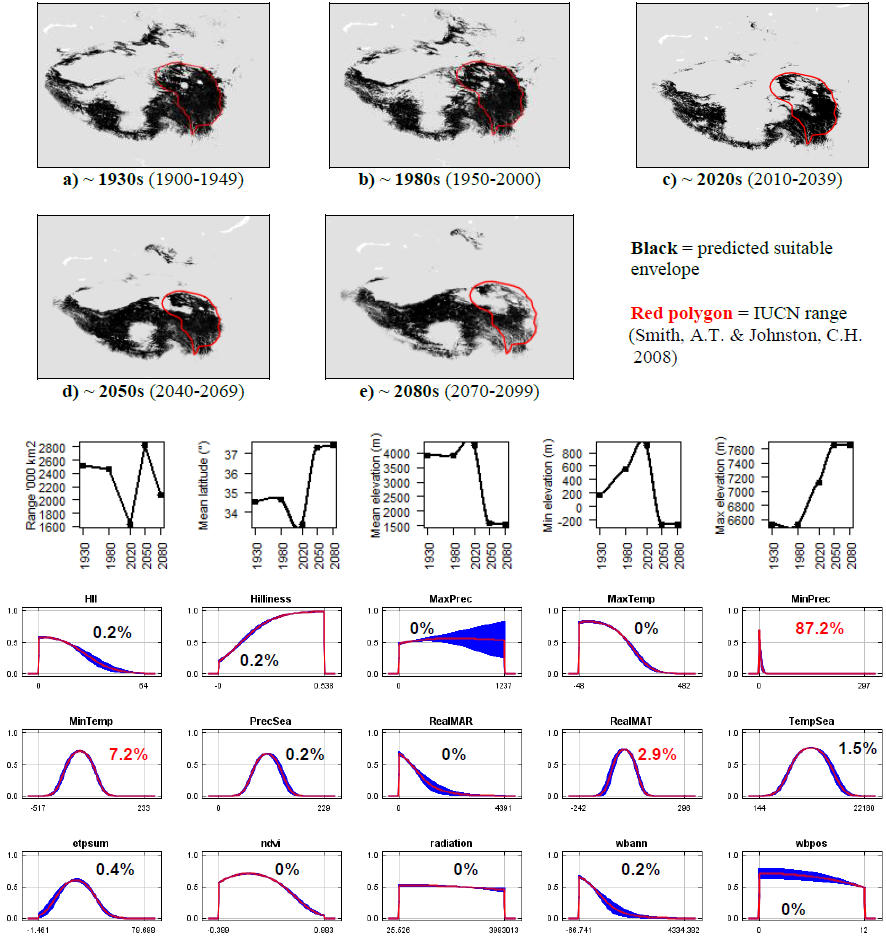

**#47 - Forrest’s pika** (*Ochotona forresti*) *n* = 9

**Expert:** Andrew Smith, Arizona State University

**Expert evaluation:** Poor

**Data:** Only modern

**Envelope:** Climatic and habitat

**Dispersal distance:** 3km/year (Average for Asian pikas)

**Status:** UNMODELLABLE; **Included in final analysis:** X

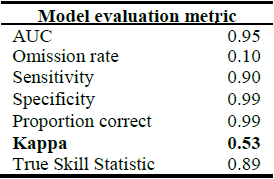

**Summary:** The Forrest’s pika’s bioclimatic envelope is predicted to decrease by 40% with a ~1° mean latitudinal polewards shift and a mean increase in elevation of ~600m driven by an increase in maximum elevation. 95% of the permutation importance of the model was contributed to by minimum precipitation (36.8%), temperature seasonality (20.6%), normalised difference vegetation index (12.1%), maximum temperature (9.3%), precipitation seasonality (7.6%), human influence index (6.2%) and surface roughness index (4.2%).

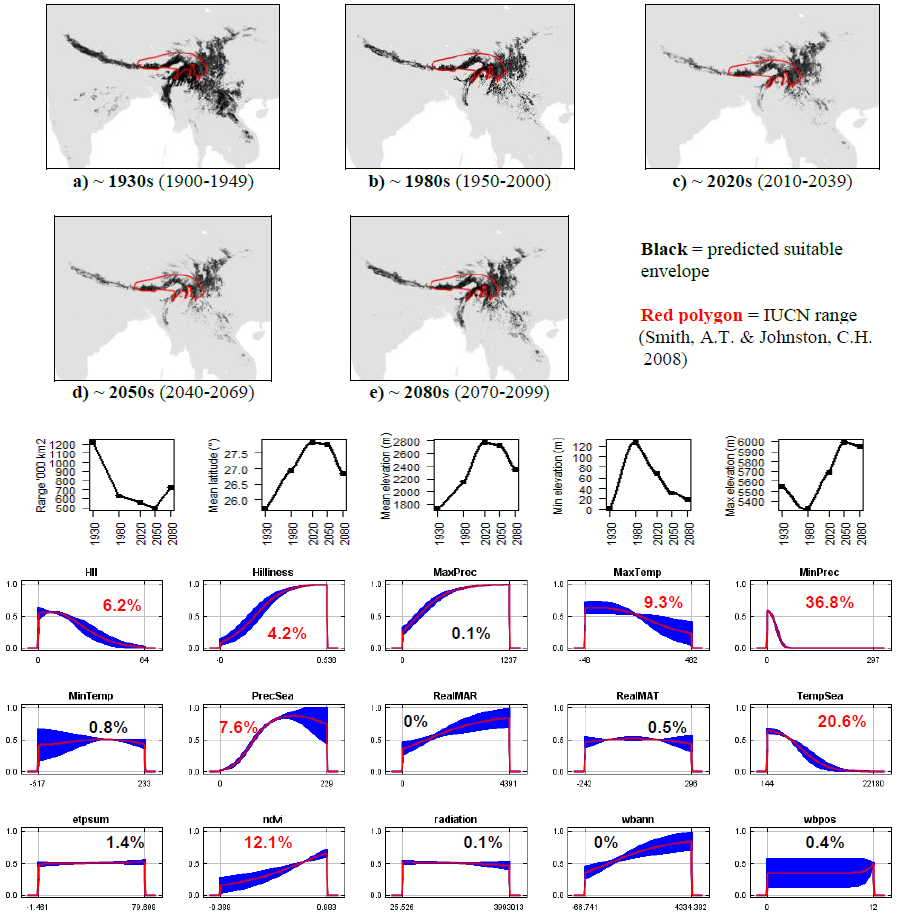

**#48 - Glover’s pika** (*Ochotona gloveri*) *n* = 22

**Expert:** Andrew Smith, Arizona State University

**Expert evaluation:** Medium

**Data:** Only modern

**Envelope:** Climatic and habitat

**Dispersal distance:** 3km/year (Average for Asian pikas)

**Status:** MODELLABLE; **Included in final analysis:** √

| Model evaluation metric |  |
| --- | --- |
| AUC | 0.99 |
| Omission rate | 0.00 |
| Sensitivity | 1.00 |
| Specificity | 0.99 |
| Proportion correct | 0.99 |
| **Kappa** | **0.65** |
| True Skill Statistic | 0.99 |

**Summary:** The Glover’s pika’s bioclimatic envelope is predicted to decrease by 50% with a ~0.5° mean latitudinal polewards shift and a mean increase in elevation of ~270m driven by an increase in minimum and maximum elevation. 95% of the permutation importance of the model was contributed to by minimum precipitation (46.3%), minimum temperature (28.8%), mean annual temperature (12.7%), human influence index (3.8%), temperature seasonality (3.1%) and precipitation seasonality (1.8%).

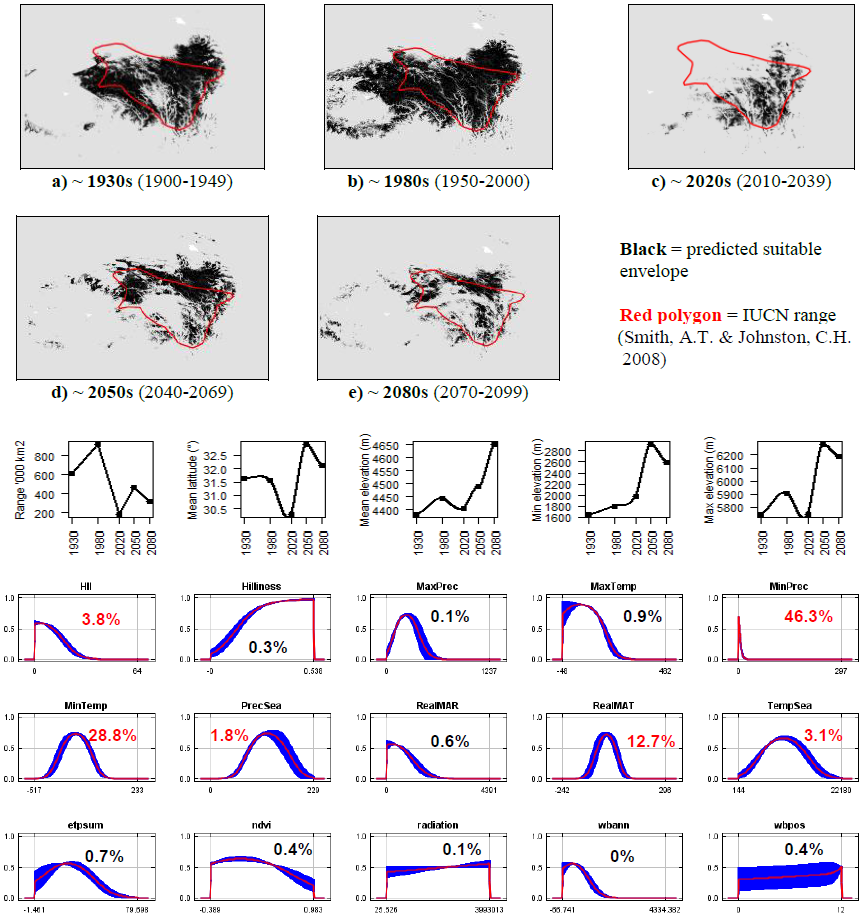

**#49 - Hoffmann’s pika** (*Ochotona hoffmanni*) *n* = 5

**Expert:** Andrey Lissovsky, Zoological Museum of Moscow State University

**Expert evaluation:** Medium

**Data:** Modern and historic

**Envelope:** Climatic and habitat

**Dispersal distance:** 3km/year (Average for Asian pikas)

**Status:** MODELLABLE; **Included in final analysis:** √

| Model evaluation metric |  |
| --- | --- |
| AUC | 0.99 |
| Omission rate | 0.00 |
| Sensitivity | 1.00 |
| Specificity | 0.99 |
| Proportion correct | 0.99 |
| **Kappa** | **0.91** |
| True Skill Statistic | 0.99 |

**Summary:** The Hoffmann’s pika’s bioclimatic envelope is predicted to decrease by 90% with a ~1° mean latitudinal shift towards the Equator and a mean increase in elevation of ~470m driven by an increase in minimum elevation. 95% of the permutation importance of the model was contributed to by precipitation seasonality (73.2%), mean annual temperature (12.6%), minimum temperature (6.5%) and minimum precipitation (5.3%).

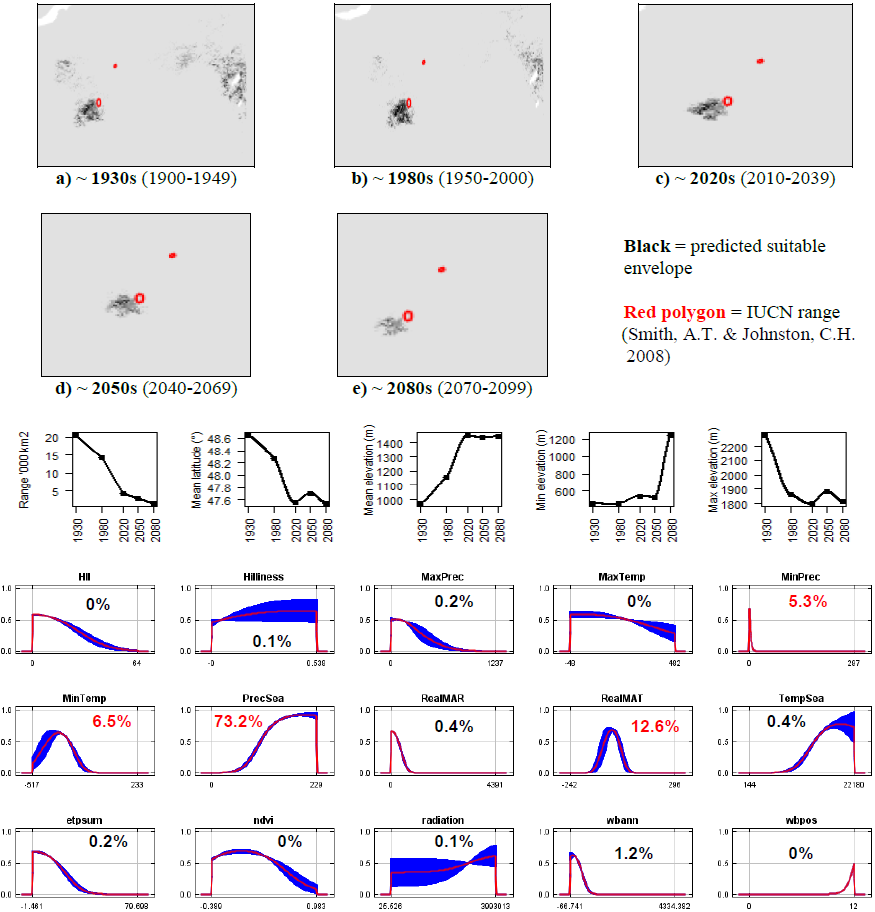

**#50 - Siberian pika** (*Ochotona hyperborea*) *n* = 16

**Expert:** Julia Witczuk, Warsaw Agricultural University,

Poland

**Expert evaluation:** Poor

**Data:** Modern and historic

**Envelope:** Climatic and habitat

**Dispersal distance:** 10km/year (Similar ecology to *O.alpina*)

**Status:** UNMODELLABLE; **Included in final analysis:** X

| Model evaluation metric |  |
| --- | --- |
| AUC | 0.97 |
| Omission rate | 0.06 |
| Sensitivity | 0.94 |
| Specificity | 0.99 |
| Proportion correct | 0.99 |
| **Kappa** | **0.40** |
| True Skill Statistic | 0.93 |

**Summary:** The Siberian pika’s bioclimatic envelope is predicted to decrease by 100% with a ~5° mean latitudinal polewards shift and a mean increase in elevation of ~200m driven by an increase in minimum and maximum elevation. 95% of the permutation importance of the model was contributed to by mean annual temperature (72.5%), annual evapotranspiration (10.4%), minimum precipitation (6.4%), human influence index (2.8%), normalised difference vegetation index (2.6%) and annual water balance (1.8%).

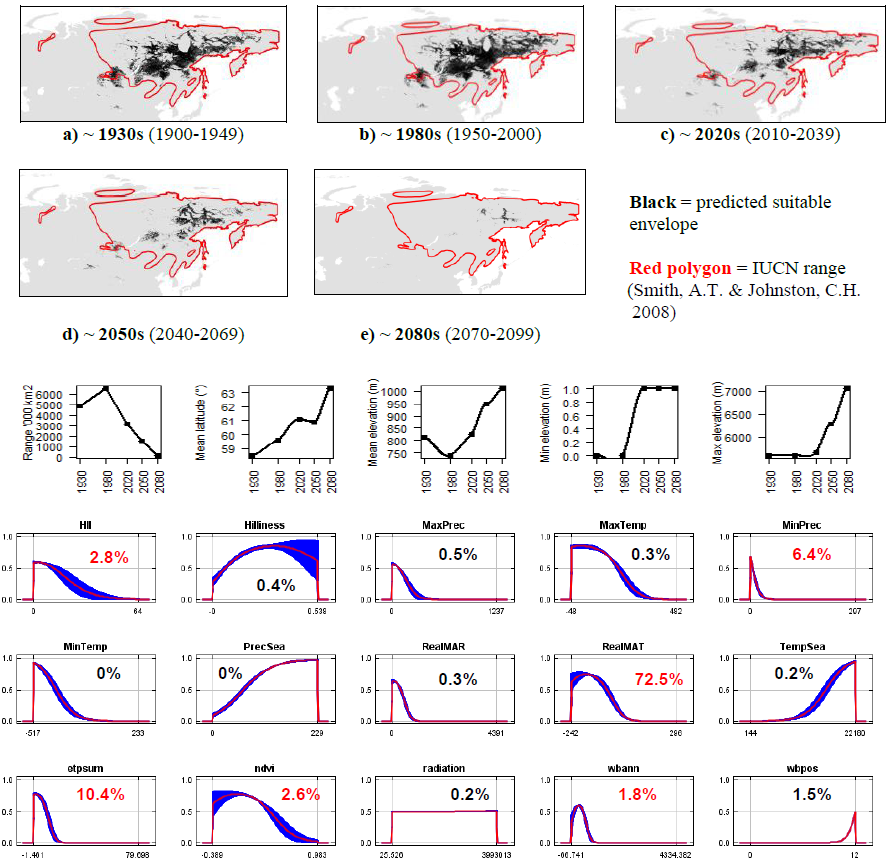

**#51- Ili pika** (*Ochotona iliensis*) *n* = 11

**Expert:** Andrew Smith, Arizona State University

**Expert evaluation:** Poor

**Data:** Only modern

**Envelope:** Climatic and habitat

**Dispersal distance:** 1km/year (Similar ecology to *O.koslowi*)

**Status:** UNMODELLABLE; **Included in final analysis:** X

| Model evaluation metric |  |
| --- | --- |
| AUC | 0.99 |
| Omission rate | 0.00 |
| Sensitivity | 1.00 |
| Specificity | 0.99 |
| Proportion correct | 0.99 |
| **Kappa** | **0.55** |
| True Skill Statistic | 0.99 |

**Summary:** The Ili pika’s bioclimatic envelope is predicted to decrease by 20% with a ~1° mean latitudinal polewards shift and a mean increase in elevation of ~80m. 95% of the permutation importance of the model was contributed to by minimum precipitation (52.7%), minimum temperature (22.2%), mean annual temperature (8.8%), maximum precipitation (6.5%), temperature seasonality (3.0%) and human influence index (2.2%).

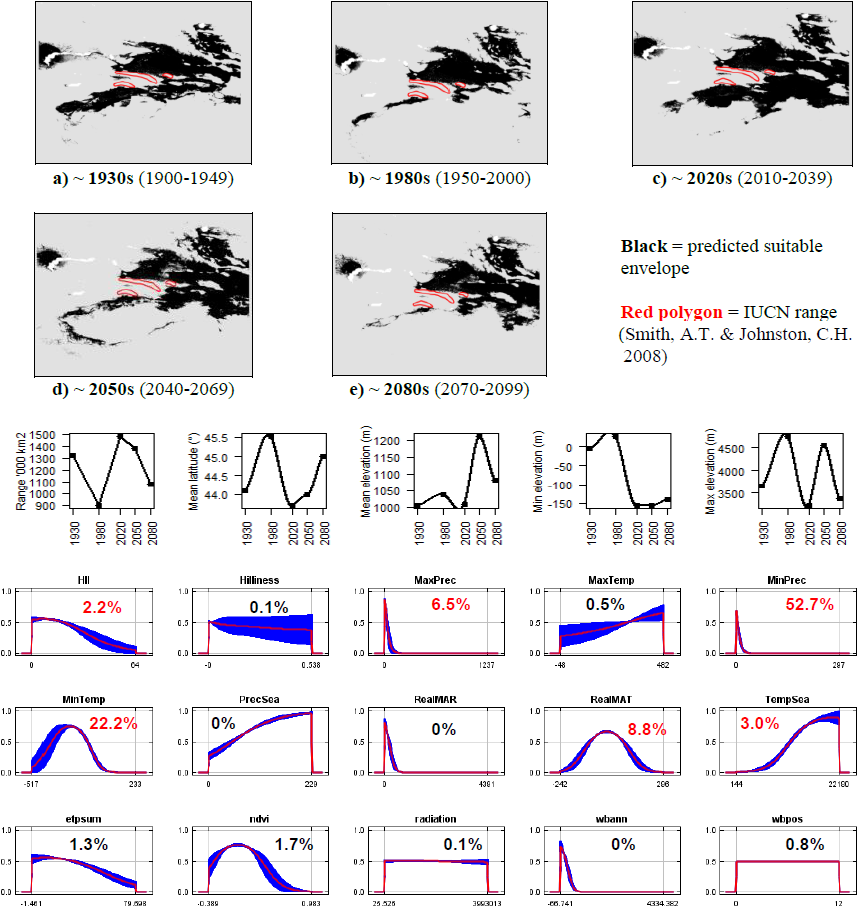

**#52 - Kozlov’s pika** (*Ochotona koslowi*) *n* = 5

**Expert:** Andrew Smith, Arizona State University

**Expert evaluation:** Medium

**Data:** Only modern

**Envelope:** Climatic and habitat

**Dispersal distance:** 1km/year (Expert)

**Status:** MODELLABLE; **Included in final analysis:** √

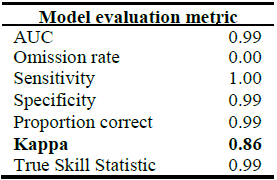

**Summary:** The Kozlov’s pika’s bioclimatic envelope is predicted to decrease by 100% (total extinction). 95% of the permutation importance of the model was contributed to by mean annual precipitation (66.8%), minimum precipitation (24.2%) and human influence index (5.8%).

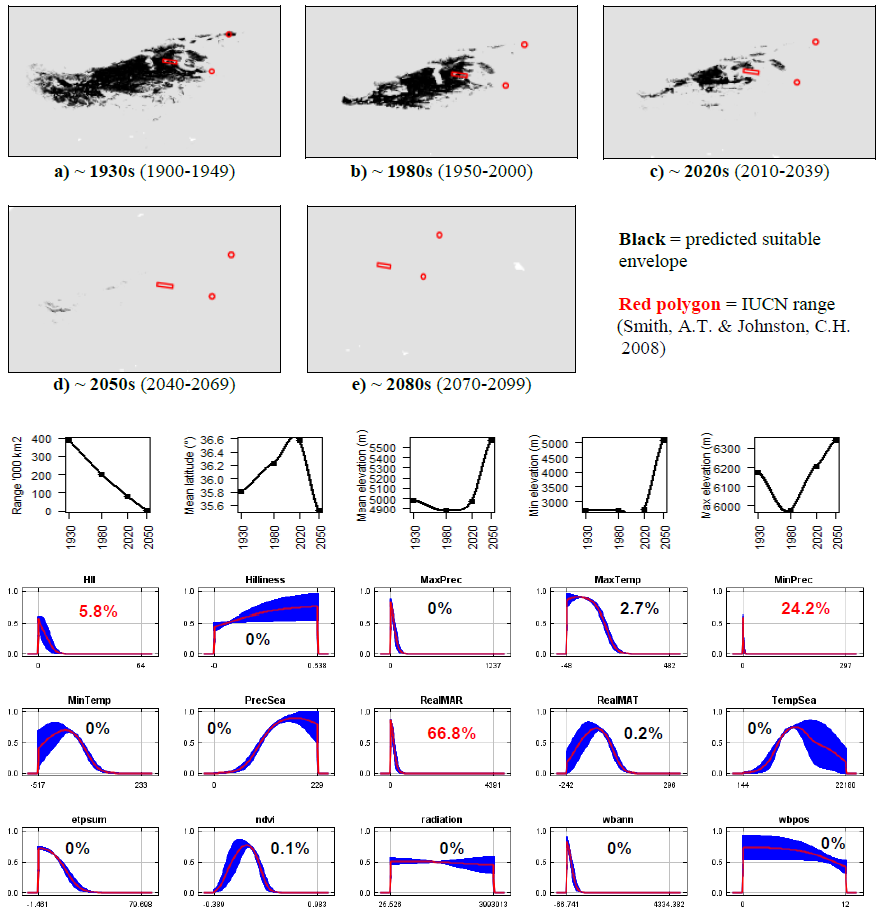

**#53 - Ladak pika** (*Ochotona ladacensis*) *n* = 18

**Expert:** Andrew Smith, Arizona State University

**Expert evaluation:** Medium

**Data:** Modern and historic

**Envelope:** Climatic and habitat

**Dispersal distance:** 0.05km/year (Similar ecology to *O.curzoniae*)

**Status:** MODELLABLE; **Included in final analysis:** √

| Model evaluation metric |  |
| --- | --- |
| AUC | 0.99 |
| Omission rate | 0.00 |
| Sensitivity | 1.00 |
| Specificity | 0.99 |
| Proportion correct | 0.99 |
| **Kappa** | **0.63** |
| True Skill Statistic | 0.99 |

**Summary:** The Ladak pika’s bioclimatic envelope is predicted to decrease by 70% with a ~1° mean latitudinal shift towards the Equator and a mean increase in elevation of ~1400m driven by an increase in minimum and maximum elevation. 95% of the permutation importance of the model was contributed to by mean annual precipitation (47.9%), minimum precipitation (41.7%) and minimum temperature (5.7%).

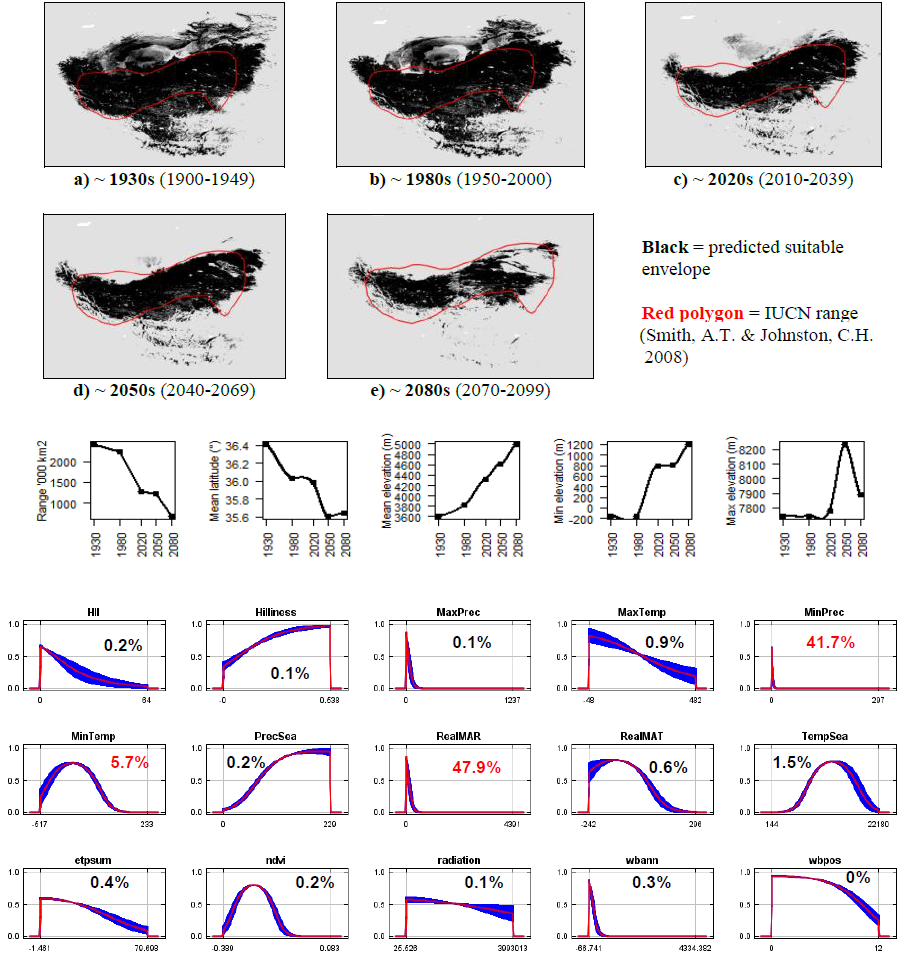

**#54 - Large-eared pika** (*Ochotona macrotis*) *n* = 49

**Expert:** Nishma Dahal, National Centre for Biological Sciences, India

**Expert evaluation:** Medium

**Data:** Modern and historic

**Envelope:** Climatic and habitat

**Dispersal distance:** 1km/year (Similar ecology to *O.roylei*)

**Status:** MODELLABLE; **Included in final analysis:** √

| Model evaluation metric |  |
| --- | --- |
| AUC | 0.94 |
| Omission rate | 0.10 |
| Sensitivity | 0.90 |
| Specificity | 0.99 |
| Proportion correct | 0.99 |
| **Kappa** | **0.43** |
| True Skill Statistic | 0.89 |

**Summary:** The Large-eared pika’s bioclimatic envelope is predicted to decrease by 40% with a ~1° mean latitudinal shift towards the Equator and a mean increase in elevation of ~880m driven by an increase in minimum elevation. 95% of the permutation importance of the model was contributed to by minimum precipitation (87.3%), minimum temperature (5.8%) and mean annual temperature (2.1%).

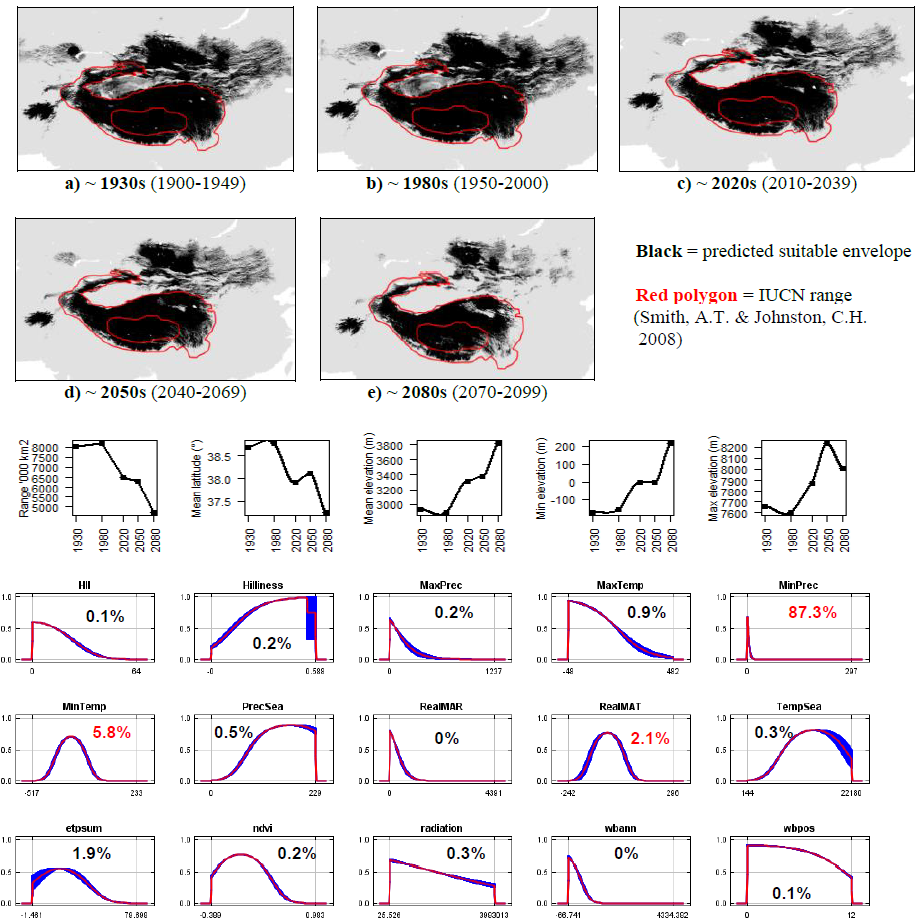

**#55 - Nubra’s pika** (*Ochotona nubrica*) *n* = 13

**Expert:** Nishma Dahal, National Centre for Biological

Sciences, India

**Expert evaluation:** Medium

**Data:** Only modern

**Envelope:** Climatic and habitat

**Dispersal distance:** 0.05km/year (Similar ecology to *O.curzoniae*)

**Status:** UNMODELLABLE; **Included in final analysis:** X

| Model evaluation metric |  |
| --- | --- |
| AUC | 0.99 |
| Omission rate | 0.00 |
| Sensitivity | 1.00 |
| Specificity | 0.99 |
| Proportion correct | 0.99 |
| **Kappa** | **0.35** |
| True Skill Statistic | 0.99 |

**Summary:** The Nubra’s pika’s bioclimatic envelope is predicted to increase by 1% with no latitudinal polewards shift, but a mean increase in elevation of ~200m driven by an increase in minimum and maximum elevation. 95% of the permutation importance of the model was contributed to by minimum precipitation (76.1%) and mean annual temperature (20.9%).

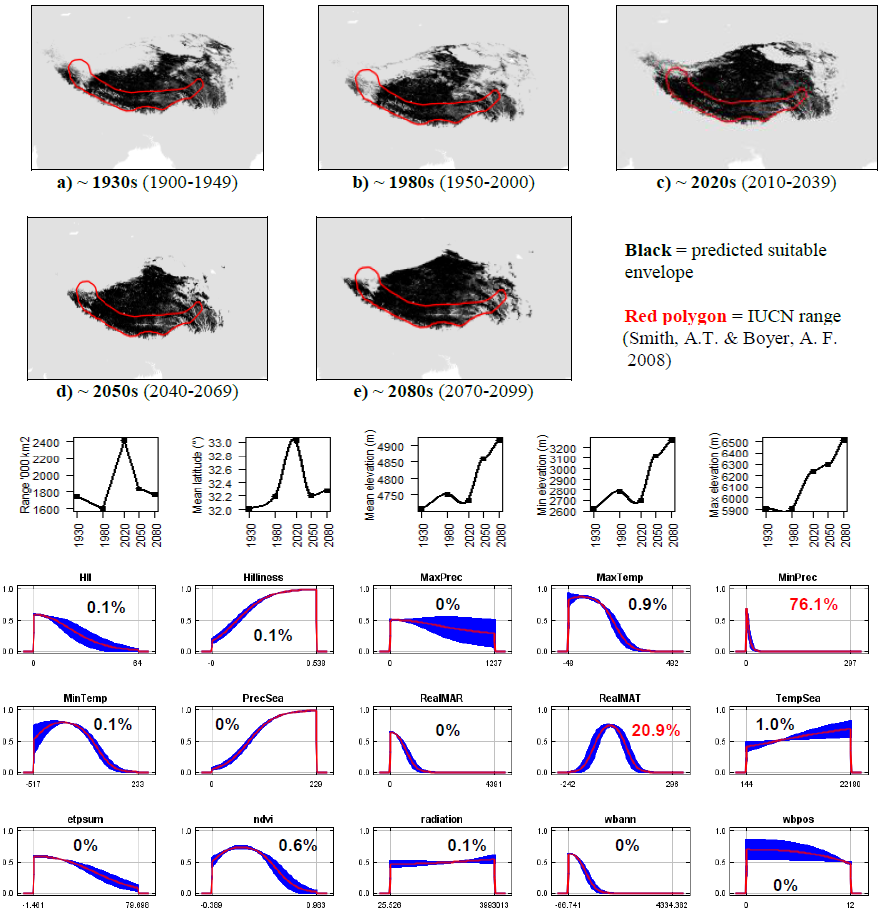

**#56 - Pallas’s pika** (*Ochotona pallasi*) *n* = 19

**Expert:** Andrew Smith, Arizona State University

**Expert evaluation:** Medium

**Data:** Only modern

**Envelope:** Climatic and habitat

**Dispersal distance:** 10km/year (Sokolov, V.E. *et al,* 2009)

**Status:** MODELLABLE; **Included in final analysis:** √

| Model evaluation metric |  |
| --- | --- |
| AUC | 0.99 |
| Omission rate | 0.00 |
| Sensitivity | 1.00 |
| Specificity | 0.99 |
| Proportion correct | 0.99 |
| **Kappa** | **0.46** |
| True Skill Statistic | 0.99 |

**Summary:** The Pallas’s pika’s bioclimatic envelope is predicted to decrease by 60% with a ~2° mean latitudinal polewards shift and a mean increase in elevation of ~40m driven by an increase in minimum elevation. 95% of the permutation importance of the model was contributed to by minimum precipitation (46.1%), mean annual precipitation (35.5%), mean annual temperature (9.4%) and minimum temperature (6.7%).

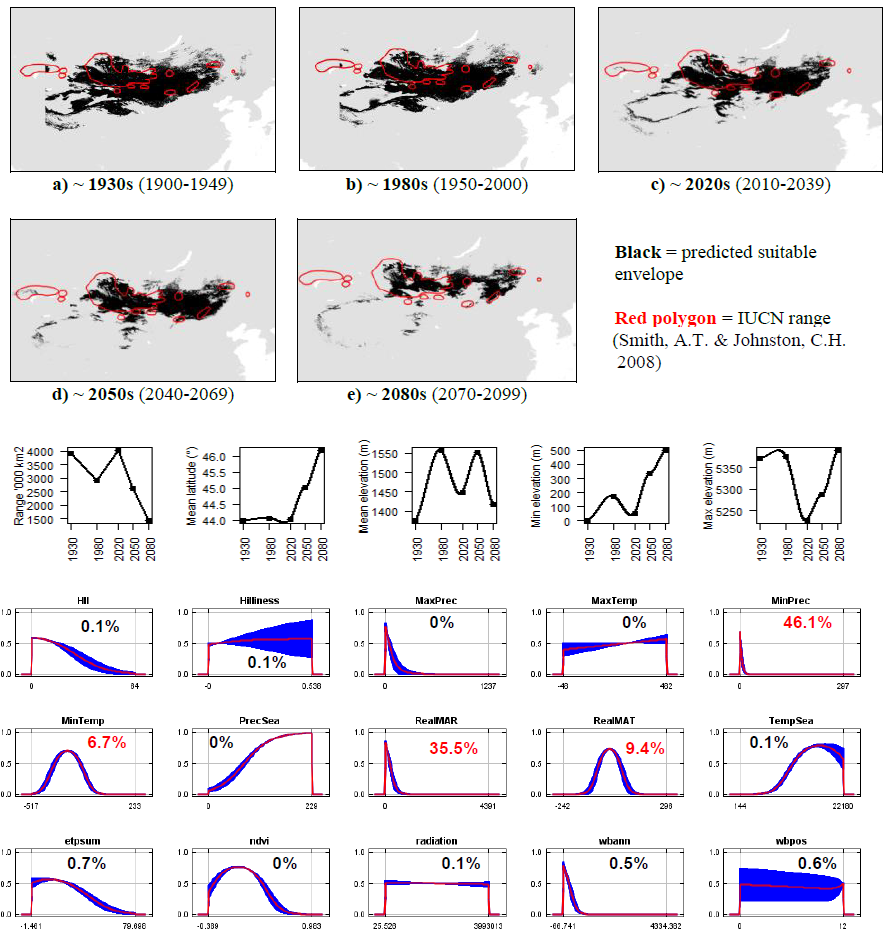

**#57 - American pika** (*Ochotona princeps*) *n* = 670

**Expert:** Andrew Smith, Arizona State University

**Expert evaluation:** Medium

**Data:** Only modern

**Envelope:** Climatic and habitat

**Dispersal distance:** 16.1km/year (Expert)

**Status:** MODELLABLE; **Included in final analysis:** √

| Model evaluation metric |  |
| --- | --- |
| AUC | 0.95 |
| Omission rate | 0.10 |
| Sensitivity | 0.9 |
| Specificity | 0.99 |
| Proportion correct | 0.98 |
| **Kappa** | **0.87** |
| True Skill Statistic | 0.89 |

**Summary:** The American pika’s bioclimatic envelope is predicted to decrease by 25% with a ~1° mean latitudinal polewards shift and a mean decrease in elevation of ~10m driven by an decrease in minimum elevation. 95% of the permutation importance of the model was contributed to by mean annual temperature (88.1%), annual evapotranspiration (5.0%) and maximum temperature (2.8%).

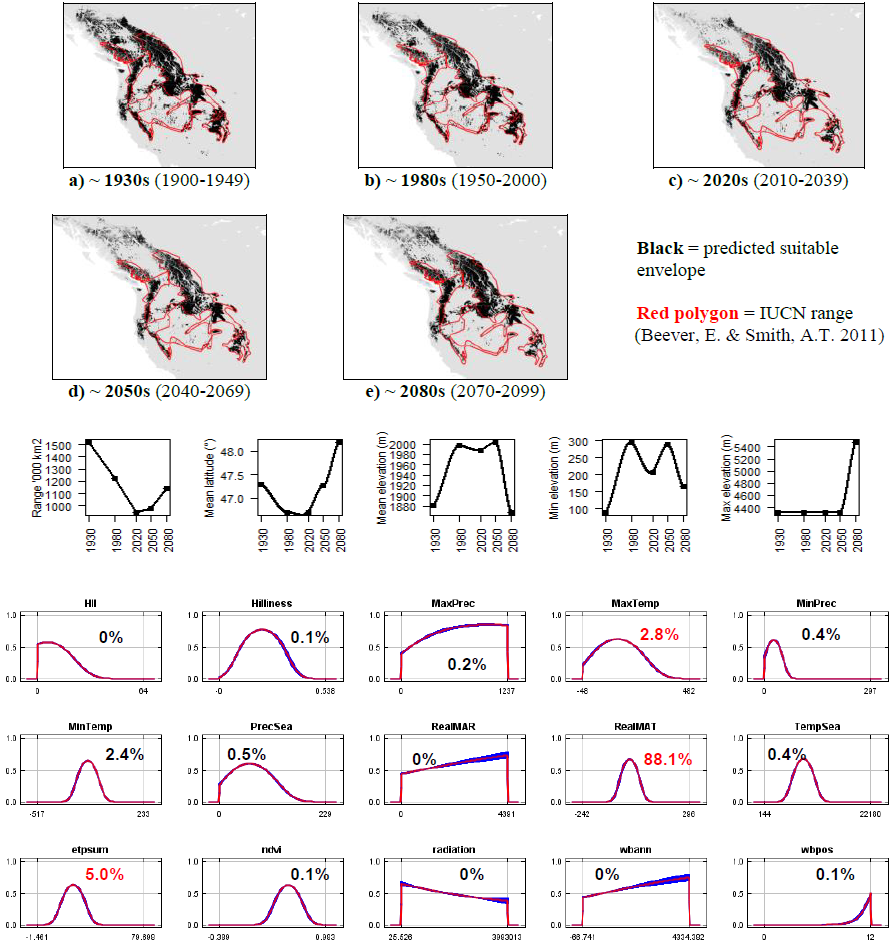

**#58 - Little pika** (*Ochotona pusilla*) *n* = 30

**Expert:** Andrew Smith, Arizona State University

**Expert evaluation:** Medium

**Data:** Modern and historic

**Envelope:** Climatic and habitat

**Dispersal distance:** 4km/year (Sokolov, V.E. *et al,* 2009)

**Status:** MODELLABLE; **Included in final analysis:** √

| Model evaluation metric |  |
| --- | --- |
| AUC | 0.93 |
| Omission rate | 0.13 |
| Sensitivity | 0.87 |
| Specificity | 0.99 |
| Proportion correct | 0.99 |
| **Kappa** | **0.58** |
| True Skill Statistic | 0.86 |

**Summary:** The Little pika’s bioclimatic envelope is predicted to increase by 20% with a ~1° mean latitudinal polewards shift and a mean increase in elevation of ~70m driven by an increase in maximum elevation. 95% of the permutation importance of the model was contributed to by temperature seasonality (70.2%), annual water balance (13.3%), mean annual temperature (10.2%) and minimum temperature (4.5%).

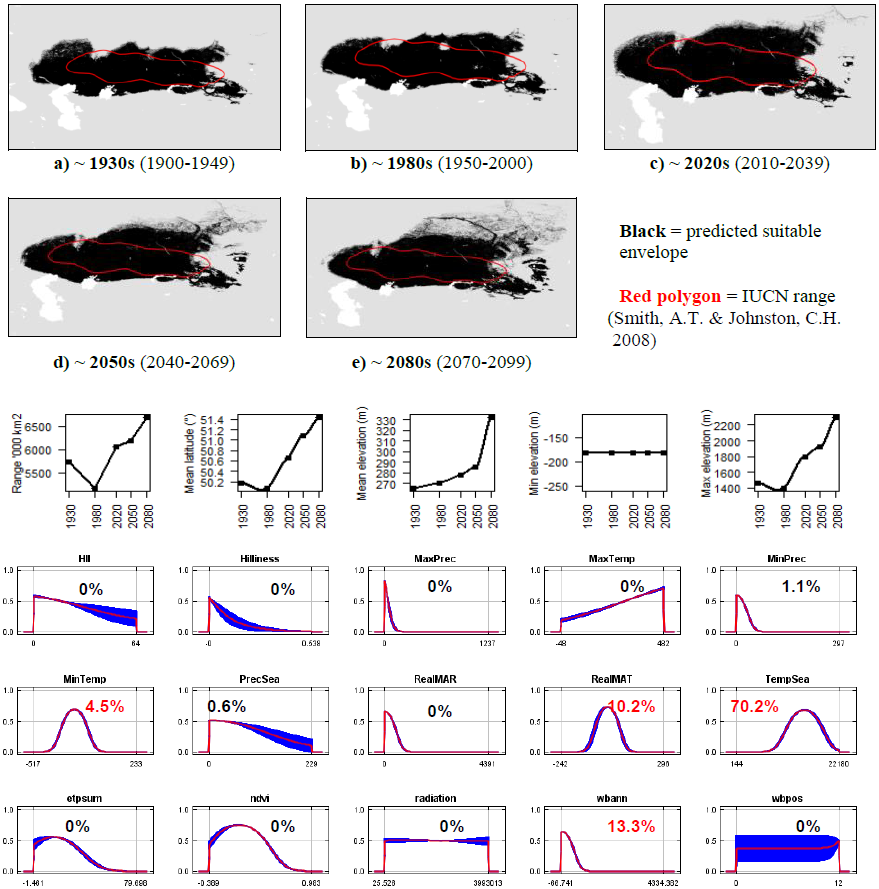

**#59 - Royle’s pika** (*Ochotona roylei*) *n* = 22

**Expert:** Sabuj Bhattacharya, Wildlife Institute of India

**Expert evaluation:** Medium

**Data:** Modern and historic

**Envelope:** Climatic and habitat

**Dispersal distance:** 1km/year (Expert)

**Status:** MODELLABLE; **Included in final analysis:** √

| Model evaluation metric |  |
| --- | --- |
| AUC | 0.98 |
| Omission rate | 0.05 |
| Sensitivity | 0.95 |
| Specificity | 0.99 |
| Proportion correct | 0.99 |
| **Kappa** | **0.76** |
| True Skill Statistic | 0.95 |

**Summary:** The Royle’s pika’s bioclimatic envelope is predicted to decrease by 20% with no latitudinal polewards shift and a mean increase in elevation of ~340m driven by an increase in minimum elevation. 95% of the permutation importance of the model was contributed to by minimum precipitation (53.2%), maximum temperature (20.4%), temperature seasonality (10.2%), human influence index (5.5%), surface roughness index (4.2%) and precipitation seasonality (3.2%).

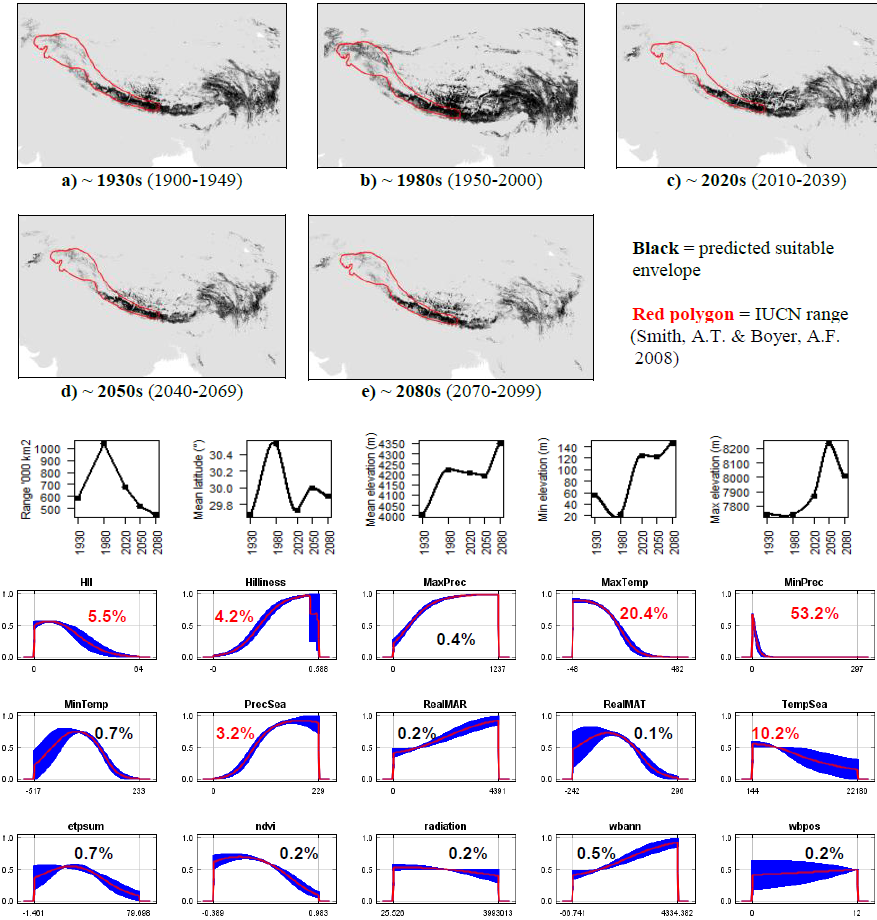

**#60 - Afghan pika** (*Ochotona rufescens*) *n* = 17

**Expert:** Chelmala Srinivasulu, Osmania University, India

**Expert evaluation:** Medium

**Data:** Modern and historic

**Envelope:** Climatic and habitat

**Dispersal distance:** 3km/year (Average for Asian pikas)

**Status:** MODELLABLE; **Included in final analysis:** √

| Model evaluation metric |  |
| --- | --- |
| AUC | 0.99 |
| Omission rate | 0.00 |
| Sensitivity | 1.00 |
| Specificity | 0.99 |
| Proportion correct | 0.99 |
| **Kappa** | **0.74** |
| True Skill Statistic | 0.99 |

**Summary:** The Afghan pika’s bioclimatic envelope is predicted to increase by 5% with a ~1° mean latitudinal polewards shift and a mean increase in elevation of ~380m driven by an increase in maximum elevation. 95% of the permutation importance of the model was contributed to by minimum precipitation (89.2%), minimum temperature (5.5%) and normalised difference vegetation index (2.3%).

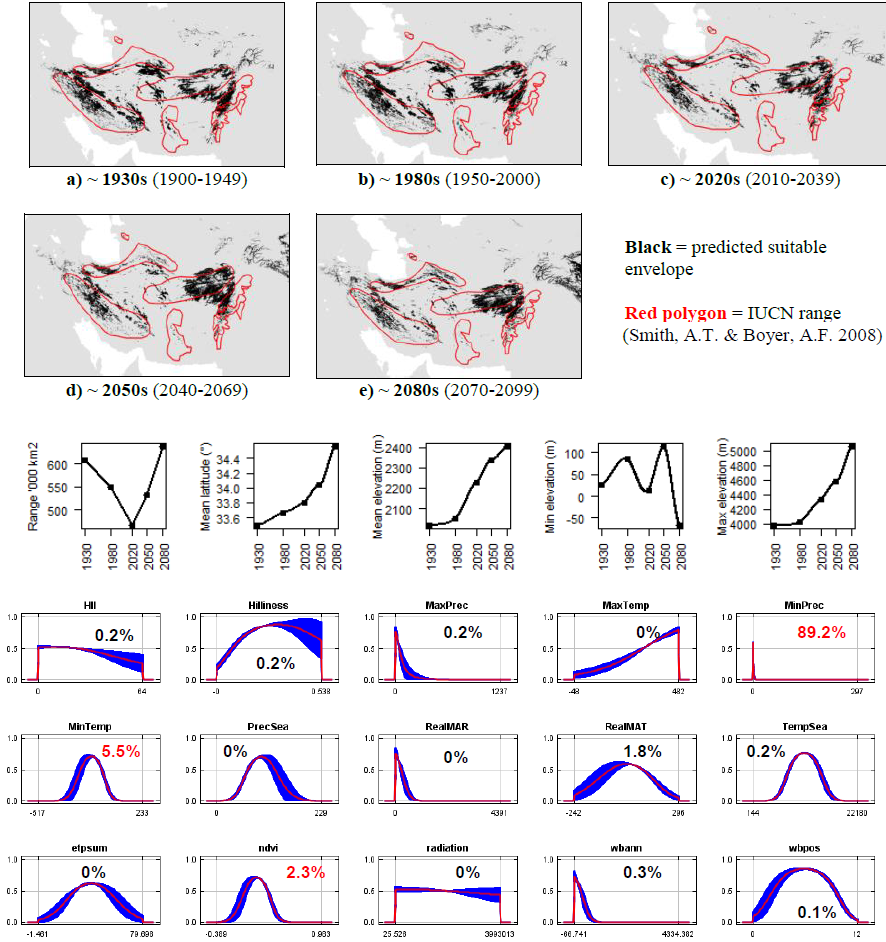

**#61 - Turkestan red pika** (*Ochotona rutila*) *n* = 13

**Expert:** Andrey Lissovsky, Zoological Museum of Moscow State University

**Expert evaluation:** Poor

**Data:** Modern and historic

**Envelope:** Climatic and habitat

**Dispersal distance:** 3km/year (Average for Asian pikas)

**Status:** UNMODELLABLE; **Included in final analysis:** X

| Model evaluation metric |  |
| --- | --- |
| AUC | 0.99 |
| Omission rate | 0.00 |
| Sensitivity | 1.00 |
| Specificity | 0.99 |
| Proportion correct | 0.99 |
| **Kappa** | **0.25** |
| True Skill Statistic | 0.99 |

**Summary:** The Turkestan red pika’s bioclimatic envelope is predicted to decrease by 10% with a ~1° mean latitudinal shift towards the Equator and a mean increase in elevation of ~630m driven by an increase in maximum elevation. 95% of the permutation importance of the model was contributed to by minimum precipitation (82.5%), minimum temperature (5.9%), human influence index (3.0%), precipitation seasonality (2.5%) and surface roughness index (1.7%).

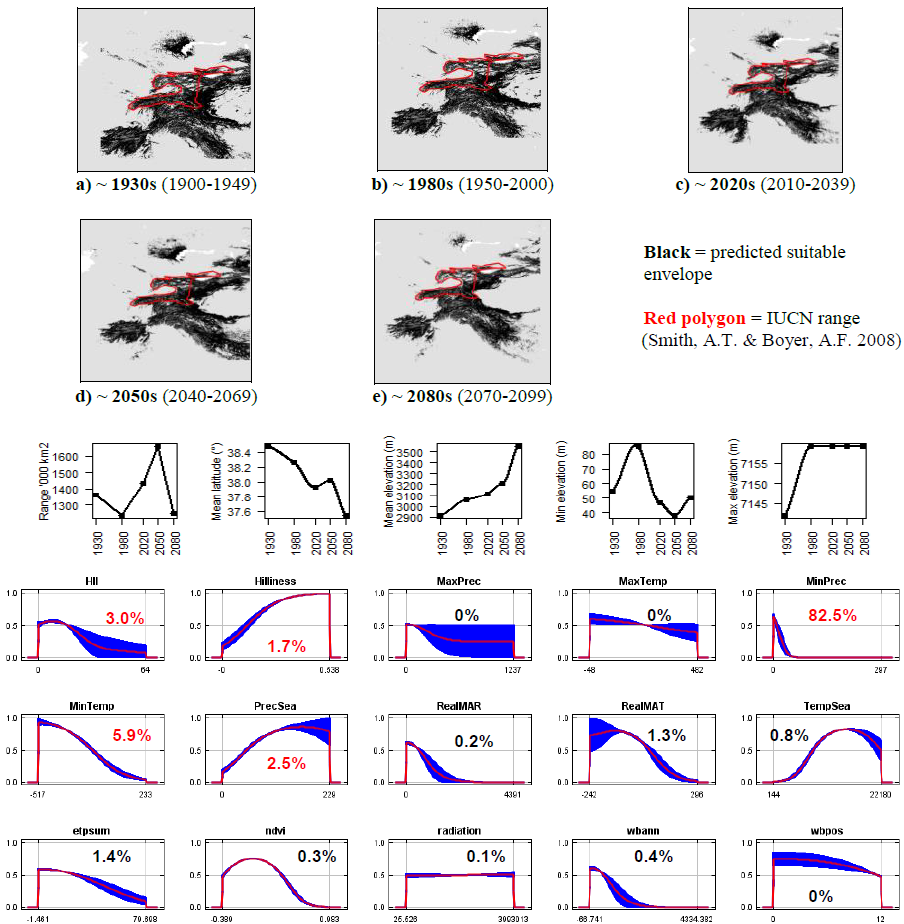

**#62 - Moupin pika** (*Ochotona thibetana) n =*95

**Expert:** Deyan Ge, Institute of Zoology, Chinese Academy of Sciences

**Expert evaluation:** Poor

**Data:** Modern and historic

**Envelope:** Climatic and habitat

**Dispersal distance:** 2km/year (Similar ecology to *O.roylei*)

**Status:** UNMODELLABLE; **Included in final analysis:** X

| Model evaluation metric |  |
| --- | --- |
| AUC | 0.93 |
| Omission rate | 0.13 |
| Sensitivity | 0.87 |
| Specificity | 0.99 |
| Proportion correct | 0.99 |
| **Kappa** | **0.52** |
| True Skill Statistic | 0.86 |

**Summary:** The Moupin pika’s bioclimatic envelope is predicted to increase by 10% with a ~2° mean latitudinal polewards shift and a mean increase in elevation of ~370m driven by an increase in maximum elevation. 95% of the permutation importance of the model was contributed to by minimum precipitation (43.5%), maximum temperature (38.5%), temperature seasonality (7.7%), annual evapotranspiration (3.7%) and mean annual precipitation (3.1%).

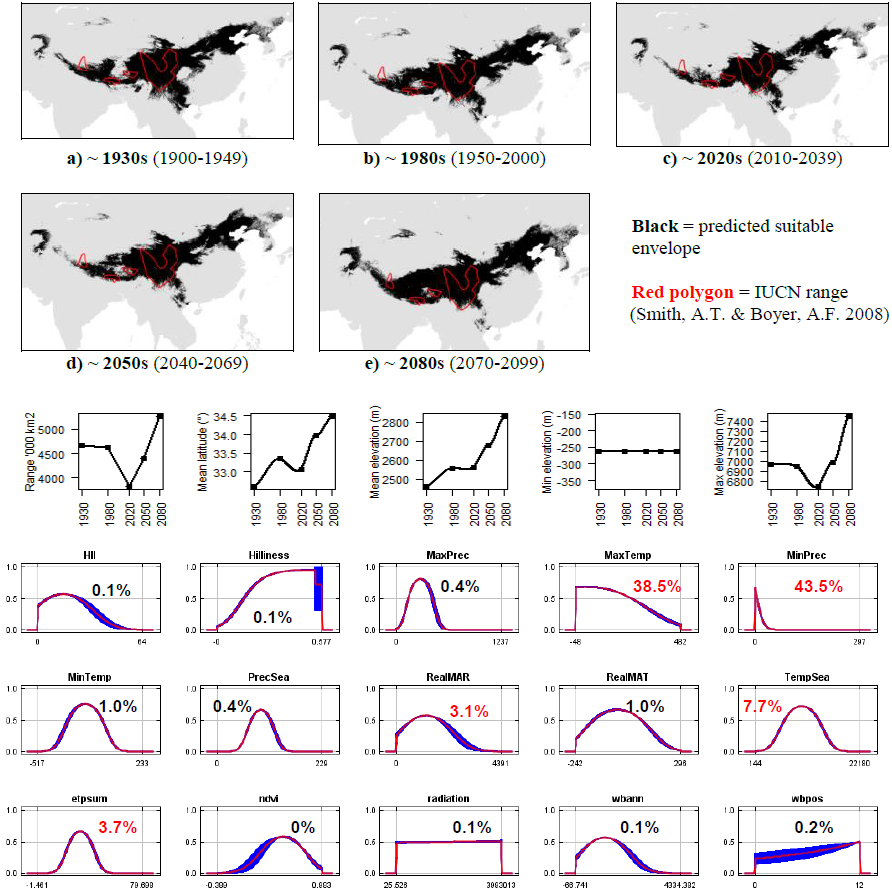

**#63 - Thomas’s pika** (*Ochotona thomasi*) *n* = 16

**Expert:** Andrew Smith, Arizona State University

**Expert evaluation:** Good

**Data:** Modern and historic

**Envelope:** Climatic and habitat

**Dispersal distance:** 1km/year (Similar ecology to *O.koslowi*)

**Status:** MODELLABLE; **Included in final analysis:** √

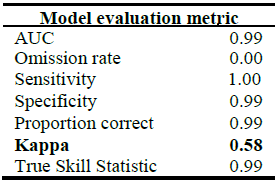

**Summary:** The Thomas’s pika’s bioclimatic envelope is predicted to decrease by 70% with a ~1.5° mean latitudinal polewards shift and a mean increase in elevation of ~590m driven by an increase in maximum and minimum elevation. 95% of the permutation importance of the model was contributed to by minimum precipitation (56.6%), mean annual temperature (23.4%), minimum temperature (14.2%) and evapotranspiration (2.8%).

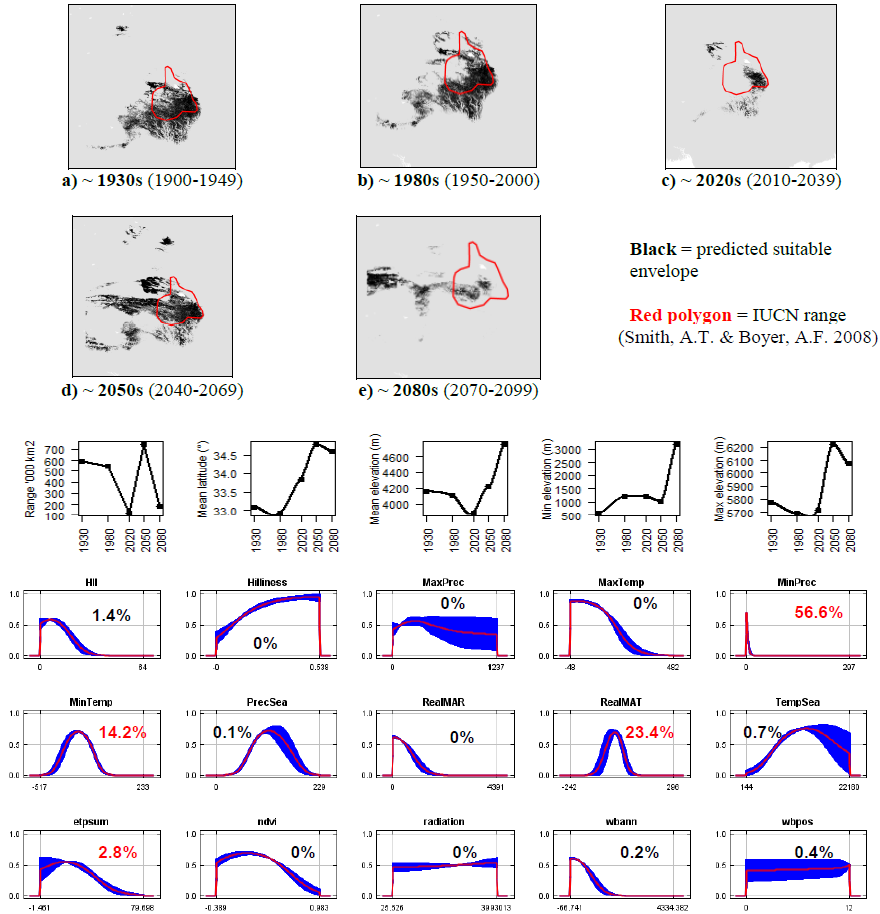

**#64 - Turuchan pika** (*Ochotona turuchanensis*) *n* = 30

**Expert:** Andrey Lissovsky, Zoological Museum of Moscow State University

**Expert evaluation:** Medium

**Data:** Modern and historic

**Envelope:** Climatic and habitat

**Dispersal distance:** 15km/year (Expert)

**Status:** MODELLABLE; **Included in final analysis:** √

| Model evaluation metric |  |
| --- | --- |
| AUC | 0.93 |
| Omission rate | 0.13 |
| Sensitivity | 0.87 |
| Specificity | 0.99 |
| Proportion correct | 0.99 |
| **Kappa** | **0.50** |
| True Skill Statistic | 0.86 |

**Summary:** The Turuchan pika’s bioclimatic envelope is predicted to increase by 50% with a ~2° mean latitudinal polewards shift and a mean increase in elevation of ~70m driven by an increase in minimum elevation. 95% of the permutation importance of the model was contributed to by minimum temperature (21.6%), minimum precipitation (19.9%), human influence index (15.2%), annual evapotranspiration (10.9%).

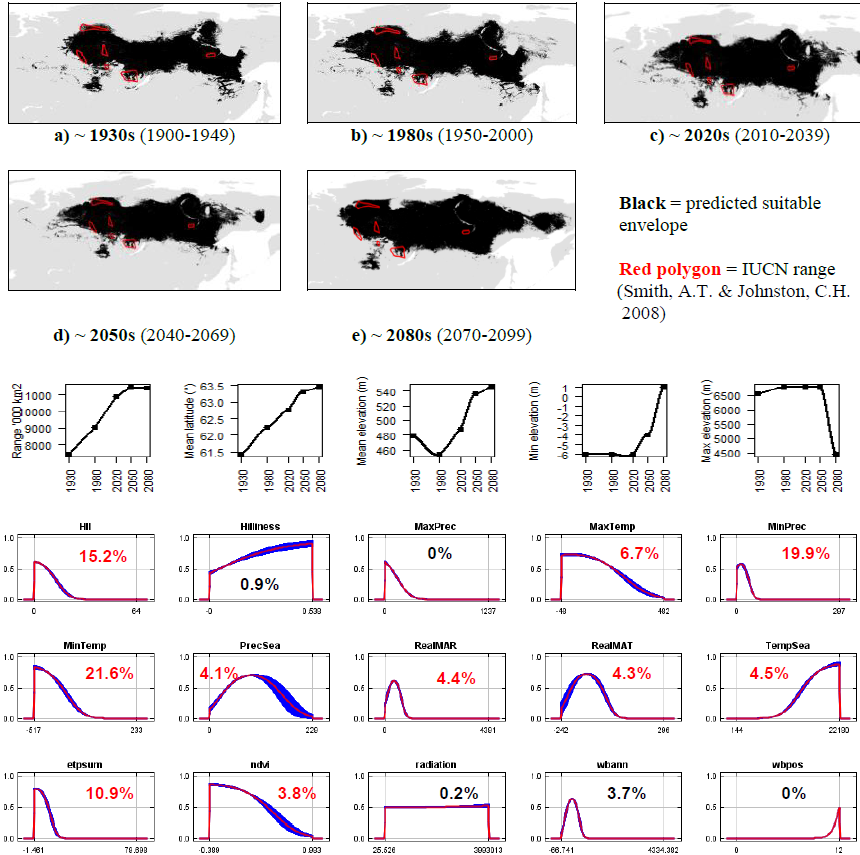

**#65 - European rabbit** (*Oryctolagus cunciculus*) *n* = 22,712

**Expert:** Neil Reid, Queen’s University Belfast

**Expert evaluation:** Medium

**Data:** Modern and historic

**Envelope:** Climatic and habitat

**Dispersal distance:** 1km/year (Expert)

**Status:** UNMODELLABLE; **Included in final analysis:** X

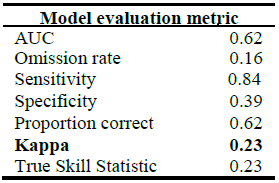

| Model evaluation metric |  |
| --- | --- |
| AUC | 0.62 |
| Omission rate | 0.16 |
| Sensitivity | 0.84 |
| Specificity | 0.39 |
| Proportion correct | 0.62 |
| **Kappa** | **0.23** |
| True Skill Statistic | 0.23 |

**Summary:** The European rabbit’s bioclimatic envelope is predicted to increase by 30% with a ~2° mean latitudinal polewards shift and a mean decrease in elevation of ~10m. 95% of the permutation importance of the model was contributed to by annual evapotranspiration (21.1%), mean annual temperature (19.0%), minimum temperature (13.9%), annual water balance (12.3%), mean annual precipitation (11.2%), human influence index (9.8%), precipitation seasonality (7.6%) and normalised difference vegetation index (2.9%).

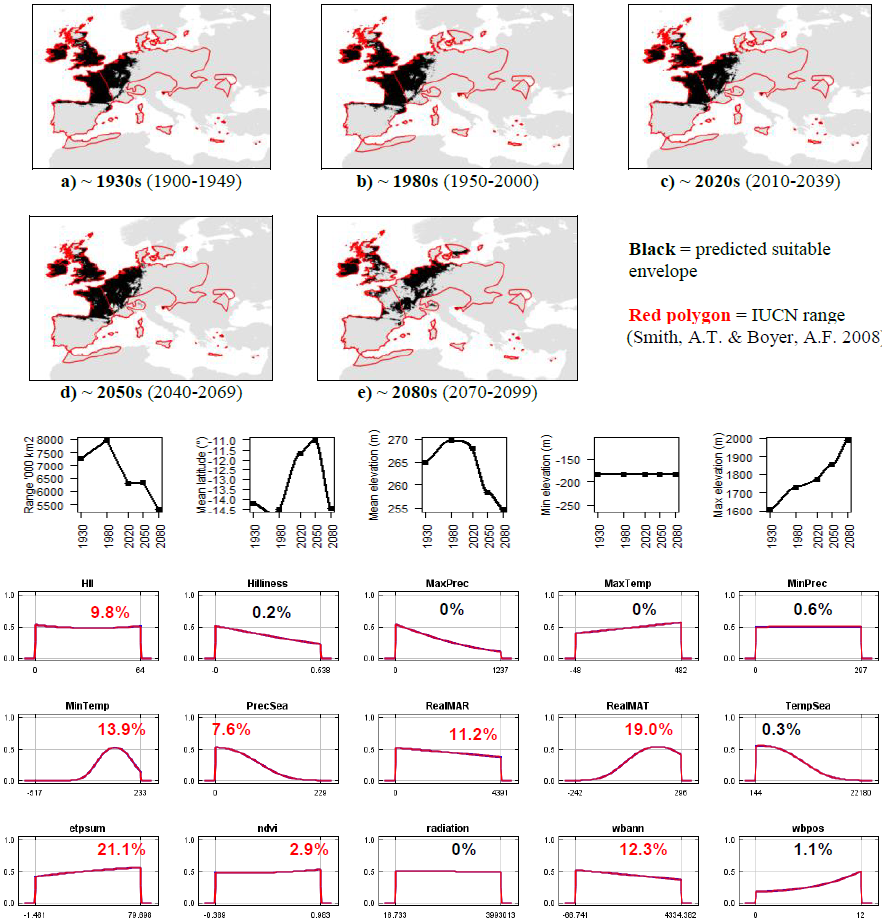

**#66 - Amami rabbit** (*Pentalagus furnessi*) *n* = 9

**Expert:** Fumio Yamada, Forestry and Forest Products Research Institute, Japan

**Expert evaluation:** Good

**Data:** Modern and historic

**Envelope:** Climatic and habitat

**Dispersal distance:** 0.01km/year (Expert)

**Status:** MODELLABLE; **Included in final analysis:** √

| Model evaluation metric |  |
| --- | --- |
| AUC | 0.99 |
| Omission rate | 0.00 |
| Sensitivity | 0.99 |
| Specificity | 0.99 |
| Proportion correct | 0.99 |
| **Kappa** | **0.95** |
| True Skill Statistic | 0.99 |

**Summary:** The Amami rabbit’s bioclimatic envelope is predicted to increase by 150% with a ~1° mean latitudinal shift towards the Equator and a mean decrease in elevation of ~25m. 95% of the permutation importance of the model was contributed to by minimum temperature (97.3%), temperature seasonality (1.7%) and annual evapotranspiration (0.8%).

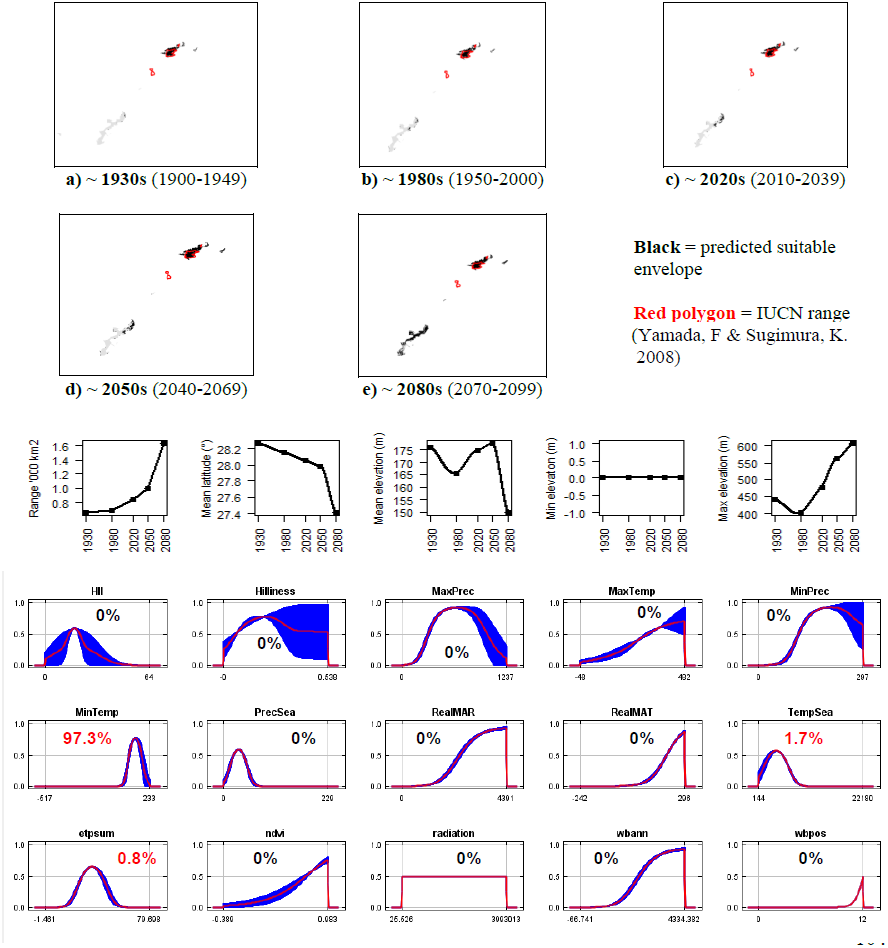

**#67 - Bunyoro rabbit** (*Poelagus marjorita*) *n* = 8

**Expert:** David Happold, Australian National University

**Expert evaluation:** Poor

**Data:** Only modern

**Envelope:** Climatic and habitat

**Dispersal distance:** 2km/year (Similar ecology to Pronolagus sp.)

**Status:** UNMODELLABLE; **Included in final analysis:** X

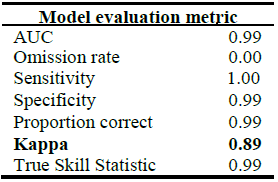

**Summary:** The Bunyoro rabbit’s bioclimatic envelope is predicted to decrease by 90% with a ~1° mean latitudinal shift towards the Equator and a mean increase in elevation of ~200m driven by an increase in minimum elevation.. 95% of the permutation importance of the model was contributed to by annual evapotranspiration (61.7%), temperature seasonality (31.4%) and number of months with a positive water balance (6.3%).

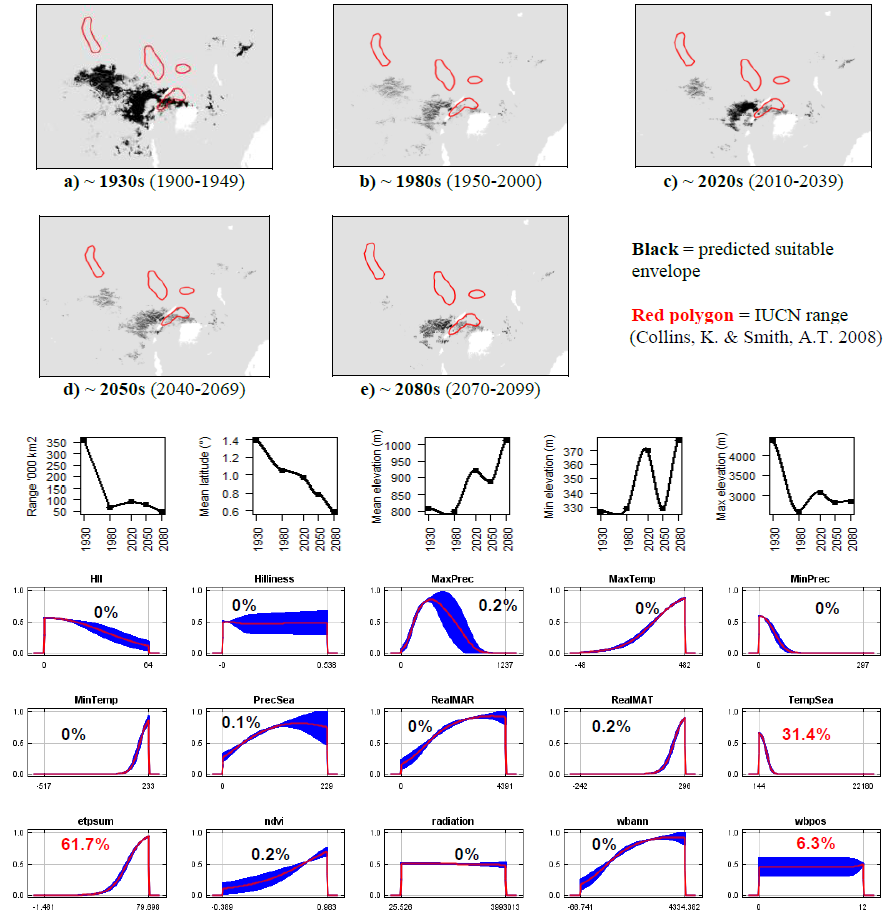

**#68 - Greater red rock hare** (*Pronolagus crassicaudatus*)

*n* = 7

**Expert:** Kai Collins, University of Pretoria

**Expert evaluation:** Poor

**Data:** Only modern

**Envelope:** Climatic and habitat

**Dispersal distance:** 2km/year (Similar ecology to *P.randensis*)

**Status:** UNMODELLABLE; **Included in final analysis:** X

| Model evaluation metric |  |
| --- | --- |
| AUC | 0.99 |
| Omission rate | 0.00 |
| Sensitivity | 1.00 |
| Specificity | 0.98 |
| Proportion correct | 0.98 |
| **Kappa** | **0.06** |
| True Skill Statistic | 0.98 |

**Summary:** The Greater red rock hare’s bioclimatic envelope is predicted to decrease by 65% with a ~3° mean latitudinal polewards shift and a mean decrease in elevation of ~20m driven by a decrease in maximum elevation. 95% of the permutation importance of the model was contributed to by human influence index (31.7%), temperature seasonality (29.3%) solar radiation (16.9%) and minimum precipitation (18.3%).

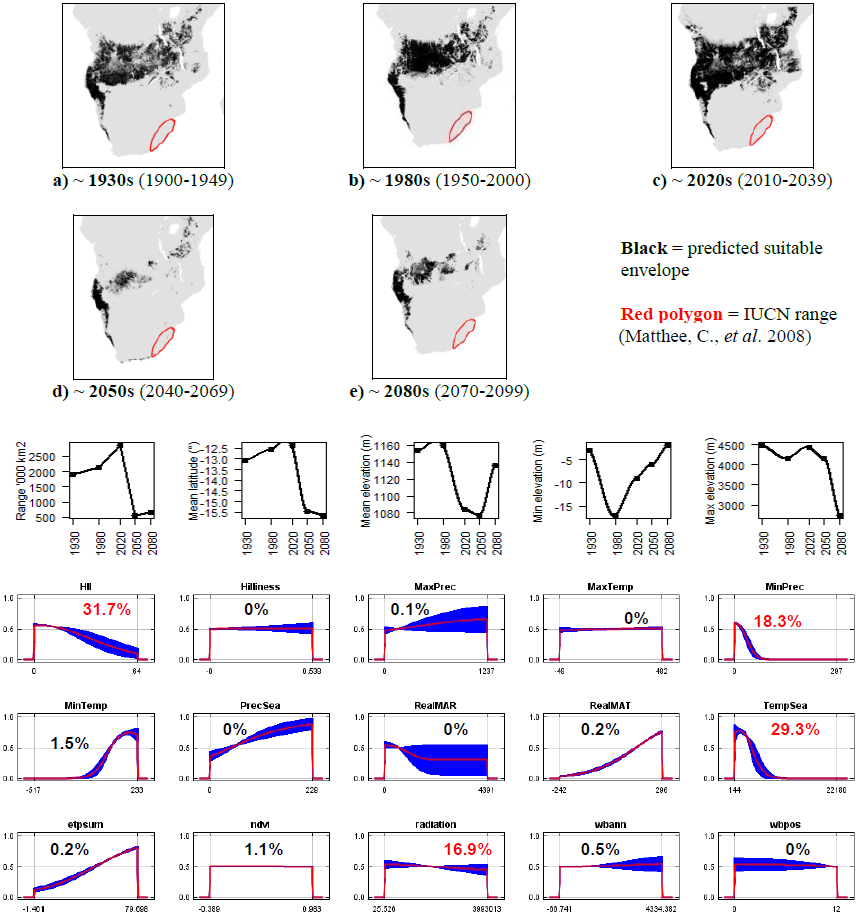

**#69 - Jameson’s red rock hare** (*Pronolagus randensis*) *n* = 27

**Expert:** Kai Collins, University of Pretoria

**Expert evaluation:** Poor

**Data:** Modern and historic

**Envelope:** Climatic and habitat

**Dispersal distance:** 2km/year (Expert)

**Status:** UNMODELLABLE; **Included in final analysis:** X

| Model evaluation metric |  |
| --- | --- |
| AUC | 0.98 |
| Omission rate | 0.04 |
| Sensitivity | 0.96 |
| Specificity | 0.99 |
| Proportion correct | 0.99 |
| **Kappa** | **0.55** |
| True Skill Statistic | 0.96 |

**Summary:** The Jameson’s red rock hare’s bioclimatic envelope is predicted to decrease by 70% with a ~5° mean latitudinal polewards shift and a mean increase in elevation of ~325m driven by an increase in maximum and minimum elevation. 95% of the permutation importance of the model was contributed to by mean annual temperature (46.2%), maximum temperature (23.3%), minimum precipitation (16.3%), minimum temperature (7.0%) and temperature seasonality (1.4%).

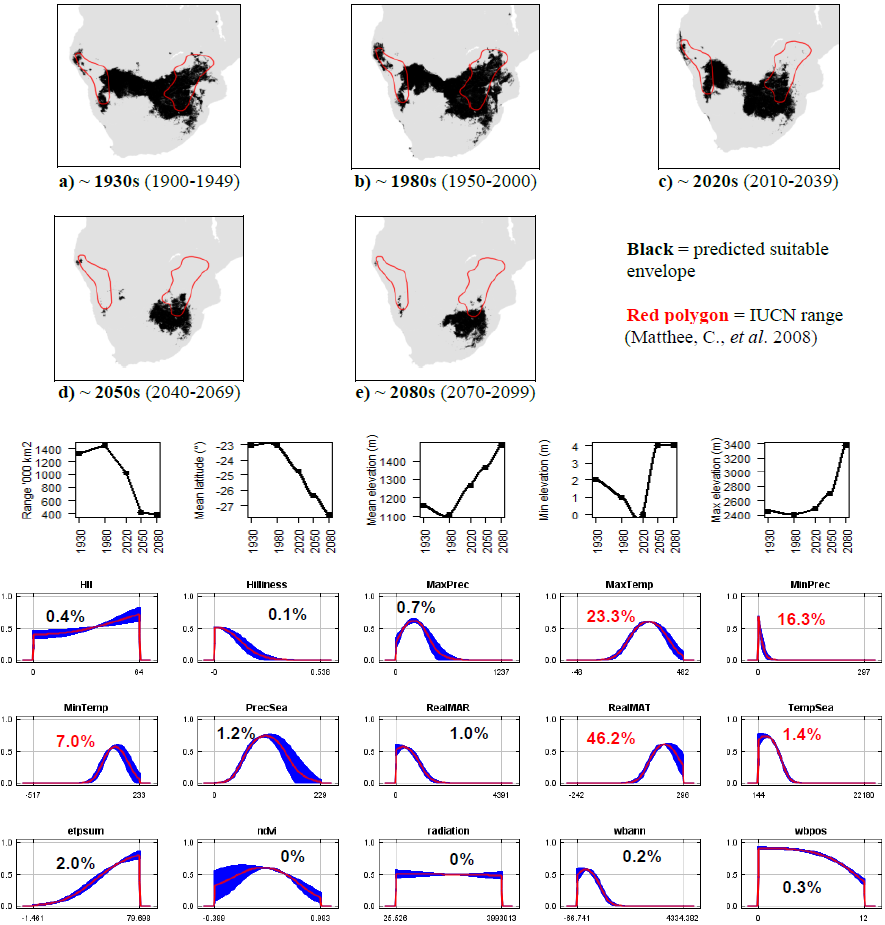

**#70 - Smith’s red rock hare** (*Pronolagus rupestris*) *n* = 9

**Expert:** Kai Collins, University of Pretoria

**Expert evaluation:** Poor

**Data:** Modern and historic

**Envelope:** Climatic and habitat

**Dispersal distance:** 2km/year (Similar ecology to *P.randensis*)

**Status:** UNMODELLABLE; **Included in final analysis:** X

| Model evaluation metric |  |
| --- | --- |
| AUC | 0.99 |
| Omission rate | 0.00 |
| Sensitivity | 1.00 |
| Specificity | 0.98 |
| Proportion correct | 0.98 |
| **Kappa** | **0.07** |
| True Skill Statistic | 0.98 |

**Summary:** The Smith’s red rock hare’s bioclimatic envelope is predicted to decrease by 60% with a ~5° mean latitudinal polewards shift and a mean decrease in elevation of ~60m. 95% of the permutation importance of the model was contributed to by temperature seasonality (44.0%), minimum precipitation (18.5%), mean annual precipitation (18.3%), normalised difference vegetation index (6.3%), human influence index (5.2%) and minimum temperature (3.4%).

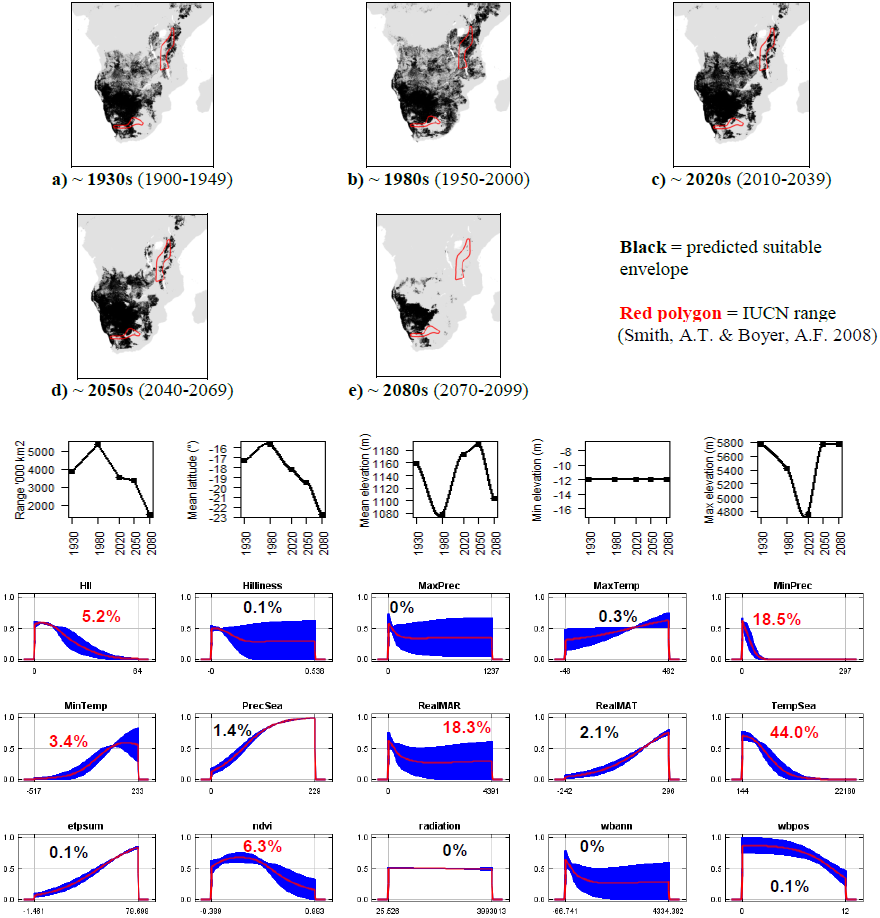

**#71 - Hewitt’s red rock hare** (*Pronolagus saundersiae*) *n* = 9

**Expert:** Kai Collins, University of Pretoria

**Expert evaluation:** Poor

**Data:** Modern and historic

**Envelope:** Climatic and habitat

**Dispersal distance:** 2km/year (Similar ecology to *P.randensis*)

**Status:** UNMODELLABLE; **Included in final analysis:** X

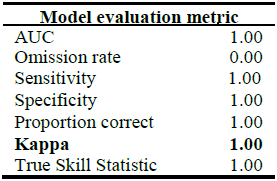

**Summary:** The Hewitt’s red rock hare’s bioclimatic envelope is predicted to decrease by 100% with a ~1° mean latitudinal polewards shift and a mean increase in elevation of ~15m driven by an increase in minimum elevation. 95% of the permutation importance of the model was contributed to by temperature seasonality (28.7%), precipitation seasonality (24.3%), maximum temperature (18.0%), annual evapotranspiration (16.0%), human influence index (5.6%), minimum temperature (1.9%) and minimum precipitation (1.6%).

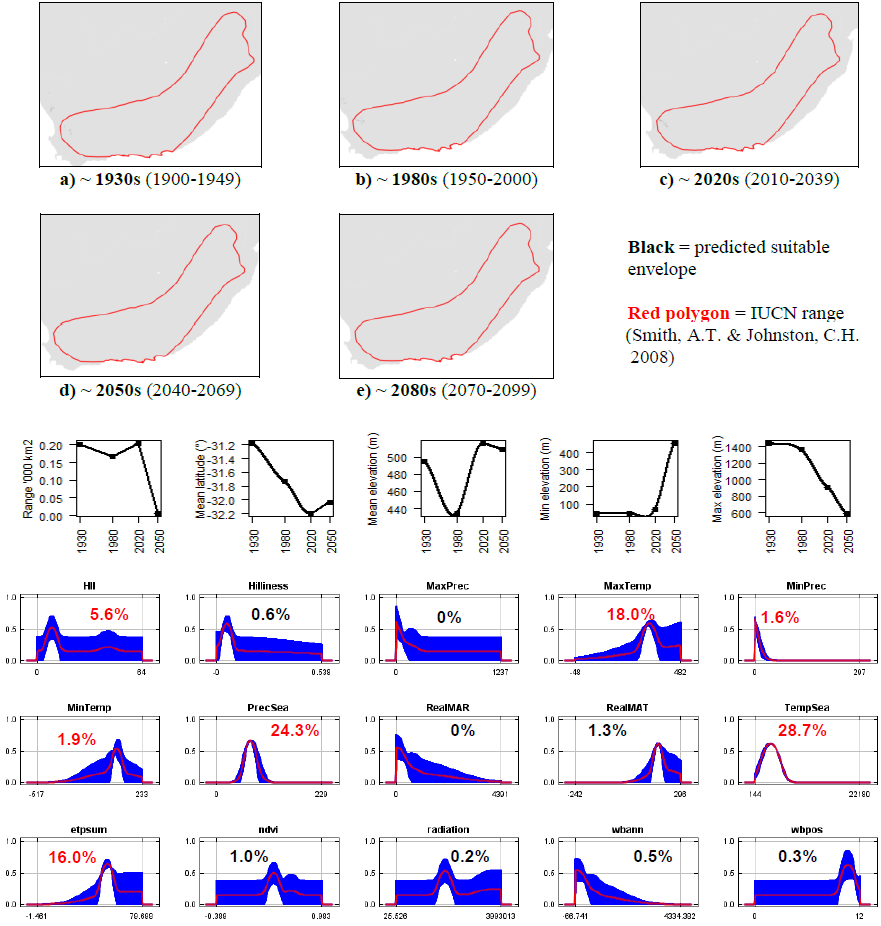

**#72 - Volcano rabbit** (*Romerolagus diazi) n =*31

**Expert:** Jose Antonio Martinez-Garcia, Universidad Autonoma Metropolitana, Mexico

**Expert evaluation:** Poor

**Data:** Only modern

**Envelope:** Climatic and habitat

**Dispersal distance:** 0.01km/year (Average for island species)

**Status:** UNMODELLABLE; **Included in final analysis:** X

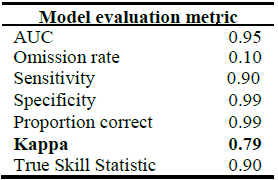

| Model evaluation metric |  |
| --- | --- |
| AUC | 0.95 |
| Omission rate | 0.10 |
| Sensitivity | 0.90 |
| Specificity | 0.99 |
| Proportion correct | 0.99 |
| **Kappa** | **0.79** |
| True Skill Statistic | 0.90 |

**Summary:** The Volcano rabbit’s bioclimatic envelope is predicted to decrease by 100% with a ~0.2° mean latitudinal polewards shift and a mean increase in elevation of ~1500m driven by increases in minimum and maximum elevation. 95% of the permutation importance of the model was contributed to by temperature seasonality (75.2%), precipitation seasonality (16.2%) and minimum temperature (6.8%).

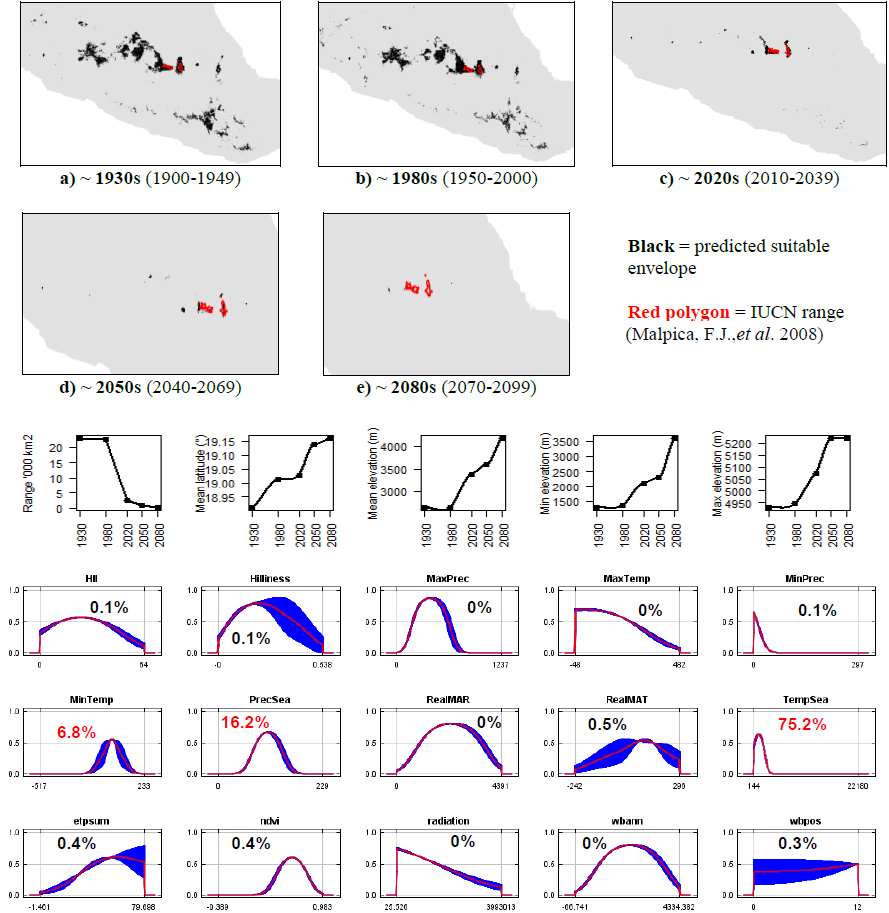

**#73 - Swamp rabbit** (*Sylvilagus aquaticus*) *n* = 66

**Expert:** Robert Kissell, Memphis State University

**Expert evaluation:** Medium

**Data:** Modern and historic

**Envelope:** Climatic and habitat

**Dispersal distance:** 25km/year (Expert)

**Status:** MODELLABLE; **Included in final analysis:** √

| Model evaluation metric |  |
| --- | --- |
| AUC | 0.95 |
| Omission rate | 0.09 |
| Sensitivity | 0.91 |
| Specificity | 0.99 |
| Proportion correct | 0.99 |
| **Kappa** | **0.76** |
| True Skill Statistic | 0.91 |

**Summary:** The Swamp rabbit’s bioclimatic envelope is predicted to increase by 200% with a ~2° mean latitudinal polewards shift and a mean increase in elevation of ~60m driven by an increase in maximum elevation. 95% of the permutation importance of the model was contributed to by temperature seasonality (38.1%), mean annual temperature (36.7%), mean annual precipitation (12.7%), precipitation seasonality (4.7%) and minimum precipitation (3.6%).

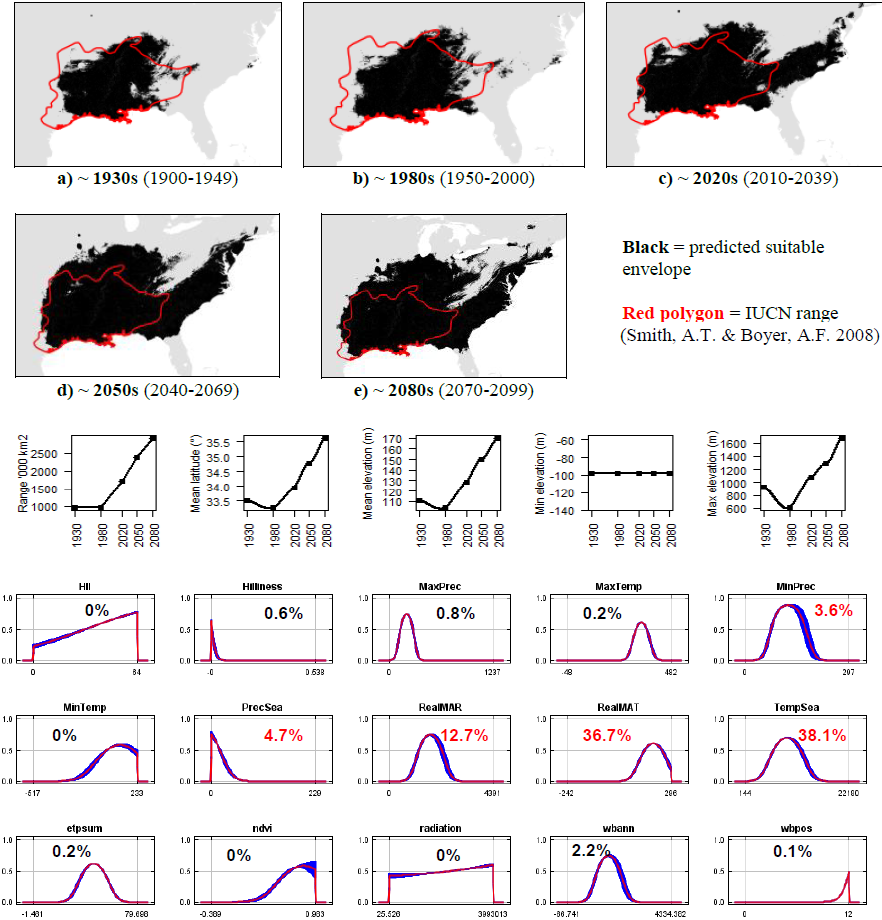

**#74 - Desert cottontail** (*Sylvilagus audubonii*) *n* = 1040

**Expert:** Consuelo Lorenzo, Departamento Conservation de la Biodiversidad, Chiapas

**Expert evaluation:** Medium

**Data:** Modern and historic

**Envelope:** Climatic and habitat

**Dispersal distance:** 7.5km/year (Similar ecology to *S.palustris*)

**Status:** MODELLABLE; **Included in final analysis:** √

| Model evaluation metric |  |
| --- | --- |
| AUC | 0.94 |
| Omission rate | 0.08 |
| Sensitivity | 0.92 |
| Specificity | 0.96 |
| Proportion correct | 0.96 |
| **Kappa** | **0.78** |
| True Skill Statistic | 0.88 |

**Summary:** The Desert cottontail’s bioclimatic envelope is predicted to increase by 5% with a ~1° mean latitudinal polewards shift and a mean increase in elevation of ~30m driven by an increase in minimum elevation. 95% of the permutation importance of the model was contributed to by precipitation seasonality (36.6%), annual evapotranspiration (27.1%), mean annual temperature (13.6%), minimum temperature (9.1%), maximum temperature (3.0%), annual water balance (2.1%), minimum precipitation (1.9%) and human influence index (1.7%).

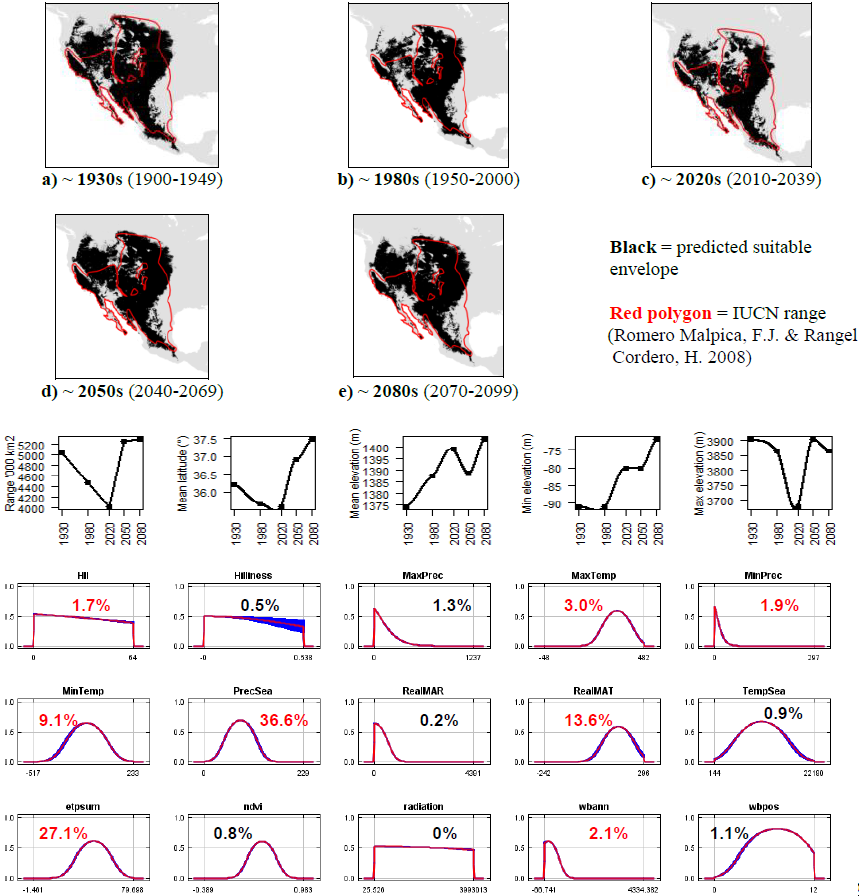

**#75 - Brush rabbit** (*Sylvilagus bachmani*) *n* = 263

**Expert:** Consuelo Lorenzo, Departamento Conservación de la

Biodiversidad, Chiapas

**Expert evaluation:** Medium

**Data:** Modern and historic

**Envelope:** Climatic and habitat

**Dispersal distance:** 3km/year (Similar ecology to *S.transitionalis*)

**Status:** MODELLABLE; **Included in final analysis:** √

| Model evaluation metric |  |
| --- | --- |
| AUC | 0.96 |
| Omission rate | 0.08 |
| Sensitivity | 0.92 |
| Specificity | 0.99 |
| Proportion correct | 0.99 |
| **Kappa** | **0.89** |
| True Skill Statistic | 0.91 |

**Summary:** The Brush rabbit’s bioclimatic envelope is predicted to decrease by 15% with a ~3° mean latitudinal polewards shift and a mean decrease in elevation of ~10m. 95% of the permutation importance of the model was contributed to by mean annual temperature (31.5%), minimum precipitation (24.5%), precipitation seasonality (14.4%), mean annual precipitation (7.1%), annual water balance (6.4%), minimum temperature (5.1%), normalised difference vegetation index (4.1%) and temperature seasonality (2.7%).

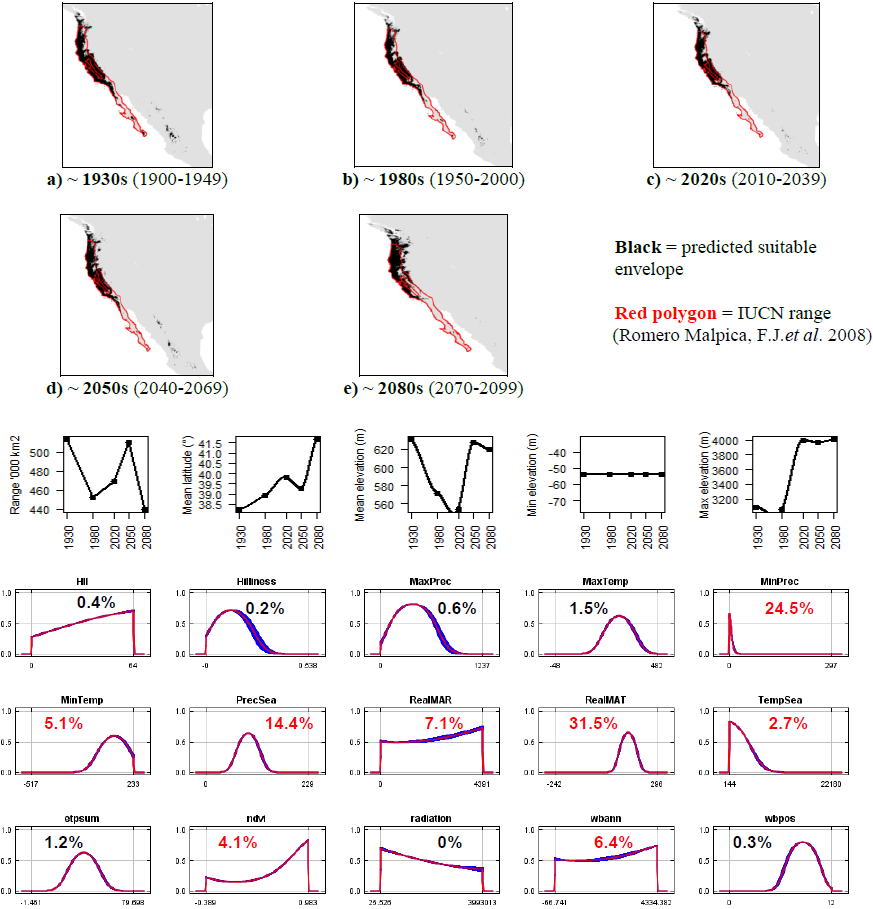

**#76 - Forest rabbit** (*Sylvilagus brasiliensis*) *n* = 181

**Expert:** Jorge Salazar-Bravo, Texas Tech University

**Expert evaluation:** Medium

**Data:** Modern and historic

**Envelope:** Climatic and habitat

**Dispersal distance:** 7.5km/year (Similar ecology to *S.palustris*)

**Status:** MODELLABLE; **Included in final analysis:** √

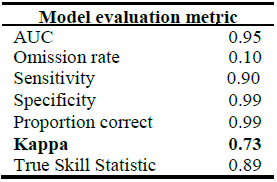

**Summary:** The Forest rabbit’s bioclimatic envelope is predicted to decrease by 50% with a ~6° mean latitudinal polewards shift and a mean increase in elevation of ~210m driven by an increase in minimum elevation. 95% of the permutation importance of the model was contributed to by annual evapotranspiration (71.4%), temperature seasonality (11.6%), normalised difference vegetation index (4.9%) and minimum temperature (4.6%).

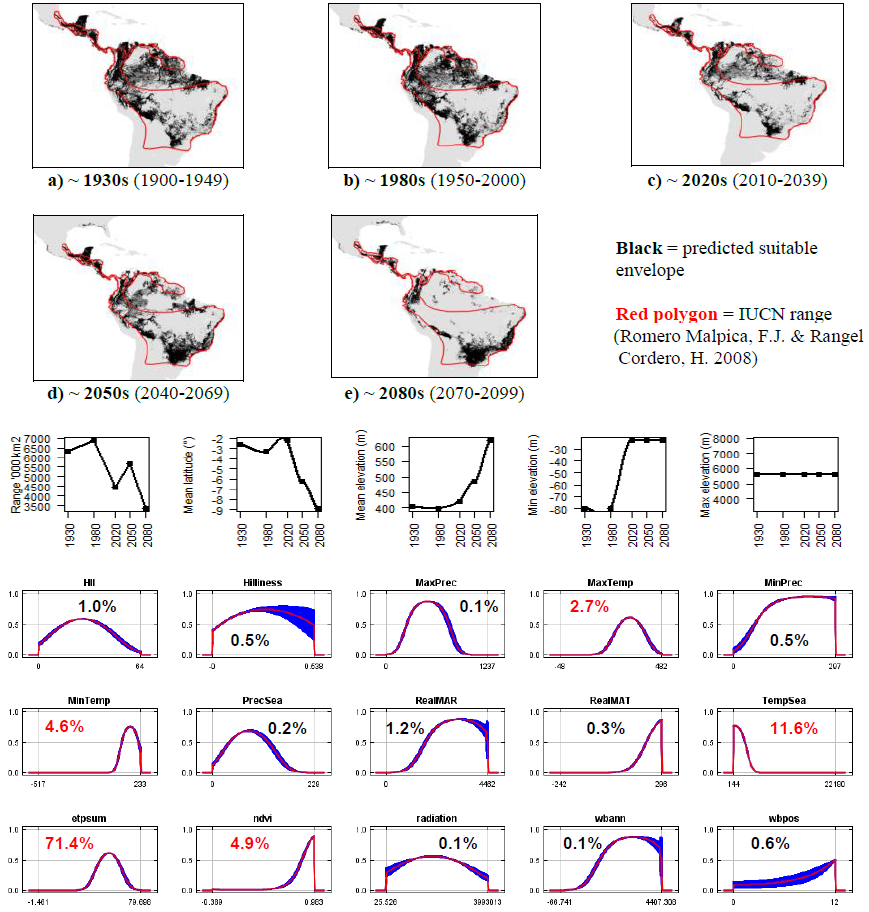

**#77 - Manzano mountain cottontail** (*Sylvilagus cognatus) n =* 7

**Expert:** Jennifer Frey, New Mexico State University

**Expert evaluation:** Medium

**Data:** Modern and historic

**Envelope:** Climatic and habitat

**Dispersal distance:** 0.01km/year (Similar ecology to *R.diazi*)

**Status:** MODELLABLE; **Included in final analysis:** √

| Model evaluation metric |  |
| --- | --- |
| AUC | 0.99 |
| Omission rate | 0.00 |
| Sensitivity | 1.00 |
| Specificity | 0.99 |
| Proportion correct | 0.99 |
| **Kappa** | **0.52** |
| True Skill Statistic | 0.99 |

**Summary:** The Manzano mountain cottontail’s bioclimatic envelope is predicted to decrease by 90% with a ~2° mean latitudinal polewards shift and a mean increase in elevation of ~230m driven by an increase in minimum elevation. 95% of the permutation importance of the model was contributed to by annual water balance (40.8%), minimum temperature (21.7%), precipitation seasonality (11.3%), mean annual temperature (8.2%), temperature seasonality (6.1%), minimum precipitation (4.3%) and mean annual precipitation (2.6%).

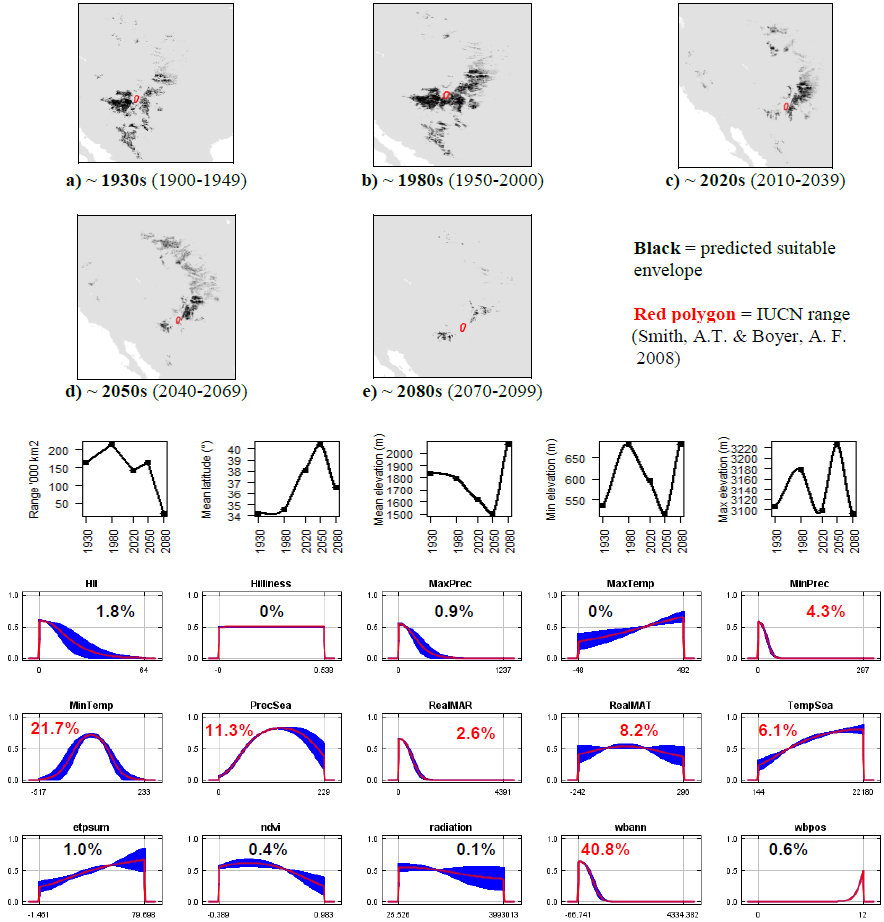

**#78 - Mexican cottontail** (*Sylvilagus cunicularius*) *n* = 76

**Expert:** Jorge Vazquez, Laboratorio de Ecologia del

Comportamiento, UAT-UNAM

**Expert evaluation:** Medium

**Data:** Only modern

**Envelope:** Climatic and habitat

**Dispersal distance:** 7.5km/year (Similar ecology to *S.palustris*)

**Status:** MODELLABLE; **Included in final analysis:** √

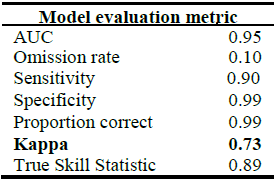

**Summary:** The Mexican cottontail’s bioclimatic envelope is predicted to decrease by 15% with a ~0.5° mean latitudinal polewards shift and a mean increase in elevation of ~200m driven by an increase in minimum elevation. 95% of the permutation importance of the model was contributed to by temperature seasonality (48.3%), precipitation seasonality (42.8%), normalised difference vegetation index (3.5%) and minimum temperature (2.2%).

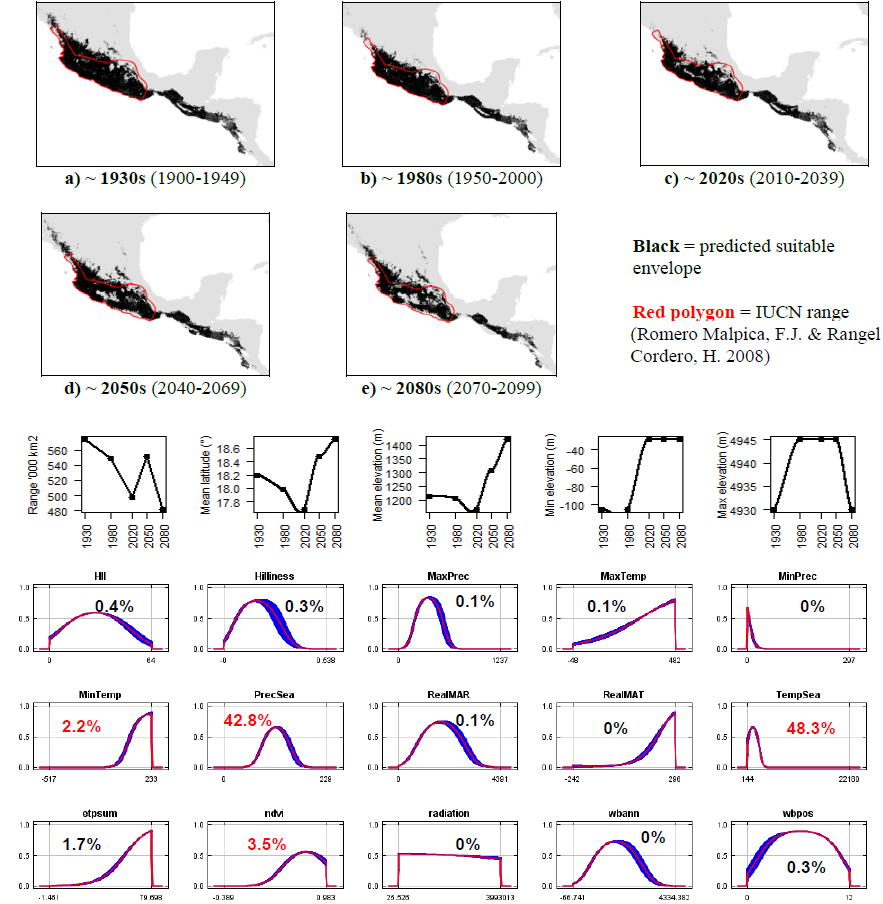

**#79 - Dice’s cottontail** (*Sylvilagus dicei*) *n* = 8

**Expert:** Jan Schipper, Arizona State University

**Expert evaluation:** Poor

**Data:** Only modern

**Envelope:** Climatic and habitat

**Dispersal distance:** 0.01km/year (Similar ecology to *S.cognatus*)

**Status:** UNMODELLABLE; **Included in final analysis:** X

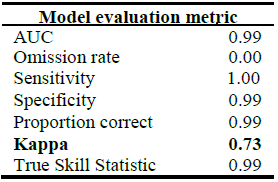

| Model evaluation metric |  |
| --- | --- |
| AUC | 0.99 |
| Omission rate | 0.00 |
| Sensitivity | 1.00 |
| Specificity | 0.99 |
| Proportion correct | 0.99 |
| **Kappa** | **0.73** |
| True Skill Statistic | 0.99 |

**Summary:** The Dice’s cottontail’s bioclimatic envelope is predicted to decrease by 50% with a ~1° mean latitudinal shift towards the Equator and a mean decrease in elevation of ~50m driven by a decrease in maximum elevation. 95% of the permutation importance of the model was contributed to by temperature seasonality (93.7%) and annual water balance (2.6%).

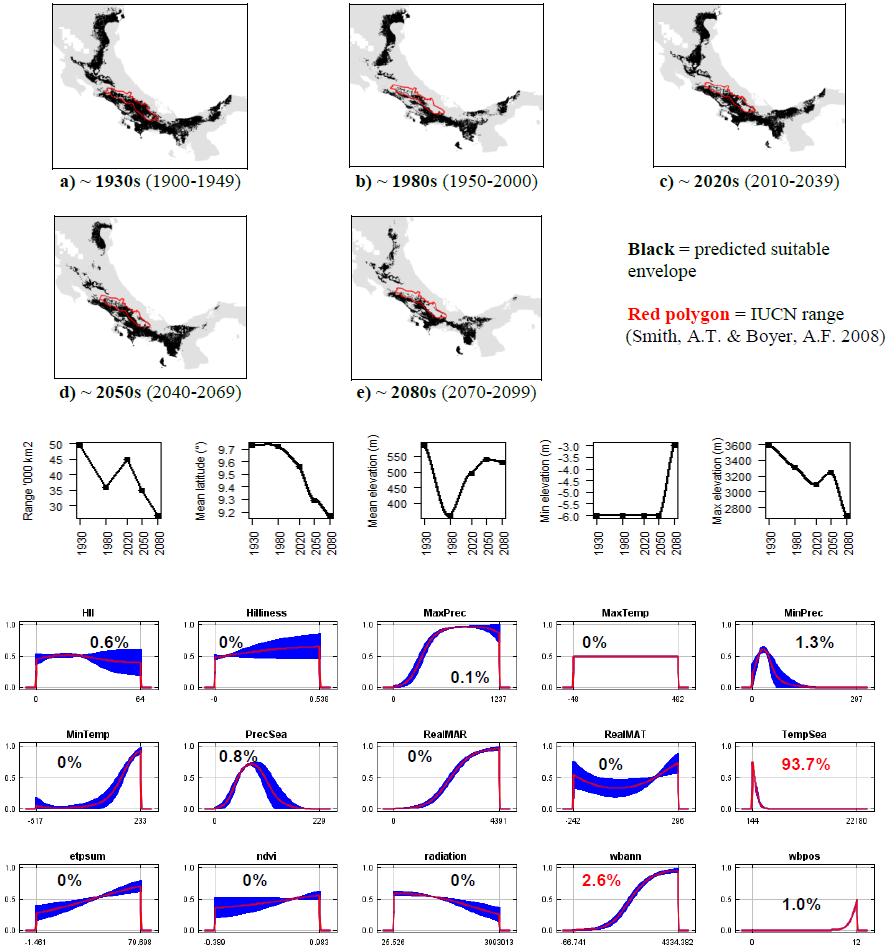

**#80 - Eastern cottontail** (*Sylvilagus floridanus*) *n* = 1104

**Expert:** Jorge Vazquez, Laboratorio de Ecologia del Comportamiento, UAT-UNAM

**Expert evaluation:** Medium

**Data:** Modern and historic

**Envelope:** Climatic and habitat

**Dispersal distance:** 7.5km/year (Similar ecology to *S.palustris*)

**Status:** MODELLABLE; **Included in final analysis:** √

| Model evaluation metric |  |
| --- | --- |
| AUC | 0.92 |
| Omission rate | 0.09 |
| Sensitivity | 0.91 |
| Specificity | 0.93 |
| Proportion correct | 0.93 |
| **Kappa** | **0.69** |
| True Skill Statistic | 0.84 |

**Summary:** The Eastern cottontail’s bioclimatic envelope is predicted to increase by 20% with a ~2° mean latitudinal polewards shift and a mean increase in elevation of ~90m driven by an increase in minimum and maximum elevation. 95% of the permutation importance of the model was contributed to by annual evapotranspiration (29.5%), precipitation seasonality (26.2%), minimum temperature (11.5%), temperature seasonality (8.5%), minimum precipitation (8.1%), maximum temperature (3.3%), normalised difference vegetation index (2.9%), maximum precipitation (2.8%) and annual water balance (2.3%).

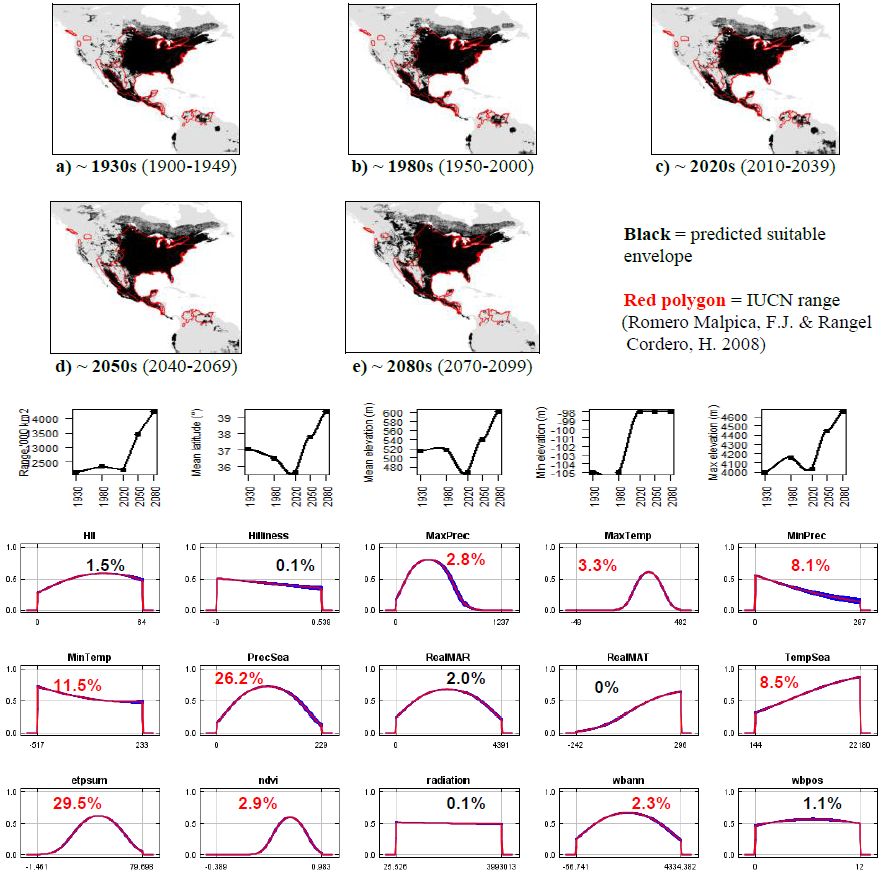

**#81 - Tres Marias cottontail** (*Sylvilagus graysoni*) *n* = 6

**Expert:** Consuelo Lorenzo, Departamento Conservation de la Biodiversidad, Chiapas

**Expert evaluation:** Good

**Data:** Modern and historic

**Envelope:** Climatic and habitat

**Dispersal distance:** 0.01km/year (Average for island species)

**Status:** MODELLABLE; **Included in final analysis:** √

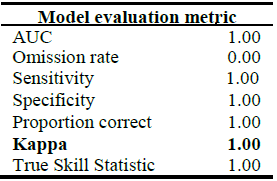

**Summary:** The Tres Marias cottontail’s bioclimatic envelope is predicted to decrease by 20% with a no latitudinal polewards shift and a mean increase in elevation of ~25m driven by an increase in minimum elevation. 95% of the permutation importance of the model was contributed to by human influence index (56.0%), minimum precipitation (32.4%), precipitation seasonality (5.2%) and annual evapotranspiration (3.2%).

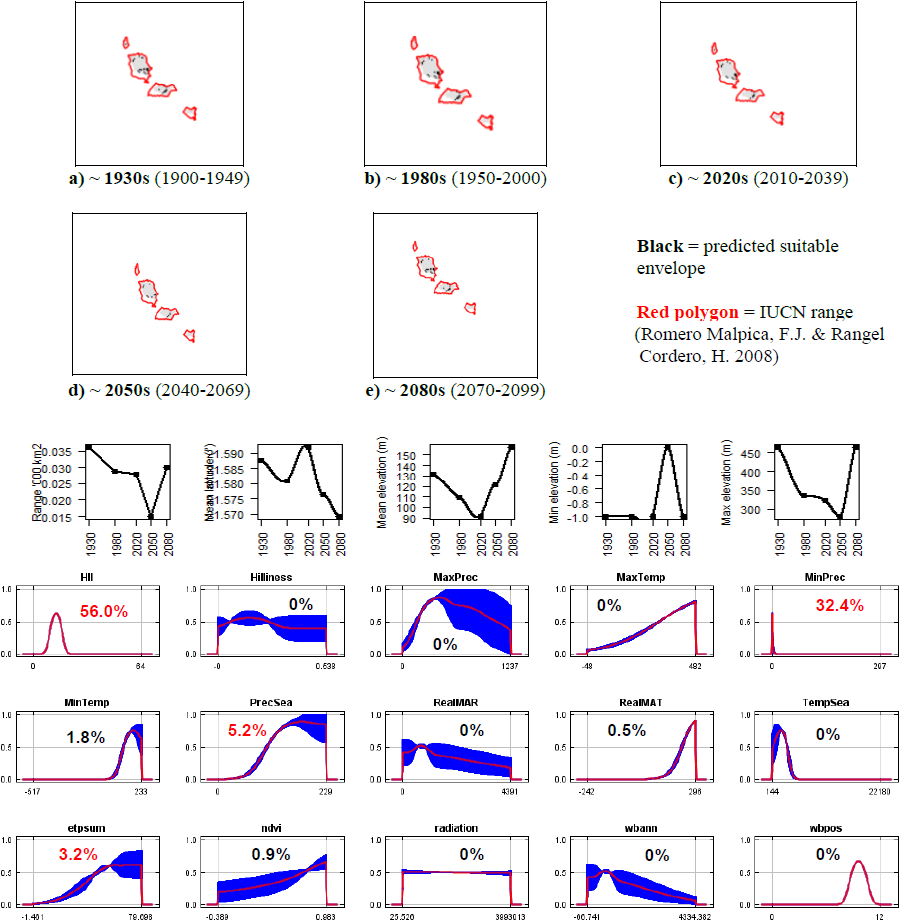

**#82 - Omilteme cottontail** (*Sylvilagus insonus*)

*n* = 3

**Expert:** Alejandro Velazquez, UNAM-Canada

**Expert evaluation:** Good

**Data:** Only modern

**Envelope:** Climatic and habitat

**Dispersal distance:** 0.01km/year (Similar ecology to *S.dicei*)

**Status:** MODELLABLE; **Included in final analysis:** √

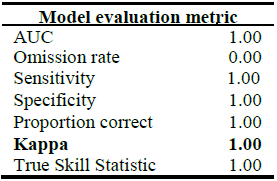

**Summary:** The Omilteme cottontail’s bioclimatic envelope is predicted to decrease by 80% with a no latitudinal polewards shift and a mean increase in elevation of ~120m driven by an increase in maximum elevation. 95% of the permutation importance of the model was contributed to by normalised difference vegetation index (78.5%), precipitation seasonality (11.3%) and temperature seasonality (5.3%).

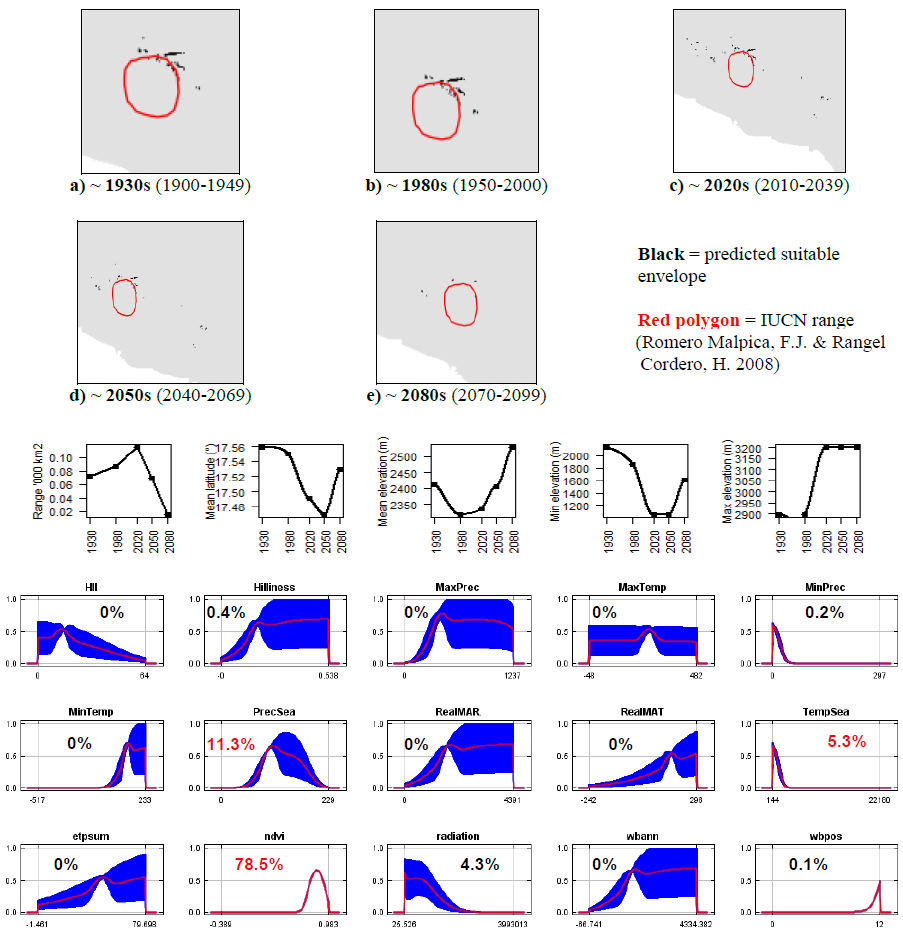

**#83 - San Jose brush rabbit** (*Sylvilagus mansuetus*) *n* = 9

**Expert:** Tamara Rioja Pardela, Universidad de Ciencias y

Artes de Chiapas, Mexico

**Expert evaluation:** Good

**Data:** Only modern

**Envelope:** Climatic and habitat

**Dispersal distance:** 0.01km/year (Average for island species)

**Status:** MODELLABLE; **Included in final analysis:** √

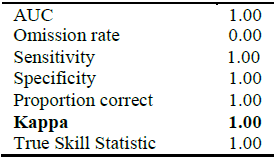

**Summary:** The San Jose brush rabbit’s bioclimatic envelope is predicted to decrease by 25% with a no latitudinal polewards shift and a mean increase in elevation of ~30m driven by an increase in minimum elevation. 95% of the permutation importance of the model was contributed to by temperature seasonality (36.7%), human influence index (34.0%), minimum precipitation (21.8%) and precipitation seasonality (3.2%).

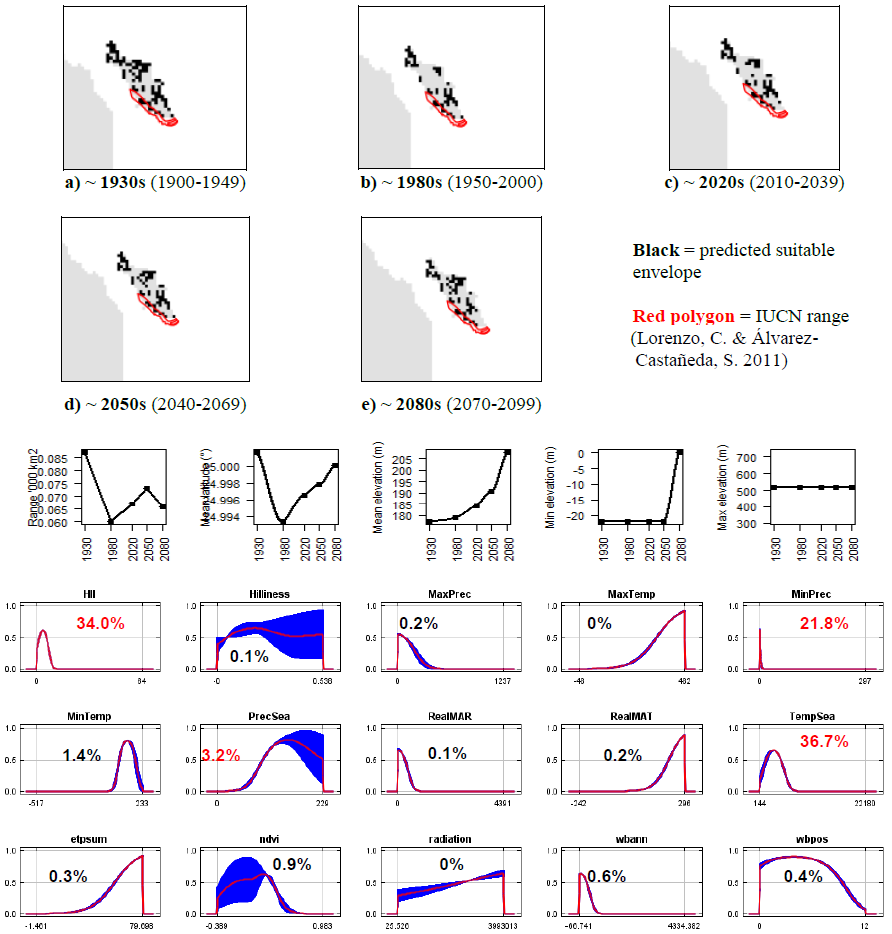

**#84 - Mountain cottontail** (*Sylvilagus nuttallii*) *n* = 290

**Expert:** Jennifer Frey, New Mexico State University

**Expert evaluation:** Medium

**Data:** Modern and historic

**Envelope:** Climatic and habitat

**Dispersal distance:** 7.5km/year (Similar ecology to *S.palustris*)

**Status:** MODELLABLE; **Included in final analysis:** √

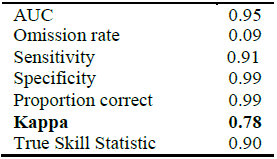

|  |  |
| --- | --- |
| AUC | 0.95 |
| Omission rate | 0.09 |
| Sensitivity | 0.91 |
| Specificity | 0.99 |
| Proportion correct | 0.99 |
| **Kappa** | **0.78** |
| True Skill Statistic | 0.90 |

**Summary:** The Mountain cottontail’s bioclimatic envelope is predicted to decrease by 20% with a ~1° mean latitudinal polewards shift and a mean increase in elevation of ~40m driven by an increase in minimum elevation. 95% of the permutation importance of the model was contributed to by mean annual temperature (64.0%), maximum temperature (26.9%), temperature seasonality (2.8%) and minimum precipitation (1.4%).

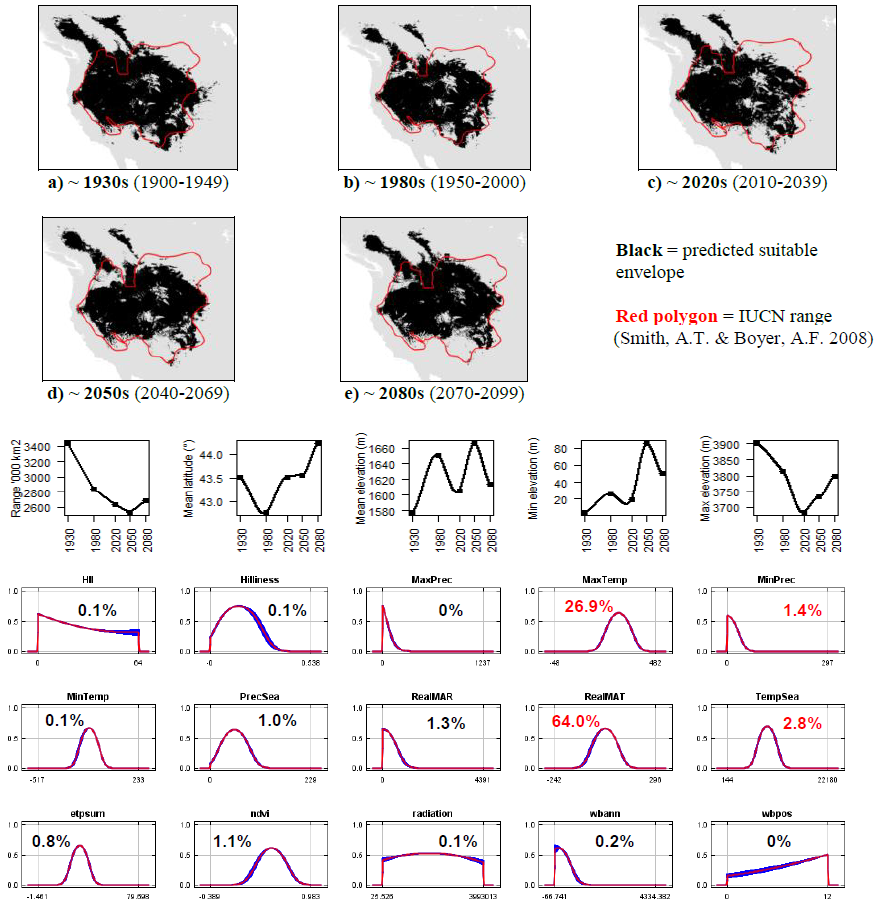

**#85 - Appalachian cottontail** (*Sylvilagus obscurus*) *n* = 39

**Expert:** Michael Barbour, Alabama Natural Heritage Program

**Expert evaluation:** Medium

**Data:** Modern and historic

**Envelope:** Climatic only

**Dispersal distance:** 0.01km/year (Similar ecology to *S.dicei*)

**Status:** MODELLABLE; **Included in final analysis:** √

|  |  |
| --- | --- |
| AUC | 0.97 |
| Omission rate | 0.05 |
| Sensitivity | 0.95 |
| Specificity | 0.99 |
| Proportion correct | 0.99 |
| **Kappa** | **0.73** |
| True Skill Statistic | 0.95 |

**Summary:** The Appalachian cottontail’s bioclimatic envelope is predicted to decrease by 10% with a ~3° mean latitudinal polewards shift and a mean increase in elevation of ~70m. 95% of the permutation importance of the model was contributed to by annual evapotranspiration (42.0%), minimum temperature (24.2%), minimum precipitation (18.8%) and precipitation seasonality (10.0%).

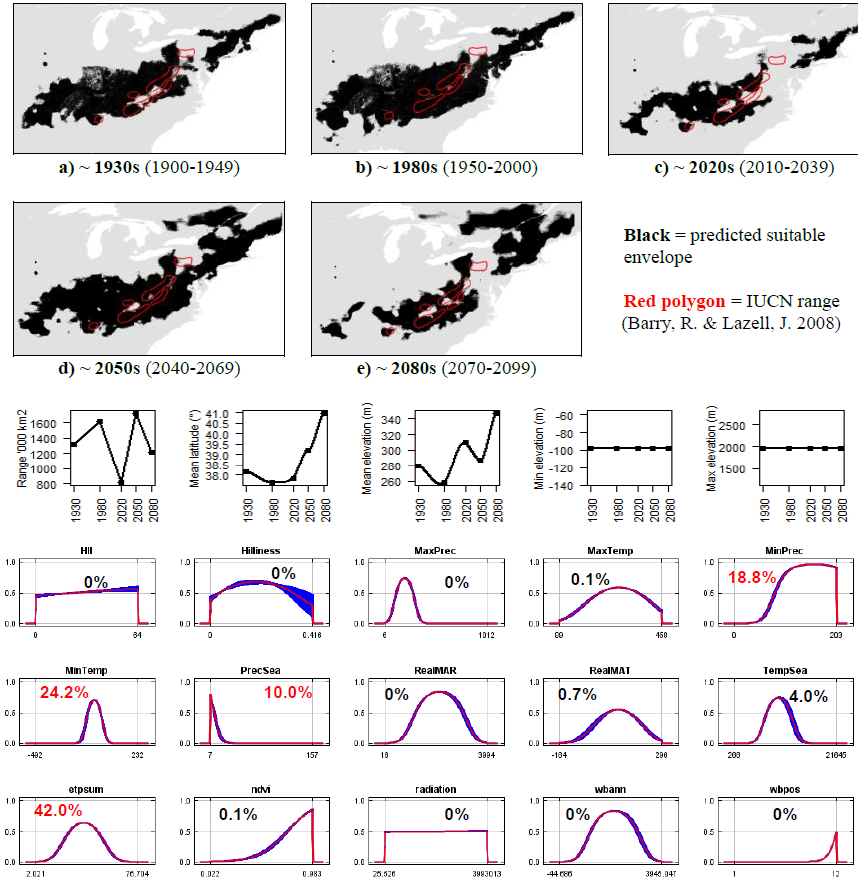

**#86 - Marsh rabbit** (*Sylvilagus palustris*) *n* = 25

**Expert:** Bob McCleery, University of Florida

**Expert evaluation:** Good

**Data:** Only modern

**Envelope:** Climatic and habitat

**Dispersal distance:** 7.5km/year (Expert)

**Status:** MODELLABLE; **Included in final analysis:** √

|  |  |
| --- | --- |
| AUC | 0.99 |
| Omission rate | 0.00 |
| Sensitivity | 1.00 |
| Specificity | 0.99 |
| Proportion correct | 0.99 |
| **Kappa** | **0.75** |
| True Skill Statistic | 0.99 |

**Summary:** The Marsh rabbit’s bioclimatic envelope is predicted to increase by 90% with a ~1° mean latitudinal polewards shift and a mean increase in elevation of ~2m driven by an increase in maximum elevation. 95% of the permutation importance of the model was contributed to by surface roughness index (73.1%), minimum precipitation (18.3%) and precipitation seasonality (4.9%).

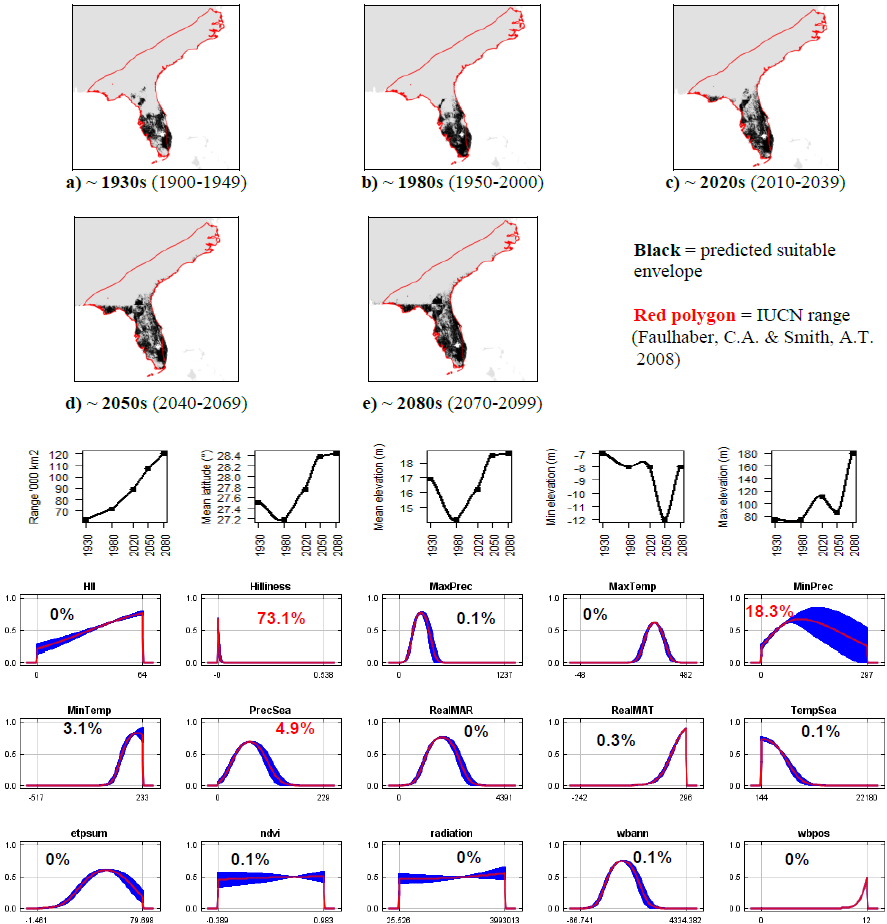

**#87 - Robust cottontail** (*Sylvilagus robustus*) *n* = 9

**Expert:** Dana Lee, Oklahoma State University

**Expert evaluation:** Poor

**Data:** Modern and historic

**Envelope:** Climatic and habitat

**Dispersal distance:** 0.01km/year (Similar ecology to *S.dicei*)

**Status:** UNMODELLABLE; **Included in final analysis:** X

|  |  |
| --- | --- |
| AUC | 0.94 |
| Omission rate | 0.11 |
| Sensitivity | 0.89 |
| Specificity | 0.99 |
| Proportion correct | 0.99 |
| **Kappa** | **0.27** |
| True Skill Statistic | 0.88 |

**Summary:** The Robust cottontail’s bioclimatic envelope is predicted to decrease by 90% with a ~4° mean latitudinal polewards shift and a mean increase in elevation of ~480m driven by an increase in minimum elevation. 95% of the permutation importance of the model was contributed to by precipitation seasonality (68.5%), minimum precipitation (8.2%), temperature seasonality (7.1%), minimum temperature (4.5%), mean annual precipitation (3.6%), annual water balance (2.6%) and number of months with a positive water balance (1.6%).

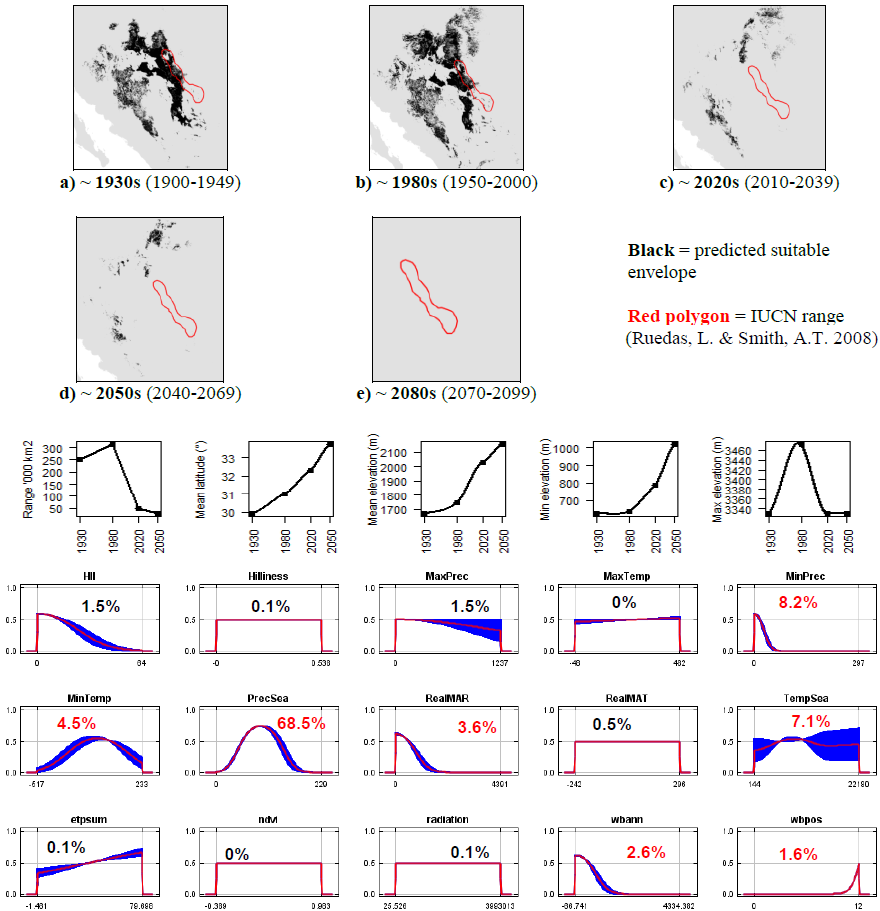

**#88 - New England cottontail** (*Sylvilagus transitionalis*) *n* = 18

**Expert:** John Litvaitis, University of New Hampshire

**Expert evaluation:** Medium

**Data:** Modern and historic

**Envelope:** Climatic and habitat

**Dispersal distance:** 3km/year (Expert)

**Status:** MODELLABLE; **Included in final analysis:** √

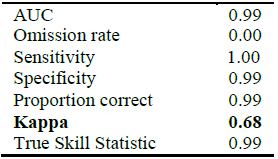

**Summary:** The New England cottontail’s bioclimatic envelope is predicted to increase by 110% with a ~1° mean latitudinal polewards shift and a mean increase in elevation of ~70m driven by an increase in maximum elevation. 95% of the permutation importance of the model was contributed to by annual evapotranspiration (23.5%), temperature seasonality (22.2%), minimum temperature (17.4%), normalised difference vegetation index (16.0%), mean annual temperature (8.6%), maximum temperature (5.9%) and minimum precipitation (5.9%).

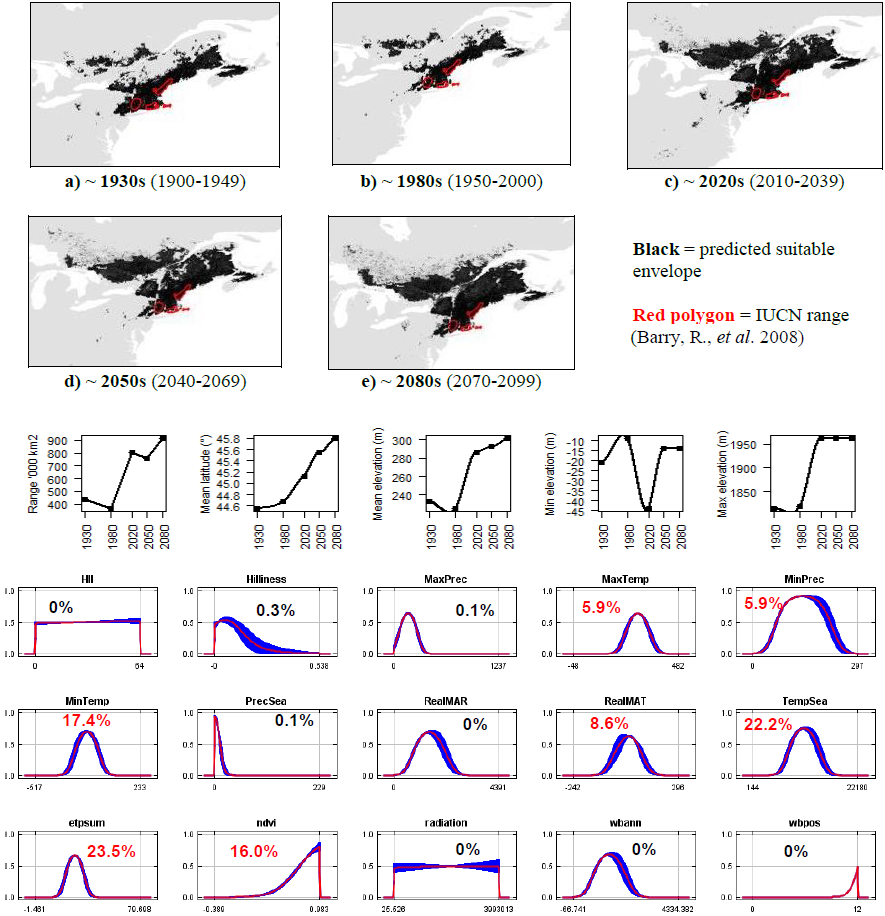

**#89 - Venezuelan lowland rabbit** (*Sylvilagus varynaensis*) *n* = 6

**Expert:** Daniel Lew, Venezuelan Institute of Scientific Research, Ecology Centre, Biodiversity Unit

**Expert evaluation:** Poor

**Data:** Only modern

**Envelope:** Climatic and habitat

**Dispersal distance:** 3km/year (Similar ecology to *S.transitionalis*)

**Status:** UNMODELLABLE; **Included in final analysis:** X

|  |  |
| --- | --- |
| AUC | 0.99 |
| Omission rate | 0.00 |
| Sensitivity | 1.00 |
| Specificity | 0.99 |
| Proportion correct | 0.99 |
| **Kappa** | **0.92** |
| True Skill Statistic | 0.99 |

**Summary:** The Venezuelan lowland rabbit’s bioclimatic envelope is predicted to decrease by 100% with a ~1.5° mean latitudinal polewards shift and a mean increase in elevation of ~275m driven by an increase in minimum elevation. 95% of the permutation importance of the model was contributed to by temperature seasonality (97.7%).

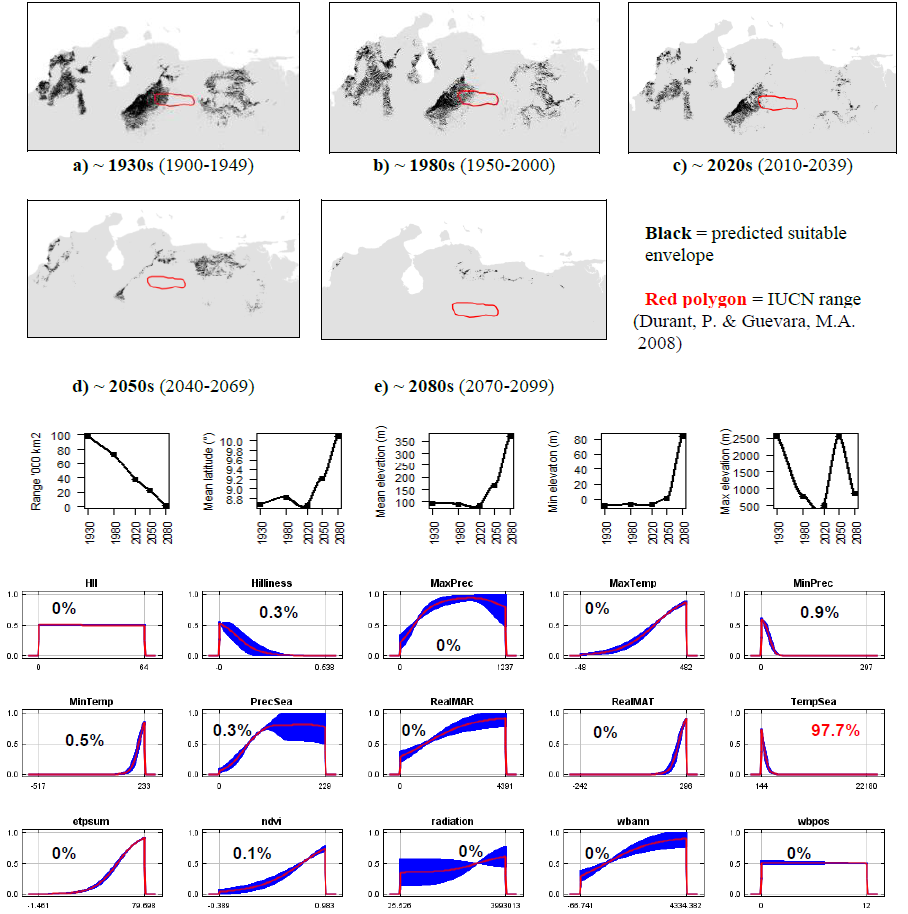

---

**Table S1.**
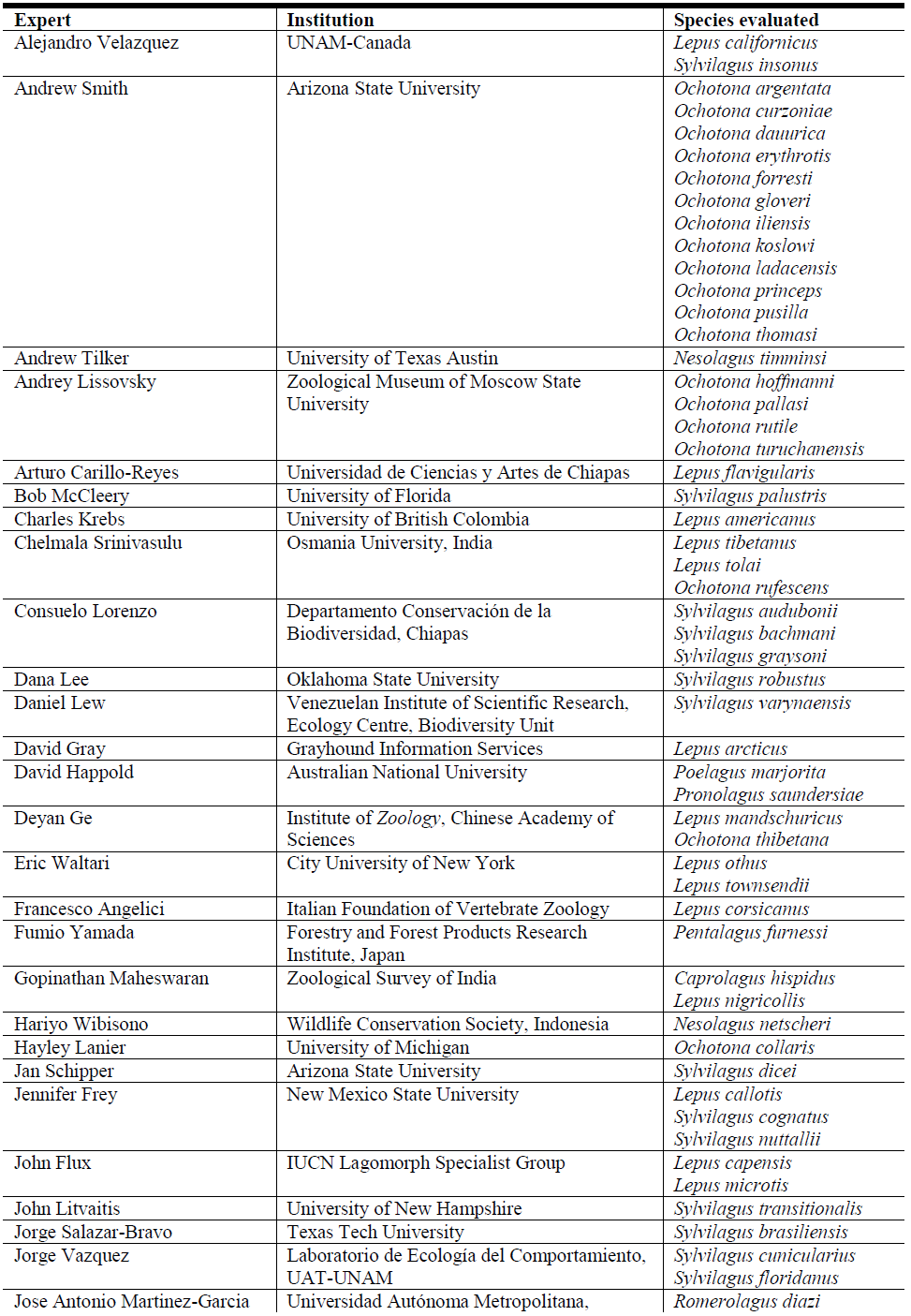
Lagomorph experts, institutions and species evaluated.

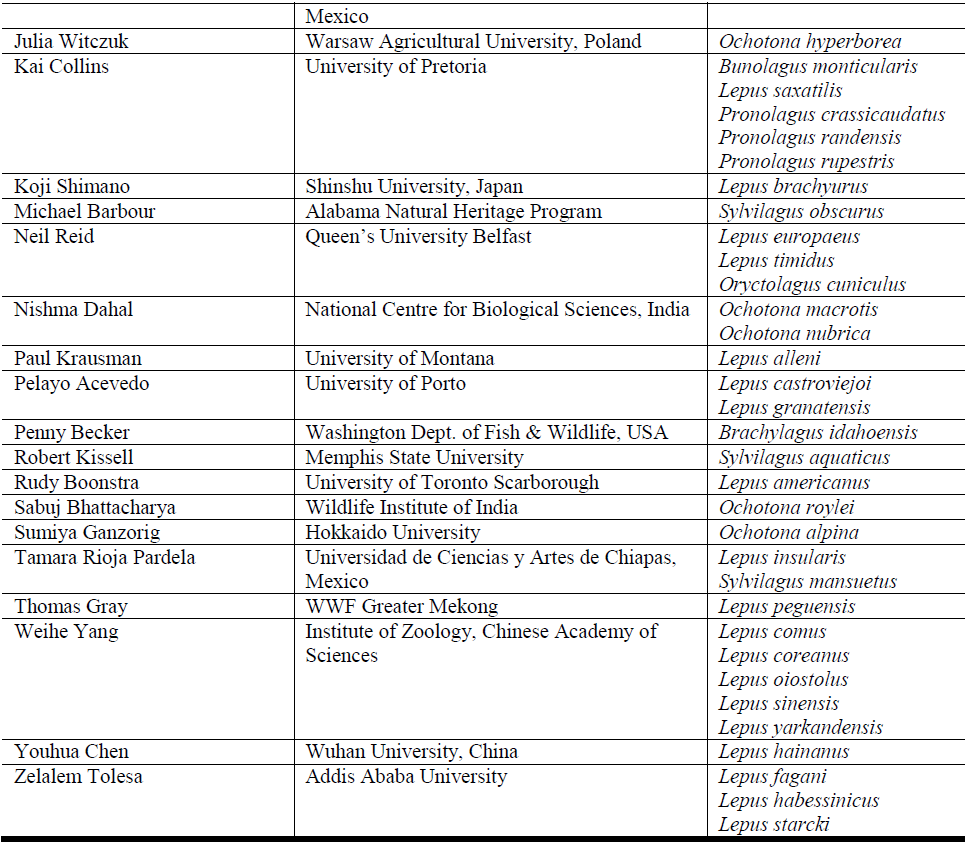

**Figure S1.**
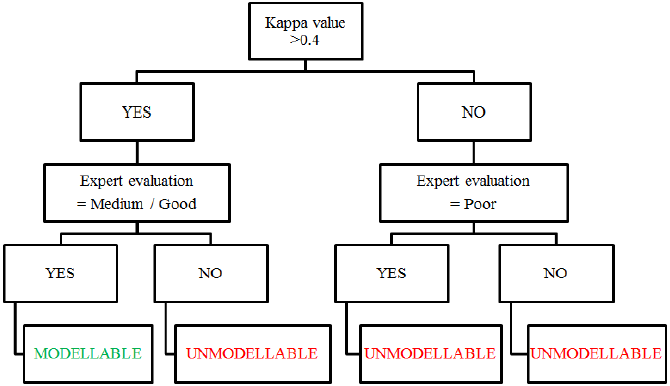
Framework for assessing whether species were “modellable” or “unmodellable” based on Kappa values and expert evaluation classification. The optimum threshold for Kappa was taken as 0.4 [9], [10], [11]. Expert evaluations were classified according to Anderson *et al*. [12].

**Table S2.** Results of Generalised Least Square models characterising predicted lagomorph bioclimatic envelope changes. Significant p values are in bold. Group refers to lagomorph taxonomy, i.e. pikas, rabbits and hares & jackrabbits.

| Response variable | Term | *F* | *Df* | *p* |
| --- | --- | --- | --- | --- |
| Range change (km)<br>$F_{df=1,234}=0.586, p=0.445$ | Group | 0.004 | 2, 229 | 0.950 |
|  | Year | 3.710 | 1, 228 | **0.026** |
|  | Group: Year | 0.537 | 2, 226 | 0.585 |
| Mean latitudinal change (°)<br>$F_{df=1,234}=13.460, p<0.001$ | Group | 12.798 | 2, 229 | **<0.001** |
|  | Year | 1.603 | 1, 228 | 0.204 |
|  | Group: Year | 0.448 | 2, 226 | 0.640 |
| Mean elevation change (m)<br>$F_{df=1,234}=44.184, p<0.0001$ | Group | 19.458 | 2, 229 | **<0.001** |
|  | Year | 21.140 | 1, 228 | **<0.001** |
|  | Group: Year | 3.541 | 2, 226 | **0.031** |

**Table S3.**
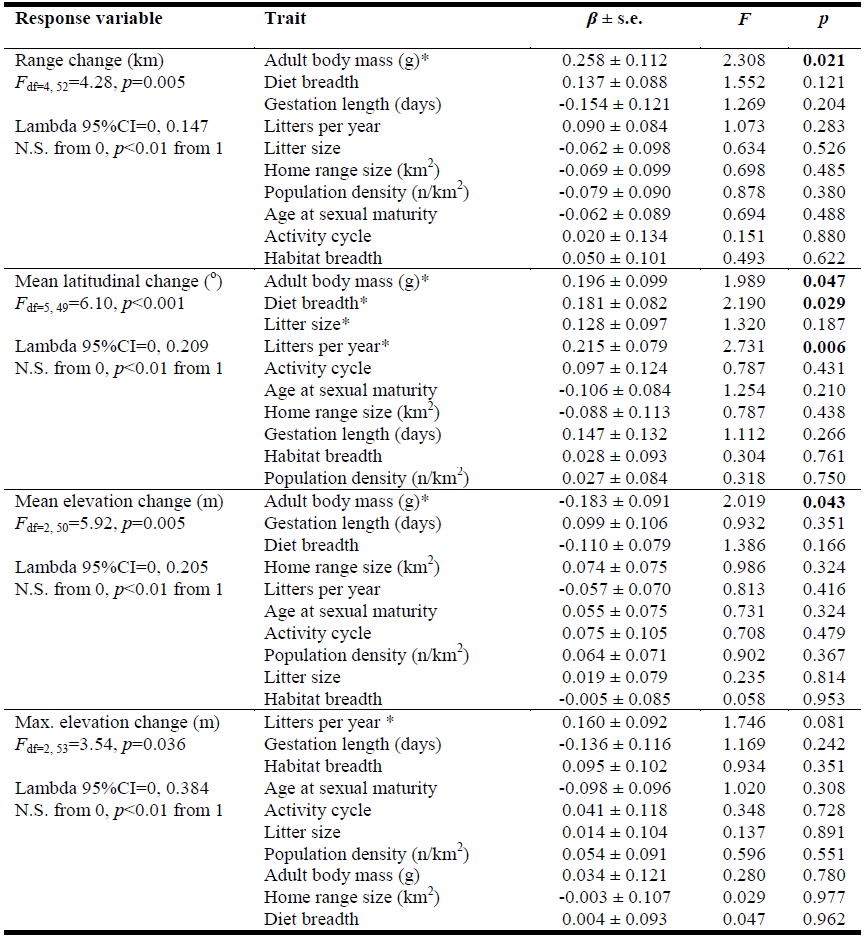
Results of phylogenetically-controlled generalised least square regressions. Significant*p* values for model-averaged coefficients are in bold. *F* and *p* values for the top model are listed under each response variables; asterisks (*) indicate traits in the top model. Lambda (X) confidence intervals and significance from 0 and 1 are also shown.

| Response variable | Trait | $\beta \pm \text{s.e.}$ | $F$ | $p$ |
| --- | --- | --- | --- | --- |
| Range change (km)<br>$F_{df=4, 52}=4.28, p=0.005$<br>Lambda 95%CI=0, 0.147<br>N.S. from 0, $p<0.01$ from 1 | Adult body mass (g)* | $0.258 \pm 0.112$ | 2.308 | **0.021** |
| | Diet breadth | $0.137 \pm 0.088$ | 1.552 | 0.121 |
| | Gestation length (days) | $-0.154 \pm 0.121$ | 1.269 | 0.204 |
| | Litters per year | $0.090 \pm 0.084$ | 1.073 | 0.283 |
| | Litter size | $-0.062 \pm 0.098$ | 0.634 | 0.526 |
| | Home range size (km<sup>2</sup>) | $-0.069 \pm 0.099$ | 0.698 | 0.485 |
| | Population density (n/km<sup>2</sup>) | $-0.079 \pm 0.090$ | 0.878 | 0.380 |
| | Age at sexual maturity | $-0.062 \pm 0.089$ | 0.694 | 0.488 |
| | Activity cycle | $0.020 \pm 0.134$ | 0.151 | 0.880 |
| | Habitat breadth | $0.050 \pm 0.101$ | 0.493 | 0.622 |
| Mean latitudinal change (°)<br>$F_{df=5, 49}=6.10, p<0.001$<br>Lambda 95%CI=0, 0.209<br>N.S. from 0, $p<0.01$ from 1 | Adult body mass (g)* | $0.196 \pm 0.099$ | 1.989 | **0.047** |
| | Diet breadth* | $0.181 \pm 0.082$ | 2.190 | **0.029** |
| | Litter size* | $0.128 \pm 0.097$ | 1.320 | 0.187 |
| | Litters per year* | $0.215 \pm 0.079$ | 2.731 | **0.006** |
| | Activity cycle | $0.097 \pm 0.124$ | 0.787 | 0.431 |
| | Age at sexual maturity | $-0.106 \pm 0.084$ | 1.254 | 0.210 |
| | Home range size (km<sup>2</sup>) | $-0.088 \pm 0.113$ | 0.787 | 0.438 |
| | Gestation length (days) | $0.147 \pm 0.132$ | 1.112 | 0.266 |
| | Habitat breadth | $0.028 \pm 0.093$ | 0.304 | 0.761 |
| | Population density (n/km<sup>2</sup>) | $0.027 \pm 0.084$ | 0.318 | 0.750 |
| Mean elevation change (m)<br>$F_{df=2, 50}=5.92, p=0.005$<br>Lambda 95%CI=0, 0.205<br>N.S. from 0, $p<0.01$ from 1 | Adult body mass (g)* | $-0.183 \pm 0.091$ | 2.019 | **0.043** |
| | Gestation length (days) | $0.099 \pm 0.106$ | 0.932 | 0.351 |
| | Diet breadth | $-0.110 \pm 0.079$ | 1.386 | 0.166 |
| | Home range size (km<sup>2</sup>) | $0.074 \pm 0.075$ | 0.986 | 0.324 |
| | Litters per year | $-0.057 \pm 0.070$ | 0.813 | 0.416 |
| | Age at sexual maturity | $0.055 \pm 0.075$ | 0.731 | 0.324 |
| | Activity cycle | $0.075 \pm 0.105$ | 0.708 | 0.479 |
| | Population density (n/km<sup>2</sup>) | $0.064 \pm 0.071$ | 0.902 | 0.367 |
| | Litter size | $0.019 \pm 0.079$ | 0.235 | 0.814 |
| | Habitat breadth | $-0.005 \pm 0.085$ | 0.058 | 0.953 |
| Max. elevation change (m)<br>$F_{df=2, 53}=3.54, p=0.036$<br>Lambda 95%CI=0, 0.384<br>N.S. from 0, $p<0.01$ from 1 | Litters per year * | $0.160 \pm 0.092$ | 1.746 | 0.081 |
| | Gestation length (days) | $-0.136 \pm 0.116$ | 1.169 | 0.242 |
| | Habitat breadth | $0.095 \pm 0.102$ | 0.934 | 0.351 |
| | Age at sexual maturity | $-0.098 \pm 0.096$ | 1.020 | 0.308 |
| | Activity cycle | $0.041 \pm 0.118$ | 0.348 | 0.728 |
| | Litter size | $0.014 \pm 0.104$ | 0.137 | 0.891 |
| | Population density (n/km<sup>2</sup>) | $0.054 \pm 0.091$ | 0.596 | 0.551 |
| | Adult body mass (g) | $0.034 \pm 0.121$ | 0.280 | 0.780 |
| | Home range size (km<sup>2</sup>) | $-0.003 \pm 0.107$ | 0.029 | 0.977 |
| | Diet breadth | $0.004 \pm 0.093$ | 0.047 | 0.962 |

**Table S4.** Results of Spearman’s rank correlation.

| **Data** | **Method** | **Estimated measure of association** | ***S*** | ***p*** |
| --- | --- | --- | --- | --- |
| Adult body mass (g) vs.<br>Dispersal distance (km/year) | Rho | 0.077 | 30002.15 | 0.565 |

